# An evolutionary-based approach to quantify the genetic barrier to drug resistance in fast-evolving viruses: an application to HIV-1 subtypes and integrase inhibitors

**DOI:** 10.1101/647297

**Authors:** Kristof Theys, Pieter Libin, Kristel Van Laethem, Ana B Abecasis

## Abstract

Viral pathogens causing global disease burdens are often characterised by high rates of evolutionary changes, facilitating escape from therapeutic or immune selective pressure. Extensive viral diversity at baseline can shorten the time to resistance emergence and alter mutational pathways, but the impact of genotypic background on the genetic barrier can be difficult to capture, in particular for antivirals in experimental stages, recently approved or expanded into new settings. We developed an evolutionary-based counting method to quantify the population genetic potential to resistance and assess differences between populations. We demonstrate its applicability to HIV-1 integrase inhibitors, as their increasing use globally contrasts with limited availability of non-B subtype resistant sequences and corresponding knowledge gap on drug resistance. A large sequence dataset encompassing most prevailing subtypes and resistance mutations of first- and second-generation inhibitors were investigated. A varying genetic potential for resistance across HIV-1 subtypes was detected for 15 mutations at 12 positions, with notably 140S in subtype B, while 140C was discarded to vary across subtypes. An additional analysis for HIV-1 reverse transcriptase inhibitors identified a higher potential for 65R in subtype C, on the basis of a differential codon usage not reported before. The evolutionary interpretation of genomic differences for antiviral treatment remains challenging. Our framework advances existing counting methods with an increased sensitivity that identified novel subtype dependencies as well as rejected previous statements. Future applications include novel HIV-1 drug classes as well as other viral pathogens.

## 1 Introduction

The advent of antiviral treatment resulted globally in significant health gains and averted deaths by preventing viral infections and improving disease outcomes. Major human pathogens targeted by antivirals are fast-evolving, genetically diverse viruses [13, 19, 20], allowing for rapid adaptation through the emergence of resistance-conferring mutations. A key concept in understanding the dynamics of resistance development is the genetic barrier, which ultimately quantifies the evolutionary time to viral escape from drug selective pressure [4, 14, 16, 39]. The virus make-up at baseline can shorten the evolutionary distance to antiviral resistance and, together with treatment- and patient-related factors, alter the mode and tempo of resistance emergence [20, 24]. The imprint of genotypic background on viral escape dynamics can be however difficult to capture. In particular for newer antivirals when resistance knowledge is initially limited to *in vitro* selection experiments informed on genetically-limited backbones and inferred from observations in well-controlled clinical studies or patient cohorts underrepresenting the full spectrum of viral diversity.

A notable example of success is the evolution of the management of Human Immunodeficiency Virus type-1 (HIV-1) infection in the last three decades, with antivirals available from multiple drug classes that drastically reduced morbidity and mortality related to HIV-1. The recent class of integrase strand transfer inhibitors (INSTIs), directed against the integrase enzyme by blocking the strand transfer step of viral DNA integration, has considerably expanded treatment options and reduced the probability of virological failure, predominantly in resource-rich settings where INSTI use is widespread. To date, first-generation INSTIs raltegravir (RTG) and elvitegravir (EVG) and second-generation INSTIs dolutegravir (DTG) and bictegravir (BIC) are approved for HIV-1 treatment and a preferred option for first-line regimens [27, 36]. INSTIs are anticipated to become also widespread in low- and middle-income countries (LMICs), and the use of INSTIs has been shown to be cost-effective [30], although rates of acquired drug resistance are increasing in these settings. An impact of HIV-1 genetic diversity, classified into groups and subtypes, on resistance development has been well documented for the historical classes of protease and reverse transcriptase (RT) inhibitors, mainly resulting from preferential codon usage across subtypes [1, 40, 42]. The evolutionary mechanisms underlying viral escape from INSTI selective pressure, despite the identification of mutational pathways, are still being unfold in particular for non-B subtypes which are prevalent in LMICs [12]. A low observed complexity of INSTI resistance profiles has consequences for the relevance of HIV-1 diversity for the genetic barrier to resistance [6, 17].

Extensive subtype mappings of integrase diversity in treatment-naive patients revealed amino acid polymorphisms at resistance-associated positions [20, 25, 35, 45]. Apart from a documented subtype B INSTI resistance pathway attributed to differential codon usage at position 140 [17, 23, 32, 38], the role of baseline nucleotide variation on INSTI resistance development has been less systematically investigated. The probabilistic models of resistance evolution that previously quantified the genetic barrier for the historical drug classes require sequence data of various subtypes from treatment-experienced patients [4, 14, 39], which are limited to date in the context of (second-generation) INSTIs [34, 35]. Alternatively, the genetic barrier can be estimated by the number and type of required nucleotide substitutions to evolve from wild-type virus to a resistance mutation [43]. While this modality was applied to HIV-1 integrase before [23, 32], these studies lacked resistance mutations of second-generation INSTIs, ignored subtype-specific effects or analyzed limited subtype distributions and relied on arbitrary substitution cost assignments.

We present a novel, optimized and evolutionary-based methodology to quantify the genetic potential to resistance in fast-evolving viral pathogens, facilitating a priori identification of an impact of varying genetic background on the emergence of resistance mutations. We demonstrate in this study our approach by the application to HIV-1 INSTIs. With the roll-out of INSTIs in LMICs, an increasing introduction of non-B subtypes in high-income countries [15, 18, 26, 28] and novel INSTIs anticipated, this study addressed the need for an in-depth understanding of the dynamics underlying INSTI resistance development across HIV-1 subtypes. We derived an optimized genetic barrier score based on empirical substitution costs, which contrasts with previous approaches that used arbitrary costs differing from *in vivo* estimates [46]. Furthermore, we advanced existing studies by considering a population-based estimate, where we take into account the most prevailing subtypes globally and calculate subtype-tailored summary scores. The framework can be easily applied to novel HIV-1 drug classes (e.g. attachment or maturation inhibitors) and extended to pathogens (e.g. Zika, Influenza, RSV, Herpes, Varicella, Dengue) which are currently targeted by antiviral drug development efforts.

## 2 Methods

### 2.1 HIV-1 dataset and drug resistance mutations

A dataset of HIV-1 integrase sequences from INSTI-naive patients was obtained from the Stanford HIV drug resistance database [35], aligned codon-correct using VIRULIGN [22] and classified by REGA subtyping tool v3 [3, 21, 31]. Only 1 sequence per patient and sequences without a stop codon were retained. Additionally, sequences with mutations 143C/H/R, 148H/K/R or 155H/S were considered a proxy for incorrect treatment status and excluded, given that transmission of INSTI resistance is infrequently reported to date. Integrase positions and mutations important for INSTI resistance were defined by an association with reduced susceptibility and virological response [6], or inclusion in HIV-1 drug resistance interpretation systems (Rega v10, HIVdb v8.7, ANRS v29) [5, 9, 44]. A total of 41 codon positions and 77 amino acid mutations implicated in viral escape from first- or second-generation INSTIs were investigated. In addition, **Supplementary Material** presents the same approach applied to quantify the evolutionary potential to resistance of the RT enzyme.

### 2.2 Codon diversity

We define the genetic barrier in terms of a particular sub-epidemic, e.g. with respect to all viruses that belong to a HIV-1 subtype in our study, and of a resistance mutation defined by an amino acid and a given position. An overview of the procedure to calculate the genetic barrier is given in Figure 1, using resistance mutation 140S as example. At every position, variability at the nucleotide level was assessed by the prevalence of nucleotide triplets (codon) in each sub-epidemic (Figure 1A). Codons with ambiguities consisting of *>*2 bases per nucleotide position or of two or more ambiguities per codon were not considered. A nucleotide ambiguity of exactly 2 bases was resolved in the two corresponding triplets, each counting for one half to their respective frequencies. All triplets with a prevalence above 1% in at least one sub-epidemic were retained, and triplets with a prevalence above 50% were defined as predominant for that sub-epidemic. The prevalence of resistance mutations and codon entropy were inferred from the triplet distribution.

**Figure 1.**
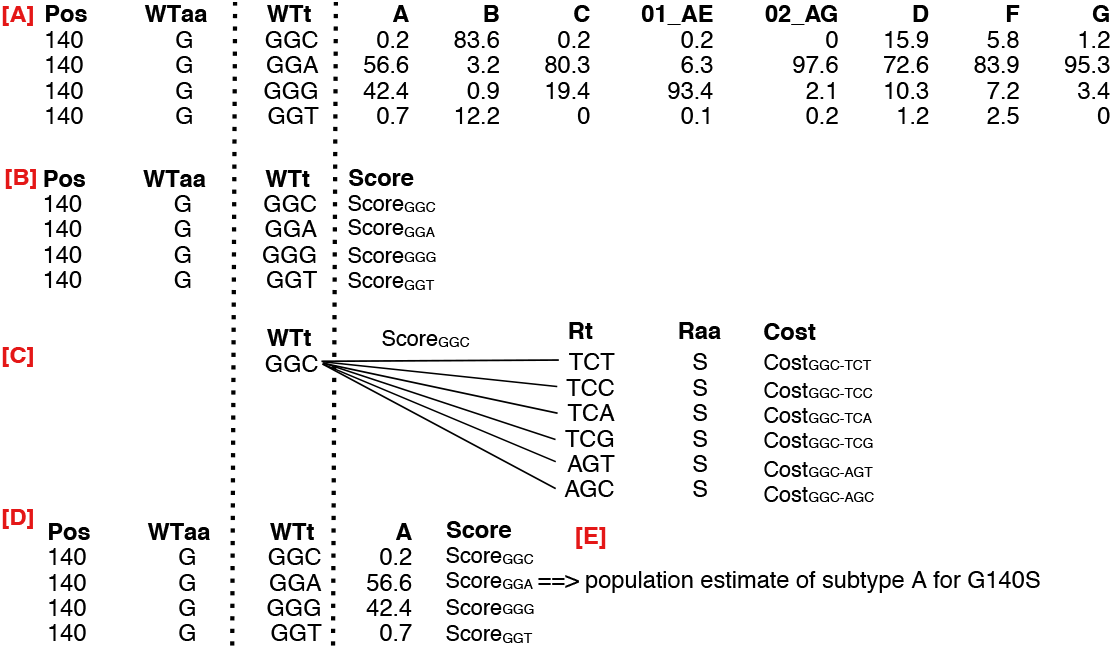
Overview of methodology for integrase resistance amino acid S (Raa) at position 140 (Pos). [A] Determine the distribution wild-type triplets (WTt) naturally present, the translated amino acid (WTaa) and their frequency across the 8 subtypes. [A] A score is assigned to each wild-type triplet to evolve into the resistance mutation S. [B] This score is obtained by iteratively determining the cumulative substitution cost for the change of WTt into each resistance triplet (Rt) translating into the amino acid S, and subsequently by the summation of the different cumulative costs. [D] Each score is iteratively weighted by the subtype frequency of the wild-type triplet, here shown for subtype A. [E] A population estimate of the genetic barrier is obtained for subtype A. This procedure is done for each subtype, resulting in a subtype-specific population genetic barrier for 140S. When all resistance-associated mutations are considered, a matrix of genetic barrier values is created where subtypes are shown as rows and mutations as columns. Figure 4 illustrates the visualisation of this matrix.

### 2.3 Substitution cost

A penalty score is assigned to the different types of nucleotide substitution through the transformation of empirically estimated substitution matrices into corresponding cost matrices. For each codon index *ci* ∈ {1, 2, 3}, the cost matrix 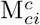 is given by the normalized and inverted substitution matrix 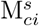 following normalisation:

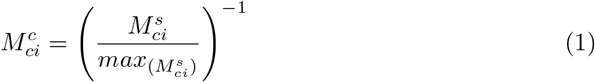

This transformation assigns the substitution with the highest rate (e.g. G → A) a cost of 1, and the costs of the other substitutions were proportionally adapted to this baseline cost. A cost of zero is assigned when no change occurs. For our study, a substitution matrix for each codon position was derived from HIV-1 integrase nucleotide sequences independently collected from the Los Alamos database, with codon-based substitution patterns and rates estimated under the General Time Reversible model (**Supplementary Material**).

### 2.4 Genetic barrier to resistance

Given a sub-epidemic *𝓔*, the genetic barrier to resistance mutation *R* is a cost function *GB*_*𝓔*_ (*R*) that quantifies the number and type of nucleotide substitutions required for the virus to evolve from wild type diversity in the virus population of *𝓔* to *R* (Figure 1). This baseline diversity is defined by the set of wild type type triplets {*W T*_*t,i*_}, and we will denote their prevalences as prev (*W T*_*t,i*_). Furthermore, as *R* is an amino acid, we enumerate all triplets that translate into *R* and will refer to this set as {*R*_*t,j*_}. First, we determine a score for a given wild type triplet *W T*_*t*_ to *R*, while considering all triplets that translate into *R* (i.e., {*R*_*t,j*_}):

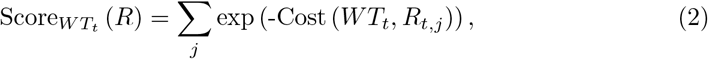

where Cost (.) quantifies the cumulative substitution cost to mutate from a *WT*_*t*_into a particular resistance triplet *R*_*t,a*_, as provided in Equation 3. This summation incorporates all possible evolution paths and subsequently assigns lower cumulative costs (i.e., more likely mutation pathways) a higher contribution to the total triplet score. Taking into account different but almost equally likely evolution paths, compared to only considering the minimum cumulative substitution cost, results in an elevated sensitivity to detect subtype-dependencies in the genetic barrier.

To compute the cumulative substitution cost for *R*_*t,a*_ (i.e., subst (*W T*_*t*_, *R*_*t,a*_)), we compute the sum of the substitution cost for each triplet position, using substitution matrices 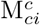 as defined in section 2.3:

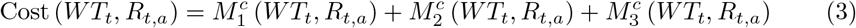

Finally, the genetic barrier *GB*_𝓔_ is defined as the sum of the scores (Equation 2) of the different wild type triplets in sub-epidemic 𝓔, weighted according to their prevalence:

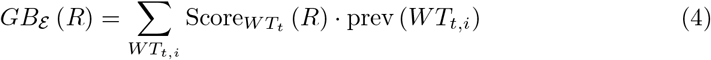

A *GB*_𝓔_ (*R*) is obtained for each resistance-associated mutation and subtype. In order to visualise most pronounced differences between sub-epidemics, we calculated the average distance in population genetic barrier for each sub-epidemic against other sub-epidemics.

### 2.5 Statistics

The non-parametric Mann-Whitney test was used to identify differences in the genetic barrier between HIV-1 subtypes, corrected for multiple hypothesis testing using the Benjamini-Hochberg method [43]. Each subtype was compared against a weighted sample (N=500) of the other subtypes, repeated one thousand times. In addition, pairwise comparisons of subtype B with each subtype separately were performed given that most knowledge on integrase resistance development is available for subtype B. Significant comparisons with a difference in the barrier score above 0.1 are retained. All analyses were done using the statistical software package R [33].

## 3 Results

### 3.1 Data

A total of 10235 viral sequences coding for the HIV-1 integrase enzyme fulfilled the inclusion criteria, resulting in 410 sequences classified as subtype A (4%), 5174 as subtype B (50.5%), 1837 as subtype C (17.9%), 1630 (15.9%) as CRF01 AE, 596 (5.8%) as CRF02 AG, 170 (1.7%) as subtype D, 257 (2.5%) as subtype F and 161 (1.6%) as subtype G. Nucleotide ambiguities only had a modest impact on data processing as the highest percentage of codons that did not fulfil the criteria was 7.6% for the highly polymorphic position 125 (**Supplementary Material**). The distributions of within-subtype pairwise diversity were unimodal and similar across subtypes **(Supplementary Material)**.

### 3.2 Natural variability of integrase

Variation in genetic barrier across subtypes requires a combination of differences in *WT*_*t*_ frequencies and in associated cost scores. We first illustrate relevant integrase genetic variability using triplet entropy calculations to elucidate variation within a single subtype. Next, identifying predominant triplets can reveal major variation between subtypes. In particular triplets associated with INSTI resistance are of interest since they result in a minimal cost.

Figure 2 shows within-subtype entropy values for each resistance position. Some of this within-subtype variation is translated into the presence of polymorphic mutations. Figure 3 shows the prevalence of resistance-associated mutations in the dataset **(Supplementary Material)**. Most prevalent mutations for subtype A were 50I (25.2%), 74I (22.4%), 203M (5.5%) and 97A (5.1%); for subtype B 156N (17.6%), 230N (10.4%), 50I (9.4%), 203M (6.4%), 119R (5.5%) and 151I (5.2%); for subtype C 50I (35.1%) and 74I (5.0%); for CRF01 AE 165I (17.2%); for CRF02 AG 74I (18.4%), 50I (10.6%), 74M (10.2%), 157Q (8.3%) and 97A (5.5%); for subtype D 203M (15.3%), 97A (6.2%) and 165I (6.2%); for subtype F 165I (30.2%), 50I (9.2%), 163R (6.2%) and 163K (5.4%); for subtype G 50I (15.1%), 74I (10.3%) and 165I (5.3%). These results illustrate both resistance-associated mutations prevalent across subtypes (e.g. 50I and 74I) as well as subtype-specific occurrences (e.g. 156N).

**Figure 2.**
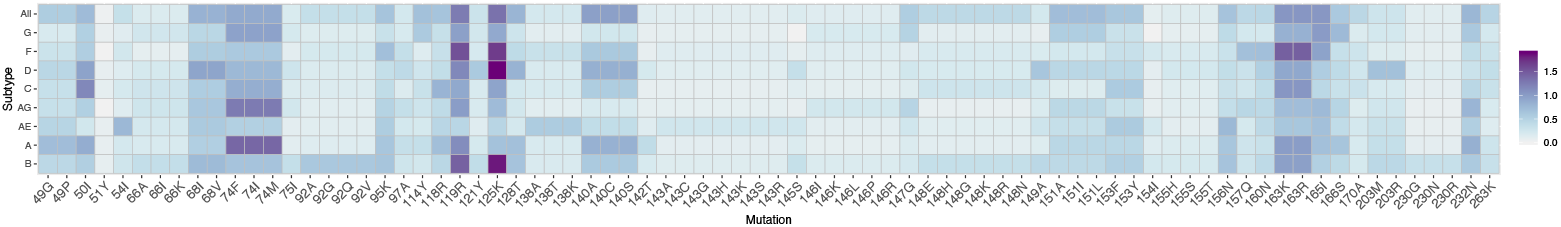
Triplet Entropy per position within each subtype, and also for all subtypes combined (All)

**Figure 3.**
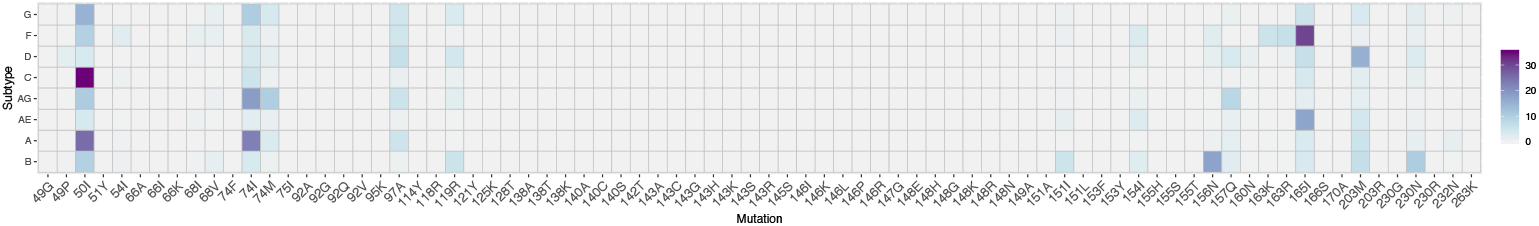
Prevalence (%) of INSTI resistance-associated mutations in treatment-naive patients. Mutations 49P, 50I, 54I, 68I/V, 74I/M, 97A, 119R, 151I, 154I, 156N, 157Q, 160N, 163K/R, 165I, 203M, 230N and 232N were observed with a frequency above 1% in at least one subtype **(Supplementary Material)**.

Figure 2 also provides a triplet entropy value of all subtypes combined for each position. High values, compared to within-subtype values, indicate between-subtype variation and suggest differences in predominant codon usage. Of the 41 integrase positions investigated, 16 positions (50, 51, 66, 75, 97, 121, 138, 142, 143, 145, 146, 149, 154, 155, 203 and 230) showed a similar predominant triplet across all subtypes. Of the remaining 25 positions, 11 positions showed a consensus in predominant *WT*_*t*_among 7 subtypes (49, 54, 74, 92, 118, 148, 153, 157, 160, 170, 263), 9 positions showed a consensus among 6 subtypes observed (68, 95, 119, 128, 140, 147, 156, 165, 166), and finally 5 positions (114, 125, 151, 163, 232) showed a consensus at 5 subtypes. These results illustrate well the extent of relevant integrate diversity within and between subtypes, encompassing varying entropy, resistance mutations and codon predominance. Some positions (e.g. 114 and 140) fully conserved at the amino acid level showed high variability in codon usage, while other positions (e.g. 50 and 74) showed limited variation in codon predominance but resistance-associated mutations occurred at increased frequencies.

### 3.3 Calculated genetic barrier

The observed baseline differences in triplet frequency between subtypes will only impact the genetic barrier to resistance when they are also accompanied by variation in triplet scores. Subtype frequencies of wild-type nucleotide triplets, translated amino acids and associated cumulative substitution costs are available as **Supplementary Material**.

Triplet score variability was subsequently investigated according to patterns in entropy and codon usage as described above. Limited variation in the evolutionary potential between subtypes is expected for the 35 resistance-associated mutations at 16 integrase positions with a similar predominant triplet and consequently similar scores. These positions are in addition generally characterised by a low frequency of resistance-associated mutations and a high level of consistent codon usage at baseline (**Supplementary Material**), although some exceptions exist with higher entropy values at positions 50, 138 and 203 (Figure 2) or resistance-associated mutations prevalent above 5% at low entropy positions 97 and 230 (Figure 3). Among the group of 25 positions, a larger heterogeneity in cost scores can be expected due to variable predominance complementing any variation in codon usage and mutation prevalence. The presence of resistance-associated mutations were 74I (subtypes A, G and CRF02 AG), 74M (CRF02 AG), 119R (subtype B), 151I (subtype B), 156N (subtype B), 157Q (CRF02 AG), 163K (subtype F), 163R (subtype F) an 165I (subtypes D, F, G and CRF01 AE) (Figure 3). In addition, subtype variation in predominant *WT*_*t*_ resulted most strongly in a different cost score for the following mutations: 54I, 68I, 74I/M, 92A/G, 95K, 119R, 125K, 128T, 140S, 147G, 148H/K/R, 151I, 156N, 160N, 163R, 165I, 166S, 170A, 232N and 263K (**Supplementary Material**).

Finally, combining into a complex interplay, reported results on genetic diversity and evolutionary costs can be captured into a single value denoting a population-based genetic potential for an INSTI mutation and subtype (Figure 4 and Figure 4). Significant differences in calculated genetic barrier were observed for 15 mutations at 12 positions (Figure 4), with a lower resistance potential for 50I (subtype D), 54I (CRF01 AE), 119R (subtypes C and F), 148H (subtype C), 151I (subtypes A, G and CRF02 AG) and 156N (subtype A and CRF01 AE) and 166S (CRF02 AG), while a higher potential for 68I (subtype A and CRF01 AE), 119R (CRF01 AE), 125K (subtype B), 140S (subtype B), 148R (subtype C), 151I (subtype B) and 156N (subtype B). Pairwise comparisons with subtype B revealed additional differences for 50I (subtypes A and C), 74I (subtypes A and AG), 163R (CRF01 AE) and 165I (subtype F and CRF01 AE) (**Supplementary Material**). A similar analysis applied to the RT enzyme revealed respectively a higher potential for 8 and a lower potential for 7 RTI resistance-associated mutations, including mutations at positions 106 and 65.

**Figure 4.**
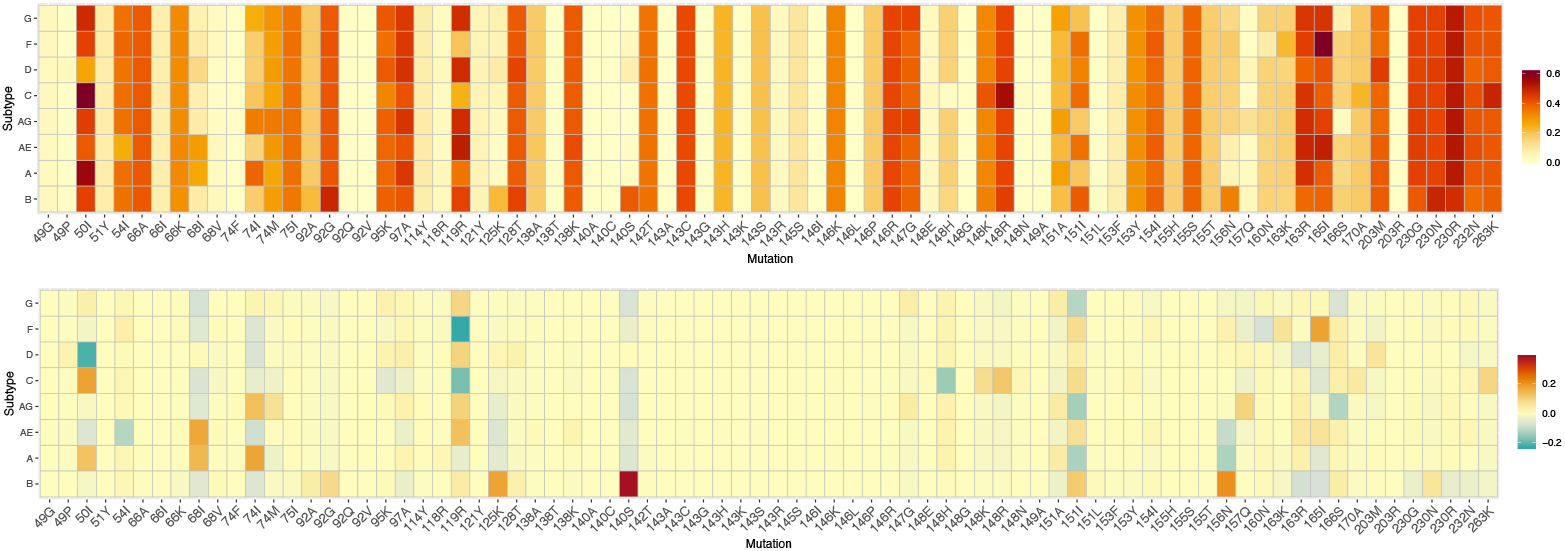
The upper panel shows the estimated population genetic barrier for each combination of a resistance-associated mutation and a subtype. A higher value (red) represents an increased potential for adaptation and is indicative for a lower genetic barrier to resistance. A subtype-specific value of this population genetic barrier is calculated by first assigning each wild-type triplet with a score that indicates the ease to evolve into the resistance amino acid (See the Methods section for a detailed description how this cost score for a wild-type triplet is derived). Next, the sum of all triplet scores is weighted by the triplet prevalence so that most frequent triplets have a larger impact on the population genetic barrier estimate. The lower panel shows for each subtype the average difference in population genetic barrier from the other subtypes. A lower genetic barrier to resistance for that subtype compared to the other subtypes is shown in red while a higher genetic barrier is shown in green.

The calculation of a wild type triplet score is based on the summation of all possible triplet cumulative costs, rather than only considering the minimum triplet cumulative cost. To illustrate the effect of this strategy, Figure 5 provides a comparison of the evolutionary potential calculated by our strategy and a simplified version which only considers the lowest cost for determine the cost. A modest impact on resistance potential was observed for a large number of resistance mutations, although for some mutations a large increase can be detected when almost equally likely paths were accounted for. Positions with high level of triplet variability (e.g. 119, 163, 230) are primarily expected to be affected across all subtypes, but important subtype-specific effects were also observed (e.g. 140S in subtype B, 148R and 263K in subtype C). **Supplementary Material** provides more comparative analyses on the impact of using empirically versus arbitrary defined substitution matrices.

**Figure 5.**
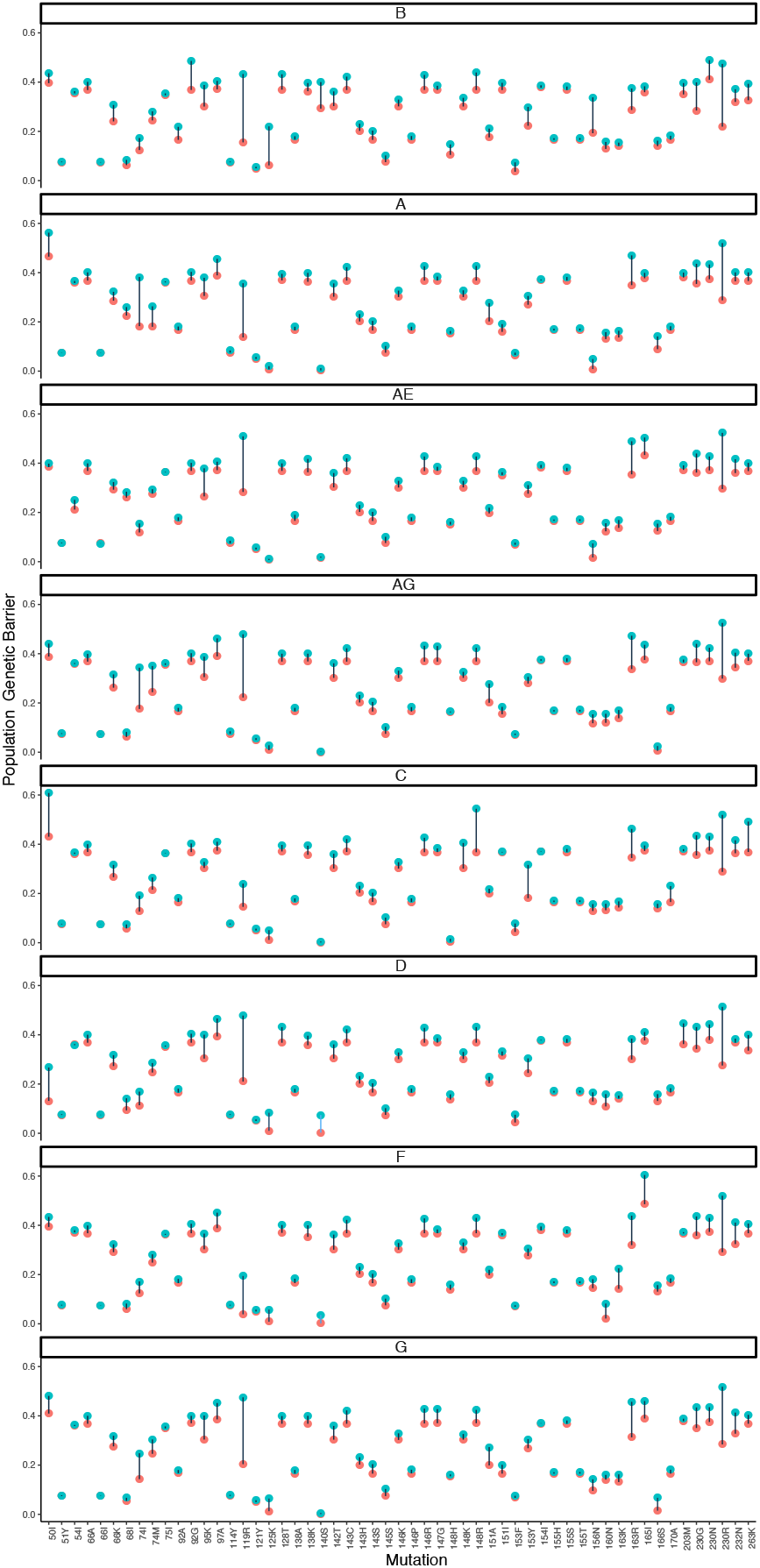
Each of 8 panels shows the estimated population genetic barrier for a single subtype, with the estimate either calculated as described in the Methods section (blue) or calculated by a simpler but similar methodology (red). Instead of obtaining the cost score for each wild-type triplet by summing all possible cumulative costs (blue), thereby taking into account almost equally likely substitution pathways of a wild-type triplet to the resistance amino acid, a cost score can also be calculated by only using the minimum cumulative cost (red) and therefore only considering the shortest substitution pathway to the resistance amino acid and ignoring other but almost equally likely mutational pathways to resistance. To increase the comparability of the two measures, we also took the negative exponential of the minimum score. Only increases in values are possible due to the summation, and we restricted the figure to the subset of mutations with a difference larger than 0.01.

## 4 Discussion

We developed an intuitive and novel methodology to quantify the evolutionary potential for viral escape from selective pressure in fast-evolving viruses, and presented an application evaluating the impact of genetic background in HIV-1 integrase on INSTI resistance development. The integrase enzyme has proven a successful drug target, and the objective of maximising the benefits of INSTIs is being translated into their global roll-out. However, as for other drug classes, the effectiveness of INSTIs can be challenged by HIV-1’s high rates of evolution when treatment options are non-optimal [2, 11, 29]. Viral adaptation to INSTI pressure through mutational pathways has been well reported, but the role of natural nucleotide variation on resistance development is less well understood [10, 17]. Additionally to resistance mutations present at baseline, differential codon usage can affect the mode and tempo of INSTI resistance pathways [23, 38]. This study presented here provided new insights into the processes underlying INSTI resistance development by quantifying the genetic barrier to resistance.

A large dataset of globally prevailing HIV-1 subtypes was investigated for their evolutionary potential for viral escape from all INSTIs available for HIV-1 treatment. This study confirms the non-polymorphic amino acid nature of most INSTI resistance positions [25, 35], as primarily single subtype-specific occurrences of resistance-associated mutations at low frequencies were detected. By contrast, variable codon predominance across subtypes was more pronounced and consequently we detected 15 resistance-associated mutations included in this study that significantly varied in estimated barrier in one or more subtypes. Modest variability in genetic barrier was observed for major INSTI resistance mutations except for 148H/R in subtype C which causes high-level resistance to all INSTIs [37]. More minor INSTI mutations, i.e. having an impact on drug activity in presence of a major mutation or compensating for defects in replication capacity [37], were characterised by increased variability in diversity and associated scores. Most notably was the varying potential for 140S in subtype B viruses, confirming previous findings [23, 32], which is an accessory mutation often occurring in combination with 148H. We and others have indeed previously reported the preference of the 148H pathway in treatment-experienced patients infected with subtype B viruses [17, 38].

Our framework improves related counting studies at several stages. Previous studies calculated the genetic barrier by only distinguishing between transitions and transversions, following an approach developed for the HIV-1 protease and reverse transcriptase enzymes [43]. We applied an empirically-derived and evolutionary-based cost for each possible substitution and triplet position, which resulted in an increased detection sensitivity, in particular transversions displayed substantial variation in costs compared to the arbitrary defined values in other studies. Furthermore, we take into account cost scores from all possible pathways, compared to restricting to the minimum score. This approach allowed to detect relevant increases in the estimated evolutionary potential (e.g. 74I), in particular for positions with high level of codon variability. To illustrate the improved sensitivity, the suggested subtype effect on the potential for 140C could not be confirmed in our study, as this mutation requires a substitution at the first triplet position, known to be evolutionary costly. In contrast to 140S, the prevalence of 140C in patients failing INSTI-based treatment does not differ across subtypes in the Stanford HIV drug resistance database [35], which strengthens our finding. A complete in-depth comparison of our findings with these studies is hindered by their limited number of HIV-1 subtypes and INSTI resistance mutations included. As most genetic barrier information to date has been established for the historical drug class of RTIs, we also applied our approach to the RT enzyme. Subtype-dependencies in genetic barrier both previously predicted and observed in clinical practice could be confirmed (e.g. 106M in subtype C). However, compared to related RTI counting studies [43], the increased sensitivity of our method also resulted in the identification of a higher potential for 65R in subtype C. A higher selection rate of 65R in this subtype has been well established [40] and attributed to a mechanistic basis of template factors [7]. However, an explanation by codon usage has been discarded [7], although we suggest an additional impact of codon usage when all possible paths are taken into account.

While our framework provides a population-based cost for a single position, it only represents a simplified estimate of the actual *in vivo* genetic barrier, which results from a multi-modal interplay that eventually defines the evolutionary time to drug resistance. Selection of a mutation and its rate of fixation heavily depend on factors such as patient adherence, treatment potency, impact on viral fitness and (non-additive) epistatic mutational interactions. Positions implicated in immune escape (e.g. 125) can also further influence the rate of resistance accumulation [8, 41]. It can be difficult to obtain information on all these influencing factors in a timely manner and hence to construct an accurate model capturing resistance evolutionary dynamics. When such an adequate model is lacking, our distance-based method offers a valuable alternative to assess the genetic barrier to resistance. Our methodology however can not predict novel resistance-associated mutations or subtype-specific pathways. A cost matrix was used for all subtypes assuming that substitution rates are equal between subtypes, however, the use of group-specific cost matrices can be accommodated by our framework.

In conclusion, the findings presented in this study are important in the context of up-scaled introduction of DTG and novel INSTIs in LMICs, where non-B subtypes prevail, although more studies are needed to further validate our results. Future applications of this reproducible framework do not only relate to novel HIV-1 drug classes or subtypes but, as principles of HIV-1 drug resistance are generally shared with other pathogens known to escape selective pressure, our methodology can be easily transferred to identify a role of genetic diversity of these pathogens, in particular for antivirals which are being evaluated in early development or clinical trials.

## Acknowledgements

Kristof Theys was funded by a post-doctoral grant from the Research Foundation - Flanders (FWO). Pieter Libin was funded by a PhD grant from the Research Foundation - Flanders (FWO). This work was supported by the Research Foundation - Flanders (FWO) - Flanders grant G.0692.14N and grant PF/10/018 from the KU Leuven. We sincerely thank Anne-Mieke Vandamme and late Ricardo Camacho for their support, valuable discussions and their heart full of warmth.

