## Supplementary Material for "An evolutionary-based approach to quantify the genetic barrier to drug resistance in fast-evolving viruses: an application to HIV-1 subtypes and integrase inhibitors"

May 14, 2019

#### 1 Genetic barrier to resistance of HIV-1 subtypes for RTIs

The developed framework to quantify the evolutionary potential to resistance was applied to HIV-1 inhibitors targeting the Reverse Transcriptase (RT) enzyme.

##### 1.1 Methods

Following a similar preprocessing analysis as described in the main manuscript, a dataset of 54903 RT sequences was collected including sequences from subtype A (n=3574), subtype B (n=28038), subtype C (n=9205), subtype D (n=1586), subtype F (n=812), subtype G (n=1301), CRF 01_AE (n=7635) and CRF 02_AG (n=2752).

A set of 77 resistance-associated mutations at 35 positions were included in the analysis. With respect to non-nucleoside RT inhibitors (NNRTI), the following mutations were considered 90I, 98G, 100I, 101E, 101H, 101P, 101Q, 101R, 101N, 103N, 103S, 103H, 103T, 103R, 103Q, 103E, 106A, 106M, 106I, 108I, 132M, 132L, 138K, 138A, 138G, 138Q, 138R, 179D, 179E, 179F, 179I, 179L, 179T, 181C, 181I, 181V, 188L, 188C, 188H, 190A, 190S, 190E, 190Q, 221Y, 225H, 227L, 227C, 230L, 230I, 232H, 234I, 236L, 238T and 238N. With respect to nucleo(s)(t)ide RT inhibitors (NRTI), the following mutations were considered: 41L, 65R, 67N, 70E, 70R, 74I, 74V, 75A, 75M, 75T, 75S, 77L, 115F, 118I, 151M, 184I, 184V, 210W, 215Y, 215F, 219E and 219Q.

##### 1.2 Results

Relevant genetic diversity at RT positions is shown in Figures 1 and 2. The prevalence of resistance-associated mutations is shown in Figure 1 while triplet entropy and amino acid entropy are shown in Figures 2.

The estimated evolutionary potential and the average difference between subtypes is shown in Figures 3 and 4. A lower evolutionary potential compared to the other subtypes was observed for 7 mutations: 77L (Subtype F), 106I in (Subtype C), 138G (Subtype C), 179I (CRF 02_AG and subtype G), 227L (Subtypes A, C, D and CRF 02_AG), 238T (CRF 01_AE), 238N (CRF 01_AE). A higher evolutionary potential compared to the other subtypes was observed for 8 mutations: 65R (Subtype C), 100I (Subtype D), 106M (Subtype C), 179E( Subtype G), 179I (Subtype A and CRF 01_AE), 179T (Subtype A),

190E (Subtype F) and 227L (Subtypes B, F and CRF 01_AE).

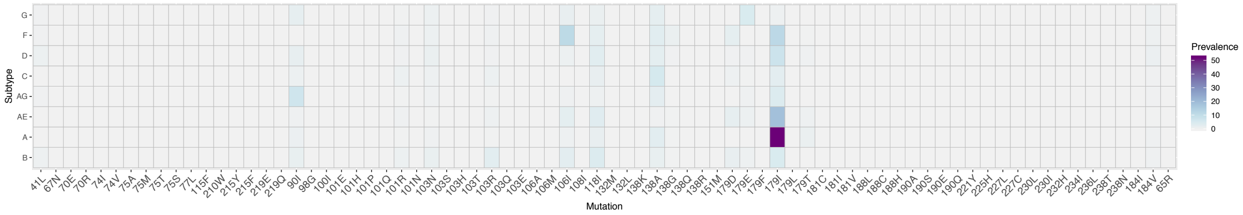


Figure 1: Prevalence (%) of resistance-associated mutations in the RT enzyme

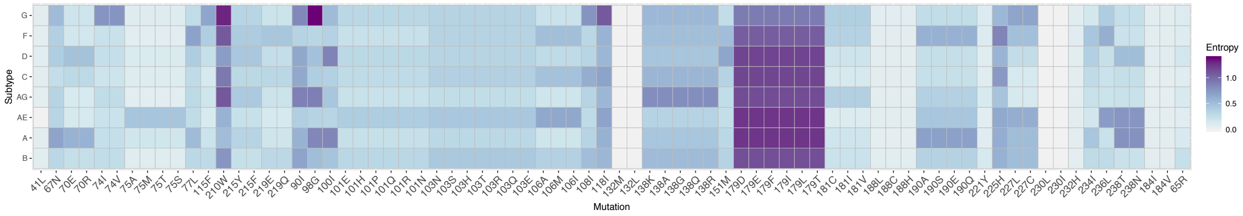


Figure 2: Triplet entropy values of resistance-associated mutations in the RT enzyme.

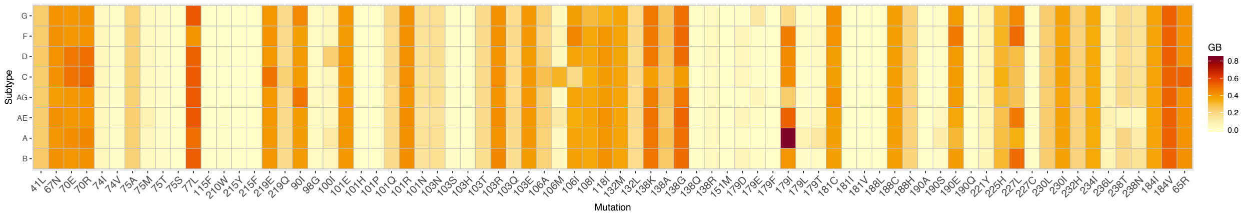


Figure 3: Population genetic barrier for HIV-1 subtypes and resistance-associated mutations in the RT enzyme.

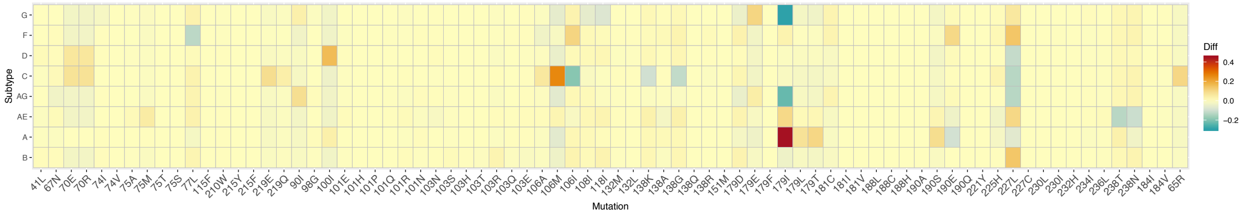


Figure 4: Average difference in population genetic barrier of one subtype compared to a weighted sample of other subtypes.

### 2 Data quality processing

#### 2.1 Codon ambiguities

As described in the main manuscript, at every resistance position, natural variability at the nucleotide level was assessed by determining the prevalence of nucleotide triplets in each HIV-1 subtype. Codons with ambiguities consisting of >2 bases per nucleotide position or of two or more ambiguities per codon were not considered. A nucleotide ambiguity of exact 2 bases was resolved in the two corresponding triplets, each counting for one half to their respective frequencies. All triplets with a prevalence above 1% in at least one HIV-1 subtype were retained for analysis.

Figures 5 shows graphically the percentage of codons per subtype and mutation that did not fulfil the inclusion criteria. It can be noticed that no major bias were encountered. More polymorphic positions are characterized by higher percentage of codons removed, which is can be expected.

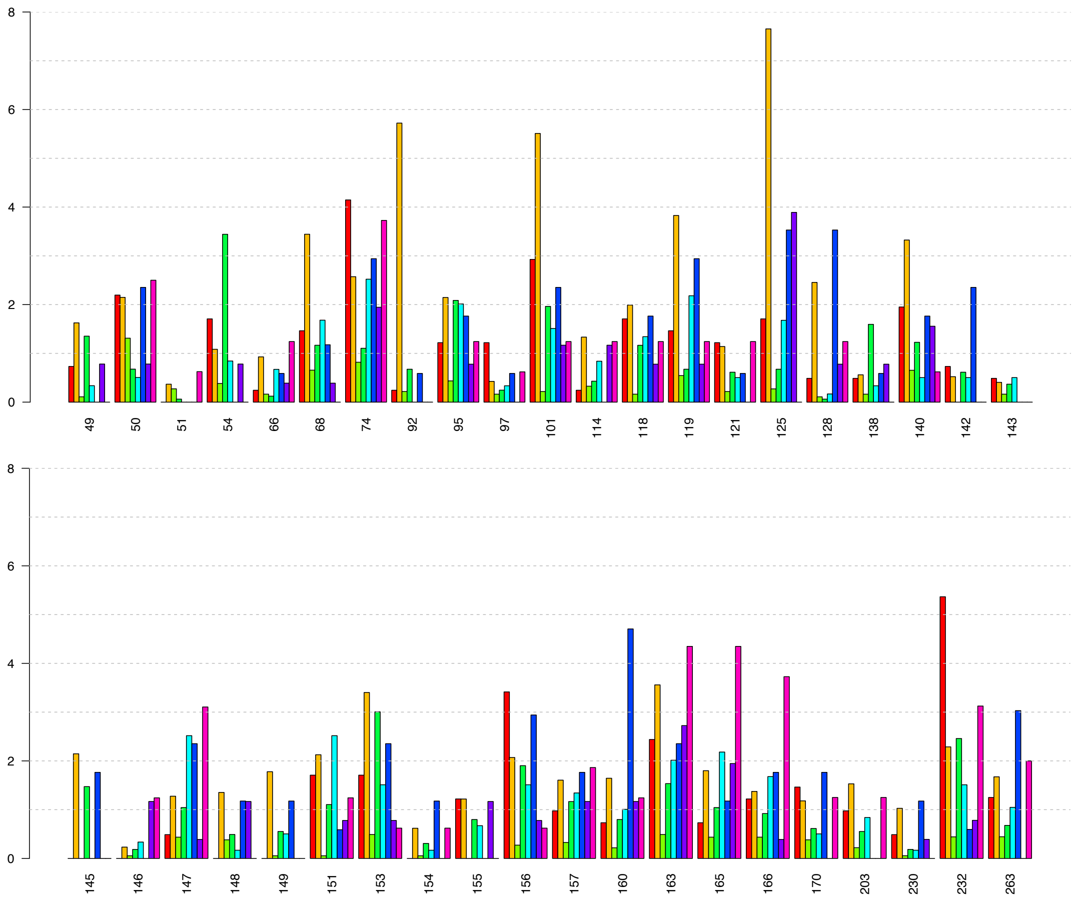


Figure 5: Prevalence (%) of codon ambiguities, grouped according to resistance mutation and subtype. Colours are as follows: subtype A (red), subtype B (yellow), subtype C (green), CRF01_AE (green), CRF02_AG (lightblue), subtype D (dark blue) F (purple), subtype G (pink).

#### 2.2 Subtype diversity distributions

Pairwise nucleotide diversity within each subtype was calculated to demonstrate homogeneity in subtype assignment within and between subtypes, visualized in Figure 6.

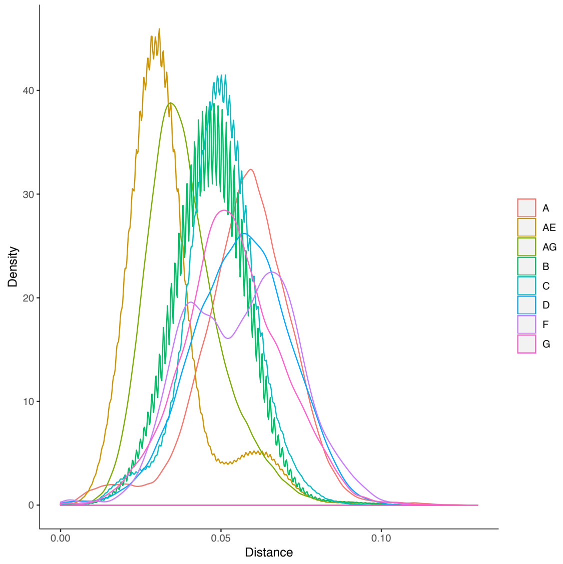


Figure 6: Pairwise distance distribution of each HIV-1 subtype.

#### 3 Substitution and cost matrices

A distinct substitution matrix for each of the three codon positions was derived from 964 HIV-1 integrase nucleotide sequences collected from Los Alamos database. The subtype distribution of these sequences was A (169), B (223), C (376), D (93), F (44), G (48), H (4), J (5) and K (2).

Estimated substitution: left = prob matrix, right = cost matrix.

| | A | T | C | G |
|---|---|---|---|---|
| A | - | 2.42 | 9.81 | 17.97 |
| T | 6.0 | - | 10.90 | 3.36 |
| C | 15.5 | 6.96 | - | 4.97 |
| G | 17.68 | 1.33 | 3.09 | - |

| | A | T | C | G |
|---|---|---|---|---|
| A | - | 7.4 | 1.8 | 1.0 |
| T | 3.0 | - | 1.6 | 5.3 |
| C | 1.2 | 2.6 | - | 3.6 |
| G | 1.0 | 13.5 | 5.8 | - |

| | A | T | C | G |
|---|---|---|---|---|
| A | - | 4.27 | 3.34 | 11.13 |
| T | 4.97 | - | 8.10 | 0.65 |
| C | 6.91 | 14.41 | - | 7.97 |
| G | 27.42 | 1.36 | 9.47 | - |

| | A | T | C | G |
|---|---|---|---|---|
| A | - | 6.4 | 8.2 | 2.5 |
| T | 5.5 | - | 3.4 | 42.2 |
| C | 4.0 | 1.9 | - | 3.4 |
| G | 1.0 | 20.2 | 2.9 | - |

| | A | T | C | G |
|---|---|---|---|---|
| A | - | 3.09 | 4.46 | 7.66 |
| T | 6.50 | - | 12.54 | 0.50 |
| C | 19.15 | 25.58 | - | 1.22 |
| G | 18.08 | 0.56 | 0.67 | - |

| | A | T | C | G |
|---|---|---|---|---|
| A | - | 8.3 | 5.7 | 3.3 |
| T | 3.9 | - | 2.0 | 51.2 |
| C | 1.3 | 1.0 | - | 21.0 |
| G | 1.4 | 45.7 | 38.2 | - |

Table 1: Upper: first codon position, Middle: second codon position, Lower: third codon position

### 4 Amino acid prevalence at resistance-associated positions

Table 2: Prevalence (%) of INSTI resistance-associated mutations across HIV-1 subtypes

| Table: Prevalence (%) of INSTI resistance-associated mutations across HIV-1 subtypes | | | | | | | | |
|---|---|---|---|---|---|---|---|---|
| **Mutation** | **Subtypes** | | | | | | | |
| | A | B | C | CRF01_AE | CRF02_AG | D | F | G |
| 49G | 0 | 0 | 0 | 0 | 0 | 0 | 0 | 0 |
| 49P | 0 | 0.4 | 0 | 0 | 0.2 | 1.8 | 0.2 | 0 |
| 50I | 25.2 | 9.4 | 35.1 | 3.8 | 10.6 | 3.5 | 9.4 | 15 |
| 51Y | 0 | 0 | 0 | 0 | 0 | 0 | 0 | 0 |
| 54I | 0.4 | 0.6 | 0.6 | 0.1 | 0 | 0 | 2.3 | 0 |
| 66A | 0 | 0 | 0 | 0 | 0 | 0 | 0 | 0 |
| 66I | 0 | 0 | 0 | 0 | 0 | 0 | 0 | 0 |
| 66K | 0 | 0 | 0 | 0 | 0 | 0 | 0 | 0 |
| 68I | 0 | 0.3 | 0 | 0.6 | 0 | 0.3 | 1.2 | 0 |
| 68V | 0 | 1.6 | 0.2 | 0.2 | 0.8 | 0 | 1.2 | 1.2 |
| 72I | 23.1 | 63.2 | 65.9 | 12.8 | 74.1 | 21.8 | 48.4 | 82.1 |
| 74F | 0 | 0 | 0 | 0 | 0 | 0 | 0 | 0 |
| 74I | 22.4 | 3.5 | 5 | 2.2 | 18.4 | 3.8 | 3.3 | 10.3 |
| 74M | 3.1 | 0.5 | 0.5 | 1.1 | 10.2 | 1.8 | 0.8 | 3.8 |
| 75I | 0 | 0 | 0 | 0 | 0 | 0 | 0 | 0 |
| 92A | 0 | 0 | 0 | 0 | 0 | 0 | 0 | 0 |
| 92G | 0 | 0 | 0 | 0 | 0 | 0 | 0 | 0 |
| 92Q | 0 | 0 | 0 | 0 | 0 | 0 | 0 | 0 |
| 92V | 0 | 0 | 0 | 0 | 0 | 0 | 0 | 0 |
| 95K | 0 | 0 | 0 | 0 | 0 | 0 | 0 | 0 |
| 97A | 5.1 | 0.5 | 1.1 | 0.7 | 5.5 | 6.2 | 4.7 | 4.7 |
| 101I | 25.6 | 44.8 | 94.6 | 9.6 | 89.3 | 26.5 | 87.8 | 79.2 |
| 114Y | 0 | 0 | 0 | 0 | 0 | 0 | 0 | 0 |
| 118R | 0 | 0 | 0 | 0 | 0 | 0 | 0 | 0 |
| 119R | 0.2 | 5.5 | 0.9 | 0.1 | 2 | 4.2 | 0.4 | 3.1 |
| 119S | 68.1 | 67.8 | 89.1 | 98 | 87.2 | 82.9 | 34.3 | 82.6 |
| 121Y | 0 | 0 | 0 | 0 | 0 | 0 | 0 | 0 |
| 124A | 69 | 21.3 | 61.9 | 84.3 | 90 | 75.8 | 58.5 | 62.7 |
| 125K | 0 | 0 | 0 | 0 | 0 | 0 | 0 | 0 |
| 128T | 0 | 0 | 0 | 0 | 0 | 0 | 0 | 0 |
| 138A | 0 | 0 | 0 | 0 | 0 | 0 | 0 | 0 |
| 138T | 0 | 0 | 0 | 0 | 0 | 0 | 0 | 0 |
| 138K | 0 | 0 | 0 | 0 | 0 | 0 | 0 | 0 |
| 140A | 0 | 0 | 0 | 0 | 0 | 0 | 0 | 0 |
| 140C | 0 | 0 | 0 | 0 | 0 | 0 | 0 | 0 |
| 140S | 0 | 0 | 0 | 0 | 0 | 0 | 0 | 0 |
| 142T | 0 | 0 | 0 | 0 | 0 | 0 | 0 | 0 |
| 143A | 0 | 0 | 0 | 0 | 0 | 0 | 0 | 0 |
| 143C | 0 | 0 | 0 | 0 | 0 | 0 | 0 | 0 |
| 143G | 0 | 0 | 0 | 0 | 0 | 0 | 0 | 0 |
| 143H | 0 | 0 | 0 | 0 | 0 | 0 | 0 | 0 |
| 143K | 0 | 0 | 0 | 0 | 0 | 0 | 0 | 0 |
| 143S | 0 | 0 | 0 | 0 | 0 | 0 | 0 | 0 |
| 143R | 0 | 0 | 0 | 0 | 0 | 0 | 0 | 0 |
| 145S | 0 | 0 | 0 | 0 | 0 | 0 | 0 | 0 |
| 146I | 0 | 0 | 0 | 0 | 0 | 0 | 0 | 0 |
| 146K | 0 | 0 | 0 | 0 | 0 | 0 | 0 | 0 |
| 146L | 0 | 0 | 0 | 0 | 0 | 0 | 0 | 0 |
| 146P | 0 | 0 | 0 | 0 | 0 | 0 | 0 | 0 |

| | | | | | | | | |
|---|---|---|---|---|---|---|---|---|
| 146R | 0 | 0 | 0 | 0 | 0 | 0 | 0 | 0 |
| 147G | 0 | 0 | 0 | 0 | 0 | 0 | 0 | 0 |
| 148E | 0 | 0 | 0 | 0 | 0 | 0 | 0 | 0 |
| 148H | 0 | 0 | 0 | 0 | 0 | 0 | 0 | 0 |
| 148G | 0 | 0 | 0 | 0 | 0 | 0 | 0 | 0 |
| 148K | 0 | 0 | 0 | 0 | 0 | 0 | 0 | 0 |
| 148R | 0 | 0 | 0 | 0 | 0 | 0 | 0 | 0 |
| 148N | 0 | 0 | 0 | 0 | 0 | 0 | 0 | 0 |
| 149A | 0 | 0 | 0 | 0 | 0 | 0 | 0 | 0 |
| 151A | 0 | 0 | 0 | 0 | 0 | 0 | 0 | 0 |
| 151I | 0 | 5.2 | 0.3 | 1.5 | 0.3 | 0 | 0.8 | 0.6 |
| 151L | 0 | 0 | 0 | 0 | 0 | 0 | 0 | 0 |
| 153F | 0 | 0 | 0 | 0 | 0 | 0 | 0 | 0 |
| 153Y | 0 | 0 | 0 | 0 | 0 | 0 | 0 | 0 |
| 154I | 0.2 | 2.4 | 0.2 | 3 | 0.9 | 1.5 | 3.1 | 0 |
| 155H | 0 | 0 | 0 | 0 | 0 | 0 | 0 | 0 |
| 155S | 0 | 0 | 0 | 0 | 0 | 0 | 0 | 0 |
| 155T | 0 | 0 | 0 | 0 | 0 | 0 | 0 | 0 |
| 156N | 0.5 | 17.6 | 0.3 | 0.1 | 0.2 | 1.5 | 2.3 | 0 |
| 157Q | 1.7 | 2.3 | 0.4 | 1.1 | 8.3 | 3.5 | 0 | 0.9 |
| 160N | 0 | 0.2 | 0.1 | 0.2 | 0.2 | 1.2 | 0 | 0 |
| 163K | 0.1 | 0 | 0 | 0 | 0 | 0 | 5.4 | 0 |
| 163R | 0.1 | 0.2 | 0 | 0 | 0 | 0.6 | 6.2 | 0 |
| 165I | 3.2 | 3.4 | 3.9 | 17.2 | 1.8 | 6.2 | 30.2 | 5.3 |
| 166S | 0 | 0 | 0 | 0 | 0 | 0 | 0 | 0 |
| 170A | 0 | 0 | 0 | 0 | 0 | 0 | 0 | 0 |
| 201I | 95.1 | 41 | 93.7 | 98.7 | 98.4 | 97.7 | 95.9 | 96.9 |
| 203M | 5.5 | 6.4 | 2.2 | 4.2 | 1.7 | 15.3 | 0.8 | 3.4 |
| 203R | 0 | 0 | 0 | 0 | 0 | 0 | 0 | 0 |
| 206S | 5.9 | 14.1 | 8.9 | 4.5 | 95 | 14.7 | 15 | 93.8 |
| 230G | 0 | 0 | 0 | 0 | 0 | 0 | 0 | 0 |
| 230N | 1.5 | 10.4 | 1.5 | 0.7 | 0.3 | 2.9 | 1.2 | 1.9 |
| 230R | 0 | 0 | 0 | 0 | 0 | 0 | 0 | 0 |
| 232N | 1.5 | 0.2 | 0 | 0 | 0 | 0 | 0 | 0.6 |
| 263K | 0 | 0 | 0 | 0 | 0 | 0 | 0 | 0 |

### 5 Triplet prevalence and costs across HIV-1 subtypes

Table 3: Prevalence of wild type triplet per position, resistance amino acid (AA) and subtype together with associated cost to resistance triplet

| Table: Prevalence of wild type triplet per position, resistance amino acid (AA) and subtype together with associated cost to resistance triplet. | | | | | | | | | | | | | |
|---|---|---|---|---|---|---|---|---|---|---|---|---|---|
| Position | Wild Type | | Subtype | | | | | | | | Resistance | | Cost |
| | AA | Triplet | A | B | C | CRF01_AE | CRF02_AG | D | F | G | AA | Triplet | |
| 49 | A | GCC | 73.5 | 92.6 | 94.4 | 8.7 | 96.8 | 91.8 | 95.5 | 95.6 | G | GGT | 5.5 |
| 49 | A | GCC | 73.5 | 92.6 | 94.4 | 8.7 | 96.8 | 91.8 | 95.5 | 95.6 | G | GGC | 3.6 |
| 49 | A | GCC | 73.5 | 92.6 | 94.4 | 8.7 | 96.8 | 91.8 | 95.5 | 95.6 | G | GGA | 7.6 |
| 49 | A | GCC | 73.5 | 92.6 | 94.4 | 8.7 | 96.8 | 91.8 | 95.5 | 95.6 | G | GGG | 7 |
| 49 | A | GCT | 25 | 3.7 | 4 | 88.9 | 2.1 | 3.5 | 1.6 | 4.4 | G | GGT | 3.6 |
| 49 | A | GCT | 25 | 3.7 | 4 | 88.9 | 2.1 | 3.5 | 1.6 | 4.4 | G | GGC | 7 |
| 49 | A | GCT | 25 | 3.7 | 4 | 88.9 | 2.1 | 3.5 | 1.6 | 4.4 | G | GGA | 9.1 |
| 49 | A | GCT | 25 | 3.7 | 4 | 88.9 | 2.1 | 3.5 | 1.6 | 4.4 | G | GGG | 45.8 |
| 49 | A | GCG | 0.2 | 2.4 | 0.1 | 0.2 | 0.1 | 1.8 | 0 | 0 | G | GGT | 23.8 |
| 49 | A | GCG | 0.2 | 2.4 | 0.1 | 0.2 | 0.1 | 1.8 | 0 | 0 | G | GGC | 6.5 |
| 49 | A | GCG | 0.2 | 2.4 | 0.1 | 0.2 | 0.1 | 1.8 | 0 | 0 | G | GGA | 4.6 |
| 49 | A | GCG | 0.2 | 2.4 | 0.1 | 0.2 | 0.1 | 1.8 | 0 | 0 | G | GGG | 3.6 |
| 49 | A | GCA | 0.7 | 0.9 | 1.3 | 2 | 0.8 | 1.2 | 2.4 | 0 | G | GGT | 10 |
| 49 | A | GCA | 0.7 | 0.9 | 1.3 | 2 | 0.8 | 1.2 | 2.4 | 0 | G | GGC | 11.8 |
| 49 | A | GCA | 0.7 | 0.9 | 1.3 | 2 | 0.8 | 1.2 | 2.4 | 0 | G | GGA | 3.6 |
| 49 | A | GCA | 0.7 | 0.9 | 1.3 | 2 | 0.8 | 1.2 | 2.4 | 0 | G | GGG | 6.1 |
| 49 | P | CCC | 0 | 0.4 | 0 | 0 | 0.2 | 1.8 | 0.2 | 0 | G | GGT | 8 |
| 49 | P | CCC | 0 | 0.4 | 0 | 0 | 0.2 | 1.8 | 0.2 | 0 | G | GGC | 7 |
| 49 | P | CCC | 0 | 0.4 | 0 | 0 | 0.2 | 1.8 | 0.2 | 0 | G | GGA | 8.3 |
| 49 | P | CCC | 0 | 0.4 | 0 | 0 | 0.2 | 1.8 | 0.2 | 0 | G | GGG | 28 |
| 49 | A | GCC | 73.5 | 92.6 | 94.4 | 8.7 | 96.8 | 91.8 | 95.5 | 95.6 | P | CCT | 7.7 |
| 49 | A | GCC | 73.5 | 92.6 | 94.4 | 8.7 | 96.8 | 91.8 | 95.5 | 95.6 | P | CCC | 5.8 |
| 49 | A | GCC | 73.5 | 92.6 | 94.4 | 8.7 | 96.8 | 91.8 | 95.5 | 95.6 | P | CCA | 9.8 |
| 49 | A | GCC | 73.5 | 92.6 | 94.4 | 8.7 | 96.8 | 91.8 | 95.5 | 95.6 | P | CCG | 9.2 |
| 49 | A | GCT | 25 | 3.7 | 4 | 88.9 | 2.1 | 3.5 | 1.6 | 4.4 | P | CCT | 5.8 |
| 49 | A | GCT | 25 | 3.7 | 4 | 88.9 | 2.1 | 3.5 | 1.6 | 4.4 | P | CCC | 9.2 |
| 49 | A | GCT | 25 | 3.7 | 4 | 88.9 | 2.1 | 3.5 | 1.6 | 4.4 | P | CCA | 11.3 |
| 49 | A | GCT | 25 | 3.7 | 4 | 88.9 | 2.1 | 3.5 | 1.6 | 4.4 | P | CCG | 48 |
| 49 | A | GCG | 0.2 | 2.4 | 0.1 | 0.2 | 0.1 | 1.8 | 0 | 0 | P | CCT | 26 |
| 49 | A | GCG | 0.2 | 2.4 | 0.1 | 0.2 | 0.1 | 1.8 | 0 | 0 | P | CCC | 8.7 |

| | | | | | | | | | | | | |
|---|---|---|---|---|---|---|---|---|---|---|---|---|
| 49 | A | GCG | 0.2 | 2.4 | 0.1 | 0.2 | 0.1 | 1.8 | 0 | 0 | P | CCA | 6.8 |
| 49 | A | GCG | 0.2 | 2.4 | 0.1 | 0.2 | 0.1 | 1.8 | 0 | 0 | P | CCG | 5.8 |
| 49 | A | GCA | 0.7 | 0.9 | 1.3 | 2 | 0.8 | 1.2 | 2.4 | 0 | P | CCT | 12.2 |
| 49 | A | GCA | 0.7 | 0.9 | 1.3 | 2 | 0.8 | 1.2 | 2.4 | 0 | P | CCC | 14 |
| 49 | A | GCA | 0.7 | 0.9 | 1.3 | 2 | 0.8 | 1.2 | 2.4 | 0 | P | CCA | 5.8 |
| 49 | A | GCA | 0.7 | 0.9 | 1.3 | 2 | 0.8 | 1.2 | 2.4 | 0 | P | CCG | 8.3 |
| 49 | P | CCC | 0 | 0.4 | 0 | 0 | 0.2 | 1.8 | 0.2 | 0 | P | CCT | 2.6 |
| 49 | P | CCC | 0 | 0.4 | 0 | 0 | 0.2 | 1.8 | 0.2 | 0 | P | CCC | 0 |
| 49 | P | CCC | 0 | 0.4 | 0 | 0 | 0.2 | 1.8 | 0.2 | 0 | P | CCA | 1.2 |
| 49 | P | CCC | 0 | 0.4 | 0 | 0 | 0.2 | 1.8 | 0.2 | 0 | P | CCG | 3.6 |
| 50 | M | ATG | 72.1 | 87.7 | 51.3 | 95.3 | 85.2 | 58.8 | 87.1 | 82.5 | I | ATT | 13.5 |
| 50 | M | ATG | 72.1 | 87.7 | 51.3 | 95.3 | 85.2 | 58.8 | 87.1 | 82.5 | I | ATC | 5.8 |
| 50 | M | ATG | 72.1 | 87.7 | 51.3 | 95.3 | 85.2 | 58.8 | 87.1 | 82.5 | I | ATA | 1 |
| 50 | I | ATA | 25.2 | 9.4 | 35.1 | 3.8 | 10.6 | 3.5 | 9.4 | 15 | I | ATT | 7.4 |
| 50 | I | ATA | 25.2 | 9.4 | 35.1 | 3.8 | 10.6 | 3.5 | 9.4 | 15 | I | ATC | 1.8 |
| 50 | I | ATA | 25.2 | 9.4 | 35.1 | 3.8 | 10.6 | 3.5 | 9.4 | 15 | I | ATA | 0 |
| 50 | T | ACA | 0.9 | 0.5 | 11.5 | 0 | 0.7 | 0 | 1.2 | 0 | I | ATT | 9 |
| 50 | T | ACA | 0.9 | 0.5 | 11.5 | 0 | 0.7 | 0 | 1.2 | 0 | I | ATC | 10.8 |
| 50 | T | ACA | 0.9 | 0.5 | 11.5 | 0 | 0.7 | 0 | 1.2 | 0 | I | ATA | 2.6 |
| 50 | T | ACG | 0.2 | 0.8 | 0.4 | 0 | 2.1 | 0.3 | 0.4 | 1.9 | I | ATT | 22.8 |
| 50 | T | ACG | 0.2 | 0.8 | 0.4 | 0 | 2.1 | 0.3 | 0.4 | 1.9 | I | ATC | 5.5 |
| 50 | T | ACG | 0.2 | 0.8 | 0.4 | 0 | 2.1 | 0.3 | 0.4 | 1.9 | I | ATA | 3.6 |
| 50 | L | TTG | 0.2 | 0.2 | 0.1 | 0.1 | 0 | 34.4 | 0.4 | 0 | I | ATT | 23.2 |
| 50 | L | TTG | 0.2 | 0.2 | 0.1 | 0.1 | 0 | 34.4 | 0.4 | 0 | I | ATC | 5.9 |
| 50 | L | TTG | 0.2 | 0.2 | 0.1 | 0.1 | 0 | 34.4 | 0.4 | 0 | I | ATA | 4 |
| 50 | L | TTA | 0.2 | 0 | 0.4 | 0 | 0 | 1.8 | 0 | 0 | I | ATT | 9.4 |
| 50 | L | TTA | 0.2 | 0 | 0.4 | 0 | 0 | 1.8 | 0 | 0 | I | ATC | 11.2 |
| 50 | L | TTA | 0.2 | 0 | 0.4 | 0 | 0 | 1.8 | 0 | 0 | I | ATA | 3 |
| 51 | H | CAT | 99.3 | 98.9 | 98.5 | 99.2 | 100 | 99.4 | 100 | 99.7 | Y | TAT | 2.6 |
| 51 | H | CAT | 99.3 | 98.9 | 98.5 | 99.2 | 100 | 99.4 | 100 | 99.7 | Y | TAC | 6 |
| 51 | H | CAC | 0.2 | 0.9 | 1.1 | 0.7 | 0 | 0.6 | 0 | 0.3 | Y | TAT | 4.5 |
| 51 | H | CAC | 0.2 | 0.9 | 1.1 | 0.7 | 0 | 0.6 | 0 | 0.3 | Y | TAC | 2.6 |
| 54 | V | GTA | 96.6 | 94.8 | 96.3 | 44.4 | 97.2 | 95.9 | 94.9 | 97.5 | I | ATT | 7.4 |
| 54 | V | GTA | 96.6 | 94.8 | 96.3 | 44.4 | 97.2 | 95.9 | 94.9 | 97.5 | I | ATC | 9.2 |

| | | | | | | | | | | | | | |
|---|---|---|---|---|---|---|---|---|---|---|---|---|---|
| 54 | V | GTA | 96.6 | 94.8 | 96.3 | 44.4 | 97.2 | 95.9 | 94.9 | 97.5 | I | ATA | 1 |
| 54 | V | GTG | 3.1 | 4.5 | 2.8 | 55.4 | 2.7 | 2.9 | 2.1 | 2.5 | I | ATT | 21.2 |
| 54 | V | GTG | 3.1 | 4.5 | 2.8 | 55.4 | 2.7 | 2.9 | 2.1 | 2.5 | I | ATC | 3.9 |
| 54 | V | GTG | 3.1 | 4.5 | 2.8 | 55.4 | 2.7 | 2.9 | 2.1 | 2.5 | I | ATA | 2 |
| 54 | I | ATA | 0.4 | 0.6 | 0.6 | 0.1 | 0 | 0 | 2.3 | 0 | I | ATT | 7.4 |
| 54 | I | ATA | 0.4 | 0.6 | 0.6 | 0.1 | 0 | 0 | 2.3 | 0 | I | ATC | 1.8 |
| 54 | I | ATA | 0.4 | 0.6 | 0.6 | 0.1 | 0 | 0 | 2.3 | 0 | I | ATA | 0 |
| 66 | T | ACA | 97.2 | 93.7 | 96.1 | 98.6 | 96.8 | 96.8 | 99 | 97.5 | A | GCT | 7.4 |
| 66 | T | ACA | 97.2 | 93.7 | 96.1 | 98.6 | 96.8 | 96.8 | 99 | 97.5 | A | GCC | 9.2 |
| 66 | T | ACA | 97.2 | 93.7 | 96.1 | 98.6 | 96.8 | 96.8 | 99 | 97.5 | A | GCA | 1 |
| 66 | T | ACA | 97.2 | 93.7 | 96.1 | 98.6 | 96.8 | 96.8 | 99 | 97.5 | A | GCG | 3.5 |
| 66 | T | ACC | 1.5 | 5 | 2.4 | 0.6 | 1.5 | 2.9 | 1 | 1.6 | A | GCT | 2.9 |
| 66 | T | ACC | 1.5 | 5 | 2.4 | 0.6 | 1.5 | 2.9 | 1 | 1.6 | A | GCC | 1 |
| 66 | T | ACC | 1.5 | 5 | 2.4 | 0.6 | 1.5 | 2.9 | 1 | 1.6 | A | GCA | 5 |
| 66 | T | ACC | 1.5 | 5 | 2.4 | 0.6 | 1.5 | 2.9 | 1 | 1.6 | A | GCG | 4.4 |
| 66 | T | ACT | 0 | 1 | 0.8 | 0.2 | 1.6 | 0 | 0 | 0.6 | A | GCT | 1 |
| 66 | T | ACT | 0 | 1 | 0.8 | 0.2 | 1.6 | 0 | 0 | 0.6 | A | GCC | 4.4 |
| 66 | T | ACT | 0 | 1 | 0.8 | 0.2 | 1.6 | 0 | 0 | 0.6 | A | GCA | 6.5 |
| 66 | T | ACT | 0 | 1 | 0.8 | 0.2 | 1.6 | 0 | 0 | 0.6 | A | GCG | 43.2 |
| 66 | T | ACG | 1.3 | 0.3 | 0.5 | 0.4 | 0.1 | 0.3 | 0 | 0.3 | A | GCT | 21.2 |
| 66 | T | ACG | 1.3 | 0.3 | 0.5 | 0.4 | 0.1 | 0.3 | 0 | 0.3 | A | GCC | 3.9 |
| 66 | T | ACG | 1.3 | 0.3 | 0.5 | 0.4 | 0.1 | 0.3 | 0 | 0.3 | A | GCA | 2 |
| 66 | T | ACG | 1.3 | 0.3 | 0.5 | 0.4 | 0.1 | 0.3 | 0 | 0.3 | A | GCG | 1 |
| 66 | T | ACA | 97.2 | 93.7 | 96.1 | 98.6 | 96.8 | 96.8 | 99 | 97.5 | I | ATT | 9 |
| 66 | T | ACA | 97.2 | 93.7 | 96.1 | 98.6 | 96.8 | 96.8 | 99 | 97.5 | I | ATC | 10.8 |
| 66 | T | ACA | 97.2 | 93.7 | 96.1 | 98.6 | 96.8 | 96.8 | 99 | 97.5 | I | ATA | 2.6 |
| 66 | T | ACC | 1.5 | 5 | 2.4 | 0.6 | 1.5 | 2.9 | 1 | 1.6 | I | ATT | 4.5 |
| 66 | T | ACC | 1.5 | 5 | 2.4 | 0.6 | 1.5 | 2.9 | 1 | 1.6 | I | ATC | 2.6 |
| 66 | T | ACC | 1.5 | 5 | 2.4 | 0.6 | 1.5 | 2.9 | 1 | 1.6 | I | ATA | 6.6 |
| 66 | T | ACT | 0 | 1 | 0.8 | 0.2 | 1.6 | 0 | 0 | 0.6 | I | ATT | 2.6 |
| 66 | T | ACT | 0 | 1 | 0.8 | 0.2 | 1.6 | 0 | 0 | 0.6 | I | ATC | 6 |
| 66 | T | ACT | 0 | 1 | 0.8 | 0.2 | 1.6 | 0 | 0 | 0.6 | I | ATA | 8.1 |
| 66 | T | ACG | 1.3 | 0.3 | 0.5 | 0.4 | 0.1 | 0.3 | 0 | 0.3 | I | ATT | 22.8 |
| 66 | T | ACG | 1.3 | 0.3 | 0.5 | 0.4 | 0.1 | 0.3 | 0 | 0.3 | I | ATC | 5.5 |

| | | | | | | | | | | | | | |
|---|---|---|---|---|---|---|---|---|---|---|---|---|---|
| 66 | T | ACG | 1.3 | 0.3 | 0.5 | 0.4 | 0.1 | 0.3 | 0 | 0.3 | I | ATA | 3.6 |
| 66 | T | ACA | 97.2 | 93.7 | 96.1 | 98.6 | 96.8 | 96.8 | 99 | 97.5 | K | AAA | 1.2 |
| 66 | T | ACA | 97.2 | 93.7 | 96.1 | 98.6 | 96.8 | 96.8 | 99 | 97.5 | K | AAG | 3.7 |
| 66 | T | ACC | 1.5 | 5 | 2.4 | 0.6 | 1.5 | 2.9 | 1 | 1.6 | K | AAA | 5.2 |
| 66 | T | ACC | 1.5 | 5 | 2.4 | 0.6 | 1.5 | 2.9 | 1 | 1.6 | K | AAG | 4.6 |
| 66 | T | ACT | 0 | 1 | 0.8 | 0.2 | 1.6 | 0 | 0 | 0.6 | K | AAA | 6.7 |
| 66 | T | ACT | 0 | 1 | 0.8 | 0.2 | 1.6 | 0 | 0 | 0.6 | K | AAG | 43.4 |
| 66 | T | ACG | 1.3 | 0.3 | 0.5 | 0.4 | 0.1 | 0.3 | 0 | 0.3 | K | AAA | 2.2 |
| 66 | T | ACG | 1.3 | 0.3 | 0.5 | 0.4 | 0.1 | 0.3 | 0 | 0.3 | K | AAG | 1.2 |
| 68 | L | TTA | 15 | 82.6 | 88 | 6.9 | 85.3 | 60 | 89.3 | 90.1 | I | ATT | 9.4 |
| 68 | L | TTA | 15 | 82.6 | 88 | 6.9 | 85.3 | 60 | 89.3 | 90.1 | I | ATC | 11.2 |
| 68 | L | TTA | 15 | 82.6 | 88 | 6.9 | 85.3 | 60 | 89.3 | 90.1 | I | ATA | 3 |
| 68 | L | CTA | 82.7 | 9.8 | 9.2 | 89.3 | 12.1 | 35 | 6 | 5.6 | I | ATT | 7.6 |
| 68 | L | CTA | 82.7 | 9.8 | 9.2 | 89.3 | 12.1 | 35 | 6 | 5.6 | I | ATC | 9.4 |
| 68 | L | CTA | 82.7 | 9.8 | 9.2 | 89.3 | 12.1 | 35 | 6 | 5.6 | I | ATA | 1.2 |
| 68 | L | CTG | 2.1 | 2.5 | 0.6 | 2.6 | 0.1 | 1.8 | 0.4 | 0 | I | ATT | 21.4 |
| 68 | L | CTG | 2.1 | 2.5 | 0.6 | 2.6 | 0.1 | 1.8 | 0.4 | 0 | I | ATC | 4.1 |
| 68 | L | CTG | 2.1 | 2.5 | 0.6 | 2.6 | 0.1 | 1.8 | 0.4 | 0 | I | ATA | 2.2 |
| 68 | L | TTG | 0 | 2.5 | 2 | 0.2 | 1.7 | 2.9 | 1.2 | 3.1 | I | ATT | 23.2 |
| 68 | L | TTG | 0 | 2.5 | 2 | 0.2 | 1.7 | 2.9 | 1.2 | 3.1 | I | ATC | 5.9 |
| 68 | L | TTG | 0 | 2.5 | 2 | 0.2 | 1.7 | 2.9 | 1.2 | 3.1 | I | ATA | 4 |
| 68 | V | GTA | 0 | 1.6 | 0.2 | 0.2 | 0.8 | 0 | 1.2 | 1.2 | I | ATT | 7.4 |
| 68 | V | GTA | 0 | 1.6 | 0.2 | 0.2 | 0.8 | 0 | 1.2 | 1.2 | I | ATC | 9.2 |
| 68 | V | GTA | 0 | 1.6 | 0.2 | 0.2 | 0.8 | 0 | 1.2 | 1.2 | I | ATA | 1 |
| 68 | I | ATA | 0 | 0.3 | 0 | 0.6 | 0 | 0.3 | 1.2 | 0 | I | ATT | 7.4 |
| 68 | I | ATA | 0 | 0.3 | 0 | 0.6 | 0 | 0.3 | 1.2 | 0 | I | ATC | 1.8 |
| 68 | I | ATA | 0 | 0.3 | 0 | 0.6 | 0 | 0.3 | 1.2 | 0 | I | ATA | 0 |
| 68 | L | TTA | 15 | 82.6 | 88 | 6.9 | 85.3 | 60 | 89.3 | 90.1 | V | GTT | 11.7 |
| 68 | L | TTA | 15 | 82.6 | 88 | 6.9 | 85.3 | 60 | 89.3 | 90.1 | V | GTC | 13.5 |
| 68 | L | TTA | 15 | 82.6 | 88 | 6.9 | 85.3 | 60 | 89.3 | 90.1 | V | GTA | 5.3 |
| 68 | L | TTA | 15 | 82.6 | 88 | 6.9 | 85.3 | 60 | 89.3 | 90.1 | V | GTG | 7.8 |
| 68 | L | CTA | 82.7 | 9.8 | 9.2 | 89.3 | 12.1 | 35 | 6 | 5.6 | V | GTT | 10 |
| 68 | L | CTA | 82.7 | 9.8 | 9.2 | 89.3 | 12.1 | 35 | 6 | 5.6 | V | GTC | 11.8 |
| 68 | L | CTA | 82.7 | 9.8 | 9.2 | 89.3 | 12.1 | 35 | 6 | 5.6 | V | GTA | 3.6 |

| | | | | | | | | | | | | |
|---|---|---|---|---|---|---|---|---|---|---|---|---|
| 68 | L | CTA | 82.7 | 9.8 | 9.2 | 89.3 | 12.1 | 35 | 6 | 5.6 | V | GTG | 6.1 |
| 68 | L | CTG | 2.1 | 2.5 | 0.6 | 2.6 | 0.1 | 1.8 | 0.4 | 0 | V | GTT | 23.8 |
| 68 | L | CTG | 2.1 | 2.5 | 0.6 | 2.6 | 0.1 | 1.8 | 0.4 | 0 | V | GTC | 6.5 |
| 68 | L | CTG | 2.1 | 2.5 | 0.6 | 2.6 | 0.1 | 1.8 | 0.4 | 0 | V | GTA | 4.6 |
| 68 | L | CTG | 2.1 | 2.5 | 0.6 | 2.6 | 0.1 | 1.8 | 0.4 | 0 | V | GTG | 3.6 |
| 68 | L | TTG | 0 | 2.5 | 2 | 0.2 | 1.7 | 2.9 | 1.2 | 3.1 | V | GTT | 25.5 |
| 68 | L | TTG | 0 | 2.5 | 2 | 0.2 | 1.7 | 2.9 | 1.2 | 3.1 | V | GTC | 8.2 |
| 68 | L | TTG | 0 | 2.5 | 2 | 0.2 | 1.7 | 2.9 | 1.2 | 3.1 | V | GTA | 6.3 |
| 68 | L | TTG | 0 | 2.5 | 2 | 0.2 | 1.7 | 2.9 | 1.2 | 3.1 | V | GTG | 5.3 |
| 68 | V | GTA | 0 | 1.6 | 0.2 | 0.2 | 0.8 | 0 | 1.2 | 1.2 | V | GTT | 7.4 |
| 68 | V | GTA | 0 | 1.6 | 0.2 | 0.2 | 0.8 | 0 | 1.2 | 1.2 | V | GTC | 1.8 |
| 68 | V | GTA | 0 | 1.6 | 0.2 | 0.2 | 0.8 | 0 | 1.2 | 1.2 | V | GTA | 0 |
| 68 | V | GTA | 0 | 1.6 | 0.2 | 0.2 | 0.8 | 0 | 1.2 | 1.2 | V | GTG | 1 |
| 68 | I | ATA | 0 | 0.3 | 0 | 0.6 | 0 | 0.3 | 1.2 | 0 | V | GTT | 7.4 |
| 68 | I | ATA | 0 | 0.3 | 0 | 0.6 | 0 | 0.3 | 1.2 | 0 | V | GTC | 9.2 |
| 68 | I | ATA | 0 | 0.3 | 0 | 0.6 | 0 | 0.3 | 1.2 | 0 | V | GTA | 1 |
| 68 | I | ATA | 0 | 0.3 | 0 | 0.6 | 0 | 0.3 | 1.2 | 0 | V | GTG | 3.5 |
| 72 | I | ATT | 9.9 | 60.8 | 3.7 | 1.2 | 70.5 | 21.8 | 46.1 | 78.6 | I | ATT | 0 |
| 72 | I | ATT | 9.9 | 60.8 | 3.7 | 1.2 | 70.5 | 21.8 | 46.1 | 78.6 | I | ATC | 1.6 |
| 72 | I | ATT | 9.9 | 60.8 | 3.7 | 1.2 | 70.5 | 21.8 | 46.1 | 78.6 | I | ATA | 3 |
| 72 | V | GTT | 5 | 33.9 | 3.7 | 3.7 | 22 | 70.6 | 43.8 | 14.3 | I | ATT | 1 |
| 72 | V | GTT | 5 | 33.9 | 3.7 | 3.7 | 22 | 70.6 | 43.8 | 14.3 | I | ATC | 4.4 |
| 72 | V | GTT | 5 | 33.9 | 3.7 | 3.7 | 22 | 70.6 | 43.8 | 14.3 | I | ATA | 6.5 |
| 72 | V | GTC | 4.4 | 1 | 29.8 | 77.8 | 1.8 | 1.8 | 7 | 3.1 | I | ATT | 2.9 |
| 72 | V | GTC | 4.4 | 1 | 29.8 | 77.8 | 1.8 | 1.8 | 7 | 3.1 | I | ATC | 1 |
| 72 | V | GTC | 4.4 | 1 | 29.8 | 77.8 | 1.8 | 1.8 | 7 | 3.1 | I | ATA | 5 |
| 72 | I | ATC | 0.5 | 2.3 | 62 | 10.8 | 1 | 0 | 1.9 | 1.6 | I | ATT | 2.6 |
| 72 | I | ATC | 0.5 | 2.3 | 62 | 10.8 | 1 | 0 | 1.9 | 1.6 | I | ATC | 0 |
| 72 | I | ATC | 0.5 | 2.3 | 62 | 10.8 | 1 | 0 | 1.9 | 1.6 | I | ATA | 1.2 |
| 72 | V | GTA | 57.5 | 0.1 | 0.1 | 4.4 | 1 | 2.9 | 0 | 0 | I | ATT | 7.4 |
| 72 | V | GTA | 57.5 | 0.1 | 0.1 | 4.4 | 1 | 2.9 | 0 | 0 | I | ATC | 9.2 |
| 72 | V | GTA | 57.5 | 0.1 | 0.1 | 4.4 | 1 | 2.9 | 0 | 0 | I | ATA | 1 |
| 72 | V | GTG | 9.3 | 0.9 | 0 | 0.5 | 0.5 | 1.8 | 0 | 0 | I | ATT | 21.2 |
| 72 | V | GTG | 9.3 | 0.9 | 0 | 0.5 | 0.5 | 1.8 | 0 | 0 | I | ATC | 3.9 |

| | | | | | | | | | | | | | |
|---|---|---|---|---|---|---|---|---|---|---|---|---|---|
| 72 | V | GTG | 9.3 | 0.9 | 0 | 0.5 | 0.5 | 1.8 | 0 | 0 | I | ATA | 2 |
| 72 | I | ATA | 12.7 | 0.1 | 0.2 | 0.8 | 2.6 | 0 | 0.4 | 1.9 | I | ATT | 7.4 |
| 72 | I | ATA | 12.7 | 0.1 | 0.2 | 0.8 | 2.6 | 0 | 0.4 | 1.9 | I | ATC | 1.8 |
| 72 | I | ATA | 12.7 | 0.1 | 0.2 | 0.8 | 2.6 | 0 | 0.4 | 1.9 | I | ATA | 0 |
| 74 | L | CTG | 46.6 | 86.3 | 78.6 | 90.8 | 58.7 | 82.6 | 86.1 | 74.4 | F | TTT | 22.8 |
| 74 | L | CTG | 46.6 | 86.3 | 78.6 | 90.8 | 58.7 | 82.6 | 86.1 | 74.4 | F | TTC | 5.5 |
| 74 | L | CTA | 15.8 | 6.6 | 9.8 | 2.9 | 4.2 | 3.8 | 6.6 | 5.9 | F | TTT | 9 |
| 74 | L | CTA | 15.8 | 6.6 | 9.8 | 2.9 | 4.2 | 3.8 | 6.6 | 5.9 | F | TTC | 10.8 |
| 74 | I | ATA | 22.4 | 3.5 | 5 | 2.2 | 18.4 | 3.8 | 3.3 | 10.3 | F | TTT | 13.8 |
| 74 | I | ATA | 22.4 | 3.5 | 5 | 2.2 | 18.4 | 3.8 | 3.3 | 10.3 | F | TTC | 15.6 |
| 74 | L | TTG | 11.2 | 2.2 | 4.9 | 2.1 | 4 | 7.1 | 2.7 | 4.1 | F | TTT | 13.5 |
| 74 | L | TTG | 11.2 | 2.2 | 4.9 | 2.1 | 4 | 7.1 | 2.7 | 4.1 | F | TTC | 5.8 |
| 74 | M | ATG | 3.1 | 0.5 | 0.5 | 1.1 | 10.2 | 1.8 | 0.8 | 3.8 | F | TTT | 27.6 |
| 74 | M | ATG | 3.1 | 0.5 | 0.5 | 1.1 | 10.2 | 1.8 | 0.8 | 3.8 | F | TTC | 10.3 |
| 74 | L | TTA | 0.2 | 0.5 | 0.6 | 0.1 | 4.5 | 0.3 | 0 | 1.2 | F | TTT | 7.4 |
| 74 | L | TTA | 0.2 | 0.5 | 0.6 | 0.1 | 4.5 | 0.3 | 0 | 1.2 | F | TTC | 1.8 |
| 74 | L | CTG | 46.6 | 86.3 | 78.6 | 90.8 | 58.7 | 82.6 | 86.1 | 74.4 | I | ATT | 21.4 |
| 74 | L | CTG | 46.6 | 86.3 | 78.6 | 90.8 | 58.7 | 82.6 | 86.1 | 74.4 | I | ATC | 4.1 |
| 74 | L | CTG | 46.6 | 86.3 | 78.6 | 90.8 | 58.7 | 82.6 | 86.1 | 74.4 | I | ATA | 2.2 |
| 74 | L | CTA | 15.8 | 6.6 | 9.8 | 2.9 | 4.2 | 3.8 | 6.6 | 5.9 | I | ATT | 7.6 |
| 74 | L | CTA | 15.8 | 6.6 | 9.8 | 2.9 | 4.2 | 3.8 | 6.6 | 5.9 | I | ATC | 9.4 |
| 74 | L | CTA | 15.8 | 6.6 | 9.8 | 2.9 | 4.2 | 3.8 | 6.6 | 5.9 | I | ATA | 1.2 |
| 74 | I | ATA | 22.4 | 3.5 | 5 | 2.2 | 18.4 | 3.8 | 3.3 | 10.3 | I | ATT | 7.4 |
| 74 | I | ATA | 22.4 | 3.5 | 5 | 2.2 | 18.4 | 3.8 | 3.3 | 10.3 | I | ATC | 1.8 |
| 74 | I | ATA | 22.4 | 3.5 | 5 | 2.2 | 18.4 | 3.8 | 3.3 | 10.3 | I | ATA | 0 |
| 74 | L | TTG | 11.2 | 2.2 | 4.9 | 2.1 | 4 | 7.1 | 2.7 | 4.1 | I | ATT | 23.2 |
| 74 | L | TTG | 11.2 | 2.2 | 4.9 | 2.1 | 4 | 7.1 | 2.7 | 4.1 | I | ATC | 5.9 |
| 74 | L | TTG | 11.2 | 2.2 | 4.9 | 2.1 | 4 | 7.1 | 2.7 | 4.1 | I | ATA | 4 |
| 74 | M | ATG | 3.1 | 0.5 | 0.5 | 1.1 | 10.2 | 1.8 | 0.8 | 3.8 | I | ATT | 13.5 |
| 74 | M | ATG | 3.1 | 0.5 | 0.5 | 1.1 | 10.2 | 1.8 | 0.8 | 3.8 | I | ATC | 5.8 |
| 74 | M | ATG | 3.1 | 0.5 | 0.5 | 1.1 | 10.2 | 1.8 | 0.8 | 3.8 | I | ATA | 1 |
| 74 | L | TTA | 0.2 | 0.5 | 0.6 | 0.1 | 4.5 | 0.3 | 0 | 1.2 | I | ATT | 9.4 |
| 74 | L | TTA | 0.2 | 0.5 | 0.6 | 0.1 | 4.5 | 0.3 | 0 | 1.2 | I | ATC | 11.2 |
| 74 | L | TTA | 0.2 | 0.5 | 0.6 | 0.1 | 4.5 | 0.3 | 0 | 1.2 | I | ATA | 3 |

| | | | | | | | | | | | | | |
|---|---|---|---|---|---|---|---|---|---|---|---|---|---|
| 74 | L | CTG | 46.6 | 86.3 | 78.6 | 90.8 | 58.7 | 82.6 | 86.1 | 74.4 | M | ATG | 1.2 |
| 74 | L | CTA | 15.8 | 6.6 | 9.8 | 2.9 | 4.2 | 3.8 | 6.6 | 5.9 | M | ATG | 3.7 |
| 74 | I | ATA | 22.4 | 3.5 | 5 | 2.2 | 18.4 | 3.8 | 3.3 | 10.3 | M | ATG | 1 |
| 74 | L | TTG | 11.2 | 2.2 | 4.9 | 2.1 | 4 | 7.1 | 2.7 | 4.1 | M | ATG | 3 |
| 74 | M | ATG | 3.1 | 0.5 | 0.5 | 1.1 | 10.2 | 1.8 | 0.8 | 3.8 | M | ATG | 0 |
| 74 | L | TTA | 0.2 | 0.5 | 0.6 | 0.1 | 4.5 | 0.3 | 0 | 1.2 | M | ATG | 5.5 |
| 75 | V | GTA | 98 | 91.1 | 98.3 | 97.9 | 96.2 | 93.8 | 98.2 | 94.7 | I | ATT | 7.4 |
| 75 | V | GTA | 98 | 91.1 | 98.3 | 97.9 | 96.2 | 93.8 | 98.2 | 94.7 | I | ATC | 9.2 |
| 75 | V | GTA | 98 | 91.1 | 98.3 | 97.9 | 96.2 | 93.8 | 98.2 | 94.7 | I | ATA | 1 |
| 75 | V | GTG | 2 | 6.3 | 1.3 | 1.8 | 3.7 | 5 | 1.8 | 5.3 | I | ATT | 21.2 |
| 75 | V | GTG | 2 | 6.3 | 1.3 | 1.8 | 3.7 | 5 | 1.8 | 5.3 | I | ATC | 3.9 |
| 75 | V | GTG | 2 | 6.3 | 1.3 | 1.8 | 3.7 | 5 | 1.8 | 5.3 | I | ATA | 2 |
| 75 | V | GTT | 0 | 2.2 | 0.2 | 0 | 0.1 | 0.6 | 0 | 0 | I | ATT | 1 |
| 75 | V | GTT | 0 | 2.2 | 0.2 | 0 | 0.1 | 0.6 | 0 | 0 | I | ATC | 4.4 |
| 75 | V | GTT | 0 | 2.2 | 0.2 | 0 | 0.1 | 0.6 | 0 | 0 | I | ATA | 6.5 |
| 92 | E | GAA | 97 | 29.6 | 96.5 | 97.6 | 97.8 | 96.2 | 94.6 | 96.3 | A | GCT | 8.2 |
| 92 | E | GAA | 97 | 29.6 | 96.5 | 97.6 | 97.8 | 96.2 | 94.6 | 96.3 | A | GCC | 10 |
| 92 | E | GAA | 97 | 29.6 | 96.5 | 97.6 | 97.8 | 96.2 | 94.6 | 96.3 | A | GCA | 1.8 |
| 92 | E | GAA | 97 | 29.6 | 96.5 | 97.6 | 97.8 | 96.2 | 94.6 | 96.3 | A | GCG | 4.3 |
| 92 | E | GAG | 3 | 70.3 | 3.4 | 2.2 | 2.2 | 3.8 | 5.4 | 3.1 | A | GCT | 22 |
| 92 | E | GAG | 3 | 70.3 | 3.4 | 2.2 | 2.2 | 3.8 | 5.4 | 3.1 | A | GCC | 4.7 |
| 92 | E | GAG | 3 | 70.3 | 3.4 | 2.2 | 2.2 | 3.8 | 5.4 | 3.1 | A | GCA | 2.8 |
| 92 | E | GAG | 3 | 70.3 | 3.4 | 2.2 | 2.2 | 3.8 | 5.4 | 3.1 | A | GCG | 1.8 |
| 92 | E | GAA | 97 | 29.6 | 96.5 | 97.6 | 97.8 | 96.2 | 94.6 | 96.3 | G | GGT | 7.4 |
| 92 | E | GAA | 97 | 29.6 | 96.5 | 97.6 | 97.8 | 96.2 | 94.6 | 96.3 | G | GGC | 9.2 |
| 92 | E | GAA | 97 | 29.6 | 96.5 | 97.6 | 97.8 | 96.2 | 94.6 | 96.3 | G | GGA | 1 |
| 92 | E | GAA | 97 | 29.6 | 96.5 | 97.6 | 97.8 | 96.2 | 94.6 | 96.3 | G | GGG | 3.5 |
| 92 | E | GAG | 3 | 70.3 | 3.4 | 2.2 | 2.2 | 3.8 | 5.4 | 3.1 | G | GGT | 21.2 |
| 92 | E | GAG | 3 | 70.3 | 3.4 | 2.2 | 2.2 | 3.8 | 5.4 | 3.1 | G | GGC | 3.9 |
| 92 | E | GAG | 3 | 70.3 | 3.4 | 2.2 | 2.2 | 3.8 | 5.4 | 3.1 | G | GGA | 2 |
| 92 | E | GAG | 3 | 70.3 | 3.4 | 2.2 | 2.2 | 3.8 | 5.4 | 3.1 | G | GGG | 1 |
| 92 | E | GAA | 97 | 29.6 | 96.5 | 97.6 | 97.8 | 96.2 | 94.6 | 96.3 | Q | CAA | 5.8 |
| 92 | E | GAA | 97 | 29.6 | 96.5 | 97.6 | 97.8 | 96.2 | 94.6 | 96.3 | Q | CAG | 8.3 |
| 92 | E | GAG | 3 | 70.3 | 3.4 | 2.2 | 2.2 | 3.8 | 5.4 | 3.1 | Q | CAA | 6.8 |

| | | | | | | | | | | | | | |
|---|---|---|---|---|---|---|---|---|---|---|---|---|---|
| 92 | E | GAG | 3 | 70.3 | 3.4 | 2.2 | 2.2 | 3.8 | 5.4 | 3.1 | Q | CAG | 5.8 |
| 92 | E | GAA | 97 | 29.6 | 96.5 | 97.6 | 97.8 | 96.2 | 94.6 | 96.3 | V | GTT | 13.8 |
| 92 | E | GAA | 97 | 29.6 | 96.5 | 97.6 | 97.8 | 96.2 | 94.6 | 96.3 | V | GTC | 15.6 |
| 92 | E | GAA | 97 | 29.6 | 96.5 | 97.6 | 97.8 | 96.2 | 94.6 | 96.3 | V | GTA | 7.4 |
| 92 | E | GAA | 97 | 29.6 | 96.5 | 97.6 | 97.8 | 96.2 | 94.6 | 96.3 | V | GTG | 9.9 |
| 92 | E | GAG | 3 | 70.3 | 3.4 | 2.2 | 2.2 | 3.8 | 5.4 | 3.1 | V | GTT | 27.6 |
| 92 | E | GAG | 3 | 70.3 | 3.4 | 2.2 | 2.2 | 3.8 | 5.4 | 3.1 | V | GTC | 10.3 |
| 92 | E | GAG | 3 | 70.3 | 3.4 | 2.2 | 2.2 | 3.8 | 5.4 | 3.1 | V | GTA | 8.4 |
| 92 | E | GAG | 3 | 70.3 | 3.4 | 2.2 | 2.2 | 3.8 | 5.4 | 3.1 | V | GTG | 7.4 |
| 95 | Q | CAG | 66.7 | 74.3 | 9.7 | 80 | 82.8 | 87.9 | 48.6 | 89.4 | K | AAA | 2.2 |
| 95 | Q | CAG | 66.7 | 74.3 | 9.7 | 80 | 82.8 | 87.9 | 48.6 | 89.4 | K | AAG | 1.2 |
| 95 | Q | CAA | 32.1 | 25.2 | 88.5 | 15 | 14.5 | 11.5 | 51 | 9.9 | K | AAA | 1.2 |
| 95 | Q | CAA | 32.1 | 25.2 | 88.5 | 15 | 14.5 | 11.5 | 51 | 9.9 | K | AAG | 3.7 |
| 95 | H | CAT | 0 | 0.1 | 0.2 | 2.8 | 0.3 | 0 | 0 | 0 | K | AAA | 6.7 |
| 95 | H | CAT | 0 | 0.1 | 0.2 | 2.8 | 0.3 | 0 | 0 | 0 | K | AAG | 43.4 |
| 97 | T | ACA | 93.4 | 96.4 | 98.2 | 98.8 | 93.6 | 90.9 | 92.6 | 95.3 | A | GCT | 7.4 |
| 97 | T | ACA | 93.4 | 96.4 | 98.2 | 98.8 | 93.6 | 90.9 | 92.6 | 95.3 | A | GCC | 9.2 |
| 97 | T | ACA | 93.4 | 96.4 | 98.2 | 98.8 | 93.6 | 90.9 | 92.6 | 95.3 | A | GCA | 1 |
| 97 | T | ACA | 93.4 | 96.4 | 98.2 | 98.8 | 93.6 | 90.9 | 92.6 | 95.3 | A | GCG | 3.5 |
| 97 | A | GCA | 5.1 | 0.5 | 1.1 | 0.7 | 5.5 | 5 | 4.7 | 4.7 | A | GCT | 7.4 |
| 97 | A | GCA | 5.1 | 0.5 | 1.1 | 0.7 | 5.5 | 5 | 4.7 | 4.7 | A | GCC | 1.8 |
| 97 | A | GCA | 5.1 | 0.5 | 1.1 | 0.7 | 5.5 | 5 | 4.7 | 4.7 | A | GCA | 0 |
| 97 | A | GCA | 5.1 | 0.5 | 1.1 | 0.7 | 5.5 | 5 | 4.7 | 4.7 | A | GCG | 1 |
| 97 | T | ACC | 0.4 | 2.5 | 0 | 0.1 | 0.3 | 2.4 | 0.8 | 0 | A | GCT | 2.9 |
| 97 | T | ACC | 0.4 | 2.5 | 0 | 0.1 | 0.3 | 2.4 | 0.8 | 0 | A | GCC | 1 |
| 97 | T | ACC | 0.4 | 2.5 | 0 | 0.1 | 0.3 | 2.4 | 0.8 | 0 | A | GCA | 5 |
| 97 | T | ACC | 0.4 | 2.5 | 0 | 0.1 | 0.3 | 2.4 | 0.8 | 0 | A | GCG | 4.4 |
| 97 | T | ACG | 0.6 | 0.3 | 0.2 | 0.1 | 0.5 | 0 | 1.2 | 0 | A | GCT | 21.2 |
| 97 | T | ACG | 0.6 | 0.3 | 0.2 | 0.1 | 0.5 | 0 | 1.2 | 0 | A | GCC | 3.9 |
| 97 | T | ACG | 0.6 | 0.3 | 0.2 | 0.1 | 0.5 | 0 | 1.2 | 0 | A | GCA | 2 |
| 97 | T | ACG | 0.6 | 0.3 | 0.2 | 0.1 | 0.5 | 0 | 1.2 | 0 | A | GCG | 1 |
| 97 | A | GCG | 0 | 0 | 0 | 0 | 0 | 1.2 | 0 | 0 | A | GCT | 13.5 |
| 97 | A | GCG | 0 | 0 | 0 | 0 | 0 | 1.2 | 0 | 0 | A | GCC | 5.8 |
| 97 | A | GCG | 0 | 0 | 0 | 0 | 0 | 1.2 | 0 | 0 | A | GCA | 1 |

| | | | | | | | | | | | | |
|---|---|---|---|---|---|---|---|---|---|---|---|---|
| 97 | A | GCG | 0 | 0 | 0 | 0 | 0 | 1.2 | 0 | 0 | A | GCG | 0 |
| 101 | I | ATA | 25.2 | 1.8 | 93.3 | 9 | 88.5 | 2.4 | 83.3 | 76.4 | I | ATT | 7.4 |
| 101 | I | ATA | 25.2 | 1.8 | 93.3 | 9 | 88.5 | 2.4 | 83.3 | 76.4 | I | ATC | 1.8 |
| 101 | I | ATA | 25.2 | 1.8 | 93.3 | 9 | 88.5 | 2.4 | 83.3 | 76.4 | I | ATA | 0 |
| 101 | L | CTC | 0.7 | 44.7 | 0.2 | 0.1 | 0.6 | 58.8 | 1.2 | 0.6 | I | ATT | 3.1 |
| 101 | L | CTC | 0.7 | 44.7 | 0.2 | 0.1 | 0.6 | 58.8 | 1.2 | 0.6 | I | ATC | 1.2 |
| 101 | L | CTC | 0.7 | 44.7 | 0.2 | 0.1 | 0.6 | 58.8 | 1.2 | 0.6 | I | ATA | 5.2 |
| 101 | I | ATC | 0.2 | 38.6 | 1 | 0.3 | 0.8 | 20.9 | 3.7 | 2.8 | I | ATT | 2.6 |
| 101 | I | ATC | 0.2 | 38.6 | 1 | 0.3 | 0.8 | 20.9 | 3.7 | 2.8 | I | ATC | 0 |
| 101 | I | ATC | 0.2 | 38.6 | 1 | 0.3 | 0.8 | 20.9 | 3.7 | 2.8 | I | ATA | 1.2 |
| 101 | L | CTG | 54.8 | 1.8 | 0.3 | 83.2 | 1.7 | 2.1 | 0 | 2.5 | I | ATT | 21.4 |
| 101 | L | CTG | 54.8 | 1.8 | 0.3 | 83.2 | 1.7 | 2.1 | 0 | 2.5 | I | ATC | 4.1 |
| 101 | L | CTG | 54.8 | 1.8 | 0.3 | 83.2 | 1.7 | 2.1 | 0 | 2.5 | I | ATA | 2.2 |
| 101 | L | CTT | 0 | 6.2 | 0 | 0 | 0.2 | 10.3 | 0 | 0 | I | ATT | 1.2 |
| 101 | L | CTT | 0 | 6.2 | 0 | 0 | 0.2 | 10.3 | 0 | 0 | I | ATC | 4.6 |
| 101 | L | CTT | 0 | 6.2 | 0 | 0 | 0.2 | 10.3 | 0 | 0 | I | ATA | 6.7 |
| 101 | L | CTA | 16.6 | 1 | 2.9 | 3.4 | 6.3 | 1.5 | 7.4 | 12.7 | I | ATT | 7.6 |
| 101 | L | CTA | 16.6 | 1 | 2.9 | 3.4 | 6.3 | 1.5 | 7.4 | 12.7 | I | ATC | 9.4 |
| 101 | L | CTA | 16.6 | 1 | 2.9 | 3.4 | 6.3 | 1.5 | 7.4 | 12.7 | I | ATA | 1.2 |
| 101 | I | ATT | 0.2 | 4.4 | 0.3 | 0.3 | 0 | 3.2 | 0.8 | 0 | I | ATT | 0 |
| 101 | I | ATT | 0.2 | 4.4 | 0.3 | 0.3 | 0 | 3.2 | 0.8 | 0 | I | ATC | 1.6 |
| 101 | I | ATT | 0.2 | 4.4 | 0.3 | 0.3 | 0 | 3.2 | 0.8 | 0 | I | ATA | 3 |
| 101 | V | GTC | 0 | 1.2 | 0 | 0 | 0 | 0 | 0 | 0 | I | ATT | 2.9 |
| 101 | V | GTC | 0 | 1.2 | 0 | 0 | 0 | 0 | 0 | 0 | I | ATC | 1 |
| 101 | V | GTC | 0 | 1.2 | 0 | 0 | 0 | 0 | 0 | 0 | I | ATA | 5 |
| 101 | V | GTG | 0.1 | 0 | 0.1 | 2.4 | 0 | 0 | 0 | 0.6 | I | ATT | 21.2 |
| 101 | V | GTG | 0.1 | 0 | 0.1 | 2.4 | 0 | 0 | 0 | 0.6 | I | ATC | 3.9 |
| 101 | V | GTG | 0.1 | 0 | 0.1 | 2.4 | 0 | 0 | 0 | 0.6 | I | ATA | 2 |
| 101 | L | TTA | 1 | 0 | 0.8 | 0.2 | 1.4 | 0 | 3.3 | 2.5 | I | ATT | 9.4 |
| 101 | L | TTA | 1 | 0 | 0.8 | 0.2 | 1.4 | 0 | 3.3 | 2.5 | I | ATC | 11.2 |
| 101 | L | TTA | 1 | 0 | 0.8 | 0.2 | 1.4 | 0 | 3.3 | 2.5 | I | ATA | 3 |
| 101 | V | GTA | 0.4 | 0.1 | 1 | 0 | 0.6 | 0.6 | 0.4 | 1.9 | I | ATT | 7.4 |
| 101 | V | GTA | 0.4 | 0.1 | 1 | 0 | 0.6 | 0.6 | 0.4 | 1.9 | I | ATC | 9.2 |
| 101 | V | GTA | 0.4 | 0.1 | 1 | 0 | 0.6 | 0.6 | 0.4 | 1.9 | I | ATA | 1 |

| | | | | | | | | | | | | | |
|---|---|---|---|---|---|---|---|---|---|---|---|---|---|
| 114 | H | CAT | 11.3 | 93.7 | 90.2 | 5.4 | 8 | 92.9 | 97.1 | 72.4 | Y | TAT | 2.6 |
| 114 | H | CAT | 11.3 | 93.7 | 90.2 | 5.4 | 8 | 92.9 | 97.1 | 72.4 | Y | TAC | 6 |
| 114 | H | CAC | 88.4 | 6.3 | 9.8 | 94.6 | 91.9 | 7.1 | 2.9 | 27.3 | Y | TAT | 4.5 |
| 114 | H | CAC | 88.4 | 6.3 | 9.8 | 94.6 | 91.9 | 7.1 | 2.9 | 27.3 | Y | TAC | 2.6 |
| 118 | G | GGC | 78.3 | 90.1 | 65.4 | 6 | 91.9 | 92.6 | 92.6 | 91.9 | R | CGT | 7.7 |
| 118 | G | GGC | 78.3 | 90.1 | 65.4 | 6 | 91.9 | 92.6 | 92.6 | 91.9 | R | CGC | 5.8 |
| 118 | G | GGC | 78.3 | 90.1 | 65.4 | 6 | 91.9 | 92.6 | 92.6 | 91.9 | R | CGA | 9.8 |
| 118 | G | GGC | 78.3 | 90.1 | 65.4 | 6 | 91.9 | 92.6 | 92.6 | 91.9 | R | CGG | 9.2 |
| 118 | G | GGC | 78.3 | 90.1 | 65.4 | 6 | 91.9 | 92.6 | 92.6 | 91.9 | R | AGA | 5 |
| 118 | G | GGC | 78.3 | 90.1 | 65.4 | 6 | 91.9 | 92.6 | 92.6 | 91.9 | R | AGG | 4.4 |
| 118 | G | GGT | 16.5 | 8 | 32.2 | 89.7 | 6.5 | 4.7 | 7.2 | 7.5 | R | CGT | 5.8 |
| 118 | G | GGT | 16.5 | 8 | 32.2 | 89.7 | 6.5 | 4.7 | 7.2 | 7.5 | R | CGC | 9.2 |
| 118 | G | GGT | 16.5 | 8 | 32.2 | 89.7 | 6.5 | 4.7 | 7.2 | 7.5 | R | CGA | 11.3 |
| 118 | G | GGT | 16.5 | 8 | 32.2 | 89.7 | 6.5 | 4.7 | 7.2 | 7.5 | R | CGG | 48 |
| 118 | G | GGT | 16.5 | 8 | 32.2 | 89.7 | 6.5 | 4.7 | 7.2 | 7.5 | R | AGA | 6.5 |
| 118 | G | GGT | 16.5 | 8 | 32.2 | 89.7 | 6.5 | 4.7 | 7.2 | 7.5 | R | AGG | 43.2 |
| 118 | G | GGA | 2.7 | 1.5 | 1.6 | 2.2 | 1.3 | 1.5 | 0 | 0.6 | R | CGT | 12.2 |
| 118 | G | GGA | 2.7 | 1.5 | 1.6 | 2.2 | 1.3 | 1.5 | 0 | 0.6 | R | CGC | 14 |
| 118 | G | GGA | 2.7 | 1.5 | 1.6 | 2.2 | 1.3 | 1.5 | 0 | 0.6 | R | CGA | 5.8 |
| 118 | G | GGA | 2.7 | 1.5 | 1.6 | 2.2 | 1.3 | 1.5 | 0 | 0.6 | R | CGG | 8.3 |
| 118 | G | GGA | 2.7 | 1.5 | 1.6 | 2.2 | 1.3 | 1.5 | 0 | 0.6 | R | AGA | 1 |
| 118 | G | GGA | 2.7 | 1.5 | 1.6 | 2.2 | 1.3 | 1.5 | 0 | 0.6 | R | AGG | 3.5 |
| 118 | G | GGG | 2.6 | 0.3 | 0.7 | 2.2 | 0.3 | 1.2 | 0.2 | 0 | R | CGT | 26 |
| 118 | G | GGG | 2.6 | 0.3 | 0.7 | 2.2 | 0.3 | 1.2 | 0.2 | 0 | R | CGC | 8.7 |
| 118 | G | GGG | 2.6 | 0.3 | 0.7 | 2.2 | 0.3 | 1.2 | 0.2 | 0 | R | CGA | 6.8 |
| 118 | G | GGG | 2.6 | 0.3 | 0.7 | 2.2 | 0.3 | 1.2 | 0.2 | 0 | R | CGG | 5.8 |
| 118 | G | GGG | 2.6 | 0.3 | 0.7 | 2.2 | 0.3 | 1.2 | 0.2 | 0 | R | AGA | 2 |
| 118 | G | GGG | 2.6 | 0.3 | 0.7 | 2.2 | 0.3 | 1.2 | 0.2 | 0 | R | AGG | 1 |
| 119 | S | AGC | 63.6 | 62.7 | 7.5 | 94.7 | 82.5 | 74.9 | 32.3 | 78.3 | R | CGT | 3.7 |
| 119 | S | AGC | 63.6 | 62.7 | 7.5 | 94.7 | 82.5 | 74.9 | 32.3 | 78.3 | R | CGC | 1.8 |
| 119 | S | AGC | 63.6 | 62.7 | 7.5 | 94.7 | 82.5 | 74.9 | 32.3 | 78.3 | R | CGA | 5.8 |
| 119 | S | AGC | 63.6 | 62.7 | 7.5 | 94.7 | 82.5 | 74.9 | 32.3 | 78.3 | R | CGG | 5.2 |
| 119 | S | AGC | 63.6 | 62.7 | 7.5 | 94.7 | 82.5 | 74.9 | 32.3 | 78.3 | R | AGA | 1.2 |
| 119 | S | AGC | 63.6 | 62.7 | 7.5 | 94.7 | 82.5 | 74.9 | 32.3 | 78.3 | R | AGG | 3.6 |

| | | | | | | | | | | | | |
|---|---|---|---|---|---|---|---|---|---|---|---|---|
| 119 | S | AGT | 3.8 | 4.8 | 81.6 | 3.3 | 4.4 | 6.8 | 1.6 | 4.3 | R | CGT | 1.8 |
| 119 | S | AGT | 3.8 | 4.8 | 81.6 | 3.3 | 4.4 | 6.8 | 1.6 | 4.3 | R | CGC | 5.2 |
| 119 | S | AGT | 3.8 | 4.8 | 81.6 | 3.3 | 4.4 | 6.8 | 1.6 | 4.3 | R | CGA | 7.3 |
| 119 | S | AGT | 3.8 | 4.8 | 81.6 | 3.3 | 4.4 | 6.8 | 1.6 | 4.3 | R | CGG | 44 |
| 119 | S | AGT | 3.8 | 4.8 | 81.6 | 3.3 | 4.4 | 6.8 | 1.6 | 4.3 | R | AGA | 3 |
| 119 | S | AGT | 3.8 | 4.8 | 81.6 | 3.3 | 4.4 | 6.8 | 1.6 | 4.3 | R | AGG | 5.3 |
| 119 | P | CCC | 27.5 | 12.2 | 0.6 | 1 | 4.8 | 3.8 | 28 | 7.5 | R | CGT | 5.5 |
| 119 | P | CCC | 27.5 | 12.2 | 0.6 | 1 | 4.8 | 3.8 | 28 | 7.5 | R | CGC | 3.6 |
| 119 | P | CCC | 27.5 | 12.2 | 0.6 | 1 | 4.8 | 3.8 | 28 | 7.5 | R | CGA | 7.6 |
| 119 | P | CCC | 27.5 | 12.2 | 0.6 | 1 | 4.8 | 3.8 | 28 | 7.5 | R | CGG | 7 |
| 119 | P | CCC | 27.5 | 12.2 | 0.6 | 1 | 4.8 | 3.8 | 28 | 7.5 | R | AGA | 5.9 |
| 119 | P | CCC | 27.5 | 12.2 | 0.6 | 1 | 4.8 | 3.8 | 28 | 7.5 | R | AGG | 25.6 |
| 119 | T | ACC | 1.1 | 5.1 | 0.1 | 0.4 | 3 | 1.8 | 30.7 | 2.5 | R | CGT | 6.2 |
| 119 | T | ACC | 1.1 | 5.1 | 0.1 | 0.4 | 3 | 1.8 | 30.7 | 2.5 | R | CGC | 5.2 |
| 119 | T | ACC | 1.1 | 5.1 | 0.1 | 0.4 | 3 | 1.8 | 30.7 | 2.5 | R | CGA | 6.5 |
| 119 | T | ACC | 1.1 | 5.1 | 0.1 | 0.4 | 3 | 1.8 | 30.7 | 2.5 | R | CGG | 26.2 |
| 119 | T | ACC | 1.1 | 5.1 | 0.1 | 0.4 | 3 | 1.8 | 30.7 | 2.5 | R | AGA | 7.6 |
| 119 | T | ACC | 1.1 | 5.1 | 0.1 | 0.4 | 3 | 1.8 | 30.7 | 2.5 | R | AGG | 7 |
| 119 | G | GGC | 1.2 | 5.6 | 0 | 0.2 | 1.8 | 4.7 | 0.4 | 1.9 | R | CGT | 7.7 |
| 119 | G | GGC | 1.2 | 5.6 | 0 | 0.2 | 1.8 | 4.7 | 0.4 | 1.9 | R | CGC | 5.8 |
| 119 | G | GGC | 1.2 | 5.6 | 0 | 0.2 | 1.8 | 4.7 | 0.4 | 1.9 | R | CGA | 9.8 |
| 119 | G | GGC | 1.2 | 5.6 | 0 | 0.2 | 1.8 | 4.7 | 0.4 | 1.9 | R | CGG | 9.2 |
| 119 | G | GGC | 1.2 | 5.6 | 0 | 0.2 | 1.8 | 4.7 | 0.4 | 1.9 | R | AGA | 5 |
| 119 | G | GGC | 1.2 | 5.6 | 0 | 0.2 | 1.8 | 4.7 | 0.4 | 1.9 | R | AGG | 4.4 |
| 119 | R | AGA | 0.2 | 5.3 | 0.7 | 0 | 1.4 | 3.6 | 0.4 | 1.9 | R | CGT | 8.2 |
| 119 | R | AGA | 0.2 | 5.3 | 0.7 | 0 | 1.4 | 3.6 | 0.4 | 1.9 | R | CGC | 10 |
| 119 | R | AGA | 0.2 | 5.3 | 0.7 | 0 | 1.4 | 3.6 | 0.4 | 1.9 | R | CGA | 1.8 |
| 119 | R | AGA | 0.2 | 5.3 | 0.7 | 0 | 1.4 | 3.6 | 0.4 | 1.9 | R | CGG | 4.3 |
| 119 | R | AGA | 0.2 | 5.3 | 0.7 | 0 | 1.4 | 3.6 | 0.4 | 1.9 | R | AGA | 0 |
| 119 | R | AGA | 0.2 | 5.3 | 0.7 | 0 | 1.4 | 3.6 | 0.4 | 1.9 | R | AGG | 1 |
| 119 | P | CCT | 0.6 | 1 | 4.7 | 0 | 0.2 | 0.3 | 1.2 | 2.5 | R | CGT | 3.6 |
| 119 | P | CCT | 0.6 | 1 | 4.7 | 0 | 0.2 | 0.3 | 1.2 | 2.5 | R | CGC | 7 |
| 119 | P | CCT | 0.6 | 1 | 4.7 | 0 | 0.2 | 0.3 | 1.2 | 2.5 | R | CGA | 9.1 |
| 119 | P | CCT | 0.6 | 1 | 4.7 | 0 | 0.2 | 0.3 | 1.2 | 2.5 | R | CGG | 45.8 |

| | | | | | | | | | | | | |
|---|---|---|---|---|---|---|---|---|---|---|---|---|
| 119 | P | CCT | 0.6 | 1 | 4.7 | 0 | 0.2 | 0.3 | 1.2 | 2.5 | R | AGA | 8.5 |
| 119 | P | CCT | 0.6 | 1 | 4.7 | 0 | 0.2 | 0.3 | 1.2 | 2.5 | R | AGG | 55.8 |
| 119 | T | ACT | 0 | 0.4 | 3.3 | 0 | 0.1 | 0.6 | 2.9 | 0 | R | CGT | 5.2 |
| 119 | T | ACT | 0 | 0.4 | 3.3 | 0 | 0.1 | 0.6 | 2.9 | 0 | R | CGC | 7.2 |
| 119 | T | ACT | 0 | 0.4 | 3.3 | 0 | 0.1 | 0.6 | 2.9 | 0 | R | CGA | 9.1 |
| 119 | T | ACT | 0 | 0.4 | 3.3 | 0 | 0.1 | 0.6 | 2.9 | 0 | R | CGG | 56.4 |
| 119 | T | ACT | 0 | 0.4 | 3.3 | 0 | 0.1 | 0.6 | 2.9 | 0 | R | AGA | 9.1 |
| 119 | T | ACT | 0 | 0.4 | 3.3 | 0 | 0.1 | 0.6 | 2.9 | 0 | R | AGG | 45.8 |
| 119 | R | AGG | 0 | 0.2 | 0.2 | 0.1 | 0.6 | 0.6 | 0 | 1.2 | R | CGT | 22 |
| 119 | R | AGG | 0 | 0.2 | 0.2 | 0.1 | 0.6 | 0.6 | 0 | 1.2 | R | CGC | 4.7 |
| 119 | R | AGG | 0 | 0.2 | 0.2 | 0.1 | 0.6 | 0.6 | 0 | 1.2 | R | CGA | 2.8 |
| 119 | R | AGG | 0 | 0.2 | 0.2 | 0.1 | 0.6 | 0.6 | 0 | 1.2 | R | CGG | 1.8 |
| 119 | R | AGG | 0 | 0.2 | 0.2 | 0.1 | 0.6 | 0.6 | 0 | 1.2 | R | AGA | 1 |
| 119 | R | AGG | 0 | 0.2 | 0.2 | 0.1 | 0.6 | 0.6 | 0 | 1.2 | R | AGG | 0 |
| 119 | S | TCC | 0.7 | 0.3 | 0 | 0 | 0.3 | 1.2 | 0.4 | 0 | R | CGT | 6 |
| 119 | S | TCC | 0.7 | 0.3 | 0 | 0 | 0.3 | 1.2 | 0.4 | 0 | R | CGC | 5 |
| 119 | S | TCC | 0.7 | 0.3 | 0 | 0 | 0.3 | 1.2 | 0.4 | 0 | R | CGA | 6.3 |
| 119 | S | TCC | 0.7 | 0.3 | 0 | 0 | 0.3 | 1.2 | 0.4 | 0 | R | CGG | 26 |
| 119 | S | TCC | 0.7 | 0.3 | 0 | 0 | 0.3 | 1.2 | 0.4 | 0 | R | AGA | 7.7 |
| 119 | S | TCC | 0.7 | 0.3 | 0 | 0 | 0.3 | 1.2 | 0.4 | 0 | R | AGG | 27.4 |
| 119 | S | AGC | 63.6 | 62.7 | 7.5 | 94.7 | 82.5 | 74.9 | 32.3 | 78.3 | S | TCT | 11.3 |
| 119 | S | AGC | 63.6 | 62.7 | 7.5 | 94.7 | 82.5 | 74.9 | 32.3 | 78.3 | S | TCC | 10.3 |
| 119 | S | AGC | 63.6 | 62.7 | 7.5 | 94.7 | 82.5 | 74.9 | 32.3 | 78.3 | S | TCA | 11.6 |
| 119 | S | AGC | 63.6 | 62.7 | 7.5 | 94.7 | 82.5 | 74.9 | 32.3 | 78.3 | S | TCG | 31.3 |
| 119 | S | AGC | 63.6 | 62.7 | 7.5 | 94.7 | 82.5 | 74.9 | 32.3 | 78.3 | S | AGT | 2.6 |
| 119 | S | AGC | 63.6 | 62.7 | 7.5 | 94.7 | 82.5 | 74.9 | 32.3 | 78.3 | S | AGC | 0 |
| 119 | S | AGT | 3.8 | 4.8 | 81.6 | 3.3 | 4.4 | 6.8 | 1.6 | 4.3 | S | TCT | 10.3 |
| 119 | S | AGT | 3.8 | 4.8 | 81.6 | 3.3 | 4.4 | 6.8 | 1.6 | 4.3 | S | TCC | 12.3 |
| 119 | S | AGT | 3.8 | 4.8 | 81.6 | 3.3 | 4.4 | 6.8 | 1.6 | 4.3 | S | TCA | 14.2 |
| 119 | S | AGT | 3.8 | 4.8 | 81.6 | 3.3 | 4.4 | 6.8 | 1.6 | 4.3 | S | TCG | 61.5 |
| 119 | S | AGT | 3.8 | 4.8 | 81.6 | 3.3 | 4.4 | 6.8 | 1.6 | 4.3 | S | AGT | 0 |
| 119 | S | AGT | 3.8 | 4.8 | 81.6 | 3.3 | 4.4 | 6.8 | 1.6 | 4.3 | S | AGC | 1.6 |
| 119 | P | CCC | 27.5 | 12.2 | 0.6 | 1 | 4.8 | 3.8 | 28 | 7.5 | S | TCT | 4.5 |
| 119 | P | CCC | 27.5 | 12.2 | 0.6 | 1 | 4.8 | 3.8 | 28 | 7.5 | S | TCC | 2.6 |

| | | | | | | | | | | | | |
|---|---|---|---|---|---|---|---|---|---|---|---|---|
| 119 | P | CCC | 27.5 | 12.2 | 0.6 | 1 | 4.8 | 3.8 | 28 | 7.5 | S | TCA | 6.6 |
| 119 | P | CCC | 27.5 | 12.2 | 0.6 | 1 | 4.8 | 3.8 | 28 | 7.5 | S | TCG | 6 |
| 119 | P | CCC | 27.5 | 12.2 | 0.6 | 1 | 4.8 | 3.8 | 28 | 7.5 | S | AGT | 5.6 |
| 119 | P | CCC | 27.5 | 12.2 | 0.6 | 1 | 4.8 | 3.8 | 28 | 7.5 | S | AGC | 4.6 |
| 119 | T | ACC | 1.1 | 5.1 | 0.1 | 0.4 | 3 | 1.8 | 30.7 | 2.5 | S | TCT | 9.3 |
| 119 | T | ACC | 1.1 | 5.1 | 0.1 | 0.4 | 3 | 1.8 | 30.7 | 2.5 | S | TCC | 7.4 |
| 119 | T | ACC | 1.1 | 5.1 | 0.1 | 0.4 | 3 | 1.8 | 30.7 | 2.5 | S | TCA | 11.4 |
| 119 | T | ACC | 1.1 | 5.1 | 0.1 | 0.4 | 3 | 1.8 | 30.7 | 2.5 | S | TCG | 10.8 |
| 119 | T | ACC | 1.1 | 5.1 | 0.1 | 0.4 | 3 | 1.8 | 30.7 | 2.5 | S | AGT | 5.5 |
| 119 | T | ACC | 1.1 | 5.1 | 0.1 | 0.4 | 3 | 1.8 | 30.7 | 2.5 | S | AGC | 3.6 |
| 119 | G | GGC | 1.2 | 5.6 | 0 | 0.2 | 1.8 | 4.7 | 0.4 | 1.9 | S | TCT | 17.4 |
| 119 | G | GGC | 1.2 | 5.6 | 0 | 0.2 | 1.8 | 4.7 | 0.4 | 1.9 | S | TCC | 16.4 |
| 119 | G | GGC | 1.2 | 5.6 | 0 | 0.2 | 1.8 | 4.7 | 0.4 | 1.9 | S | TCA | 17.7 |
| 119 | G | GGC | 1.2 | 5.6 | 0 | 0.2 | 1.8 | 4.7 | 0.4 | 1.9 | S | TCG | 37.4 |
| 119 | G | GGC | 1.2 | 5.6 | 0 | 0.2 | 1.8 | 4.7 | 0.4 | 1.9 | S | AGT | 2.9 |
| 119 | G | GGC | 1.2 | 5.6 | 0 | 0.2 | 1.8 | 4.7 | 0.4 | 1.9 | S | AGC | 1 |
| 119 | R | AGA | 0.2 | 5.3 | 0.7 | 0 | 1.4 | 3.6 | 0.4 | 1.9 | S | TCT | 18.6 |
| 119 | R | AGA | 0.2 | 5.3 | 0.7 | 0 | 1.4 | 3.6 | 0.4 | 1.9 | S | TCC | 16 |
| 119 | R | AGA | 0.2 | 5.3 | 0.7 | 0 | 1.4 | 3.6 | 0.4 | 1.9 | S | TCA | 10.3 |
| 119 | R | AGA | 0.2 | 5.3 | 0.7 | 0 | 1.4 | 3.6 | 0.4 | 1.9 | S | TCG | 13.6 |
| 119 | R | AGA | 0.2 | 5.3 | 0.7 | 0 | 1.4 | 3.6 | 0.4 | 1.9 | S | AGT | 7.4 |
| 119 | R | AGA | 0.2 | 5.3 | 0.7 | 0 | 1.4 | 3.6 | 0.4 | 1.9 | S | AGC | 1.8 |
| 119 | P | CCT | 0.6 | 1 | 4.7 | 0 | 0.2 | 0.3 | 1.2 | 2.5 | S | TCT | 2.6 |
| 119 | P | CCT | 0.6 | 1 | 4.7 | 0 | 0.2 | 0.3 | 1.2 | 2.5 | S | TCC | 6 |
| 119 | P | CCT | 0.6 | 1 | 4.7 | 0 | 0.2 | 0.3 | 1.2 | 2.5 | S | TCA | 8.1 |
| 119 | P | CCT | 0.6 | 1 | 4.7 | 0 | 0.2 | 0.3 | 1.2 | 2.5 | S | TCG | 44.8 |
| 119 | P | CCT | 0.6 | 1 | 4.7 | 0 | 0.2 | 0.3 | 1.2 | 2.5 | S | AGT | 4.6 |
| 119 | P | CCT | 0.6 | 1 | 4.7 | 0 | 0.2 | 0.3 | 1.2 | 2.5 | S | AGC | 6.6 |
| 119 | T | ACT | 0 | 0.4 | 3.3 | 0 | 0.1 | 0.6 | 2.9 | 0 | S | TCT | 7.4 |
| 119 | T | ACT | 0 | 0.4 | 3.3 | 0 | 0.1 | 0.6 | 2.9 | 0 | S | TCC | 10.8 |
| 119 | T | ACT | 0 | 0.4 | 3.3 | 0 | 0.1 | 0.6 | 2.9 | 0 | S | TCA | 12.9 |
| 119 | T | ACT | 0 | 0.4 | 3.3 | 0 | 0.1 | 0.6 | 2.9 | 0 | S | TCG | 49.6 |
| 119 | T | ACT | 0 | 0.4 | 3.3 | 0 | 0.1 | 0.6 | 2.9 | 0 | S | AGT | 3.6 |
| 119 | T | ACT | 0 | 0.4 | 3.3 | 0 | 0.1 | 0.6 | 2.9 | 0 | S | AGC | 7 |

| | | | | | | | | | | | | | |
|---|---|---|---|---|---|---|---|---|---|---|---|---|---|
| 119 | R | AGG | 0 | 0.2 | 0.2 | 0.1 | | 0.6 | 0.6 | 0 | 1.2 | S | TCT | 56 |
| 119 | R | AGG | 0 | 0.2 | 0.2 | 0.1 | | 0.6 | 0.6 | 0 | 1.2 | S | TCC | 48.5 |
| 119 | R | AGG | 0 | 0.2 | 0.2 | 0.1 | | 0.6 | 0.6 | 0 | 1.2 | S | TCA | 11.7 |
| 119 | R | AGG | 0 | 0.2 | 0.2 | 0.1 | | 0.6 | 0.6 | 0 | 1.2 | S | TCG | 10.3 |
| 119 | R | AGG | 0 | 0.2 | 0.2 | 0.1 | | 0.6 | 0.6 | 0 | 1.2 | S | AGT | 13.5 |
| 119 | R | AGG | 0 | 0.2 | 0.2 | 0.1 | | 0.6 | 0.6 | 0 | 1.2 | S | AGC | 5.8 |
| 119 | S | TCC | 0.7 | 0.3 | 0 | 0 | | 0.3 | 1.2 | 0.4 | 0 | S | TCT | 2.6 |
| 119 | S | TCC | 0.7 | 0.3 | 0 | 0 | | 0.3 | 1.2 | 0.4 | 0 | S | TCC | 0 |
| 119 | S | TCC | 0.7 | 0.3 | 0 | 0 | | 0.3 | 1.2 | 0.4 | 0 | S | TCA | 1.2 |
| 119 | S | TCC | 0.7 | 0.3 | 0 | 0 | | 0.3 | 1.2 | 0.4 | 0 | S | TCG | 3.6 |
| 119 | S | TCC | 0.7 | 0.3 | 0 | 0 | | 0.3 | 1.2 | 0.4 | 0 | S | AGT | 7.4 |
| 119 | S | TCC | 0.7 | 0.3 | 0 | 0 | | 0.3 | 1.2 | 0.4 | 0 | S | AGC | 6.4 |
| 121 | F | TTC | 94.8 | 87.2 | 95.4 | 97.2 | | 95.6 | 72.1 | 94.6 | 93.8 | Y | TAT | 4.9 |
| 121 | F | TTC | 94.8 | 87.2 | 95.4 | 97.2 | | 95.6 | 72.1 | 94.6 | 93.8 | Y | TAC | 3 |
| 121 | F | TTT | 5.2 | 12.8 | 4.5 | 2.8 | | 4.4 | 27.9 | 5.1 | 6.2 | Y | TAT | 3 |
| 121 | F | TTT | 5.2 | 12.8 | 4.5 | 2.8 | | 4.4 | 27.9 | 5.1 | 6.2 | Y | TAC | 6.4 |
| 124 | A | GCT | 65.5 | 18.9 | 60.3 | 83 | | 87.4 | 65 | 23.4 | 61.2 | A | GCT | 0 |
| 124 | A | GCT | 65.5 | 18.9 | 60.3 | 83 | | 87.4 | 65 | 23.4 | 61.2 | A | GCC | 1.6 |
| 124 | A | GCT | 65.5 | 18.9 | 60.3 | 83 | | 87.4 | 65 | 23.4 | 61.2 | A | GCA | 3 |
| 124 | A | GCT | 65.5 | 18.9 | 60.3 | 83 | | 87.4 | 65 | 23.4 | 61.2 | A | GCG | 5.3 |
| 124 | T | ACT | 3.7 | 48.5 | 12.8 | 7.6 | | 3.3 | 10.6 | 23 | 5.6 | A | GCT | 1 |
| 124 | T | ACT | 3.7 | 48.5 | 12.8 | 7.6 | | 3.3 | 10.6 | 23 | 5.6 | A | GCC | 4.4 |
| 124 | T | ACT | 3.7 | 48.5 | 12.8 | 7.6 | | 3.3 | 10.6 | 23 | 5.6 | A | GCA | 6.5 |
| 124 | T | ACT | 3.7 | 48.5 | 12.8 | 7.6 | | 3.3 | 10.6 | 23 | 5.6 | A | GCG | 43.2 |
| 124 | N | AAT | 3.7 | 18.3 | 19 | 5.2 | | 2.2 | 7.6 | 4.1 | 21.7 | A | GCT | 9.2 |
| 124 | N | AAT | 3.7 | 18.3 | 19 | 5.2 | | 2.2 | 7.6 | 4.1 | 21.7 | A | GCC | 11.2 |
| 124 | N | AAT | 3.7 | 18.3 | 19 | 5.2 | | 2.2 | 7.6 | 4.1 | 21.7 | A | GCA | 13.1 |
| 124 | N | AAT | 3.7 | 18.3 | 19 | 5.2 | | 2.2 | 7.6 | 4.1 | 21.7 | A | GCG | 60.4 |
| 124 | T | ACC | 0 | 5.7 | 0.2 | 0.2 | | 0 | 0 | 4.7 | 0 | A | GCT | 2.9 |
| 124 | T | ACC | 0 | 5.7 | 0.2 | 0.2 | | 0 | 0 | 4.7 | 0 | A | GCC | 1 |
| 124 | T | ACC | 0 | 5.7 | 0.2 | 0.2 | | 0 | 0 | 4.7 | 0 | A | GCA | 5 |
| 124 | T | ACC | 0 | 5.7 | 0.2 | 0.2 | | 0 | 0 | 4.7 | 0 | A | GCG | 4.4 |
| 124 | S | AGT | 20 | 1.2 | 3.9 | 0.4 | | 0.8 | 0.9 | 2 | 5.6 | A | GCT | 3.9 |
| 124 | S | AGT | 20 | 1.2 | 3.9 | 0.4 | | 0.8 | 0.9 | 2 | 5.6 | A | GCC | 5.9 |

| | | | | | | | | | | | | | |
|---|---|---|---|---|---|---|---|---|---|---|---|---|---|
| 124 | S | AGT | 20 | 1.2 | 3.9 | 0.4 | 0.8 | 0.9 | 2 | 5.6 | A | GCA | 7.8 |
| 124 | S | AGT | 20 | 1.2 | 3.9 | 0.4 | 0.8 | 0.9 | 2 | 5.6 | A | GCG | 55.1 |
| 124 | A | GCC | 0.7 | 1.3 | 0.3 | 0.4 | 1.1 | 0 | 32.8 | 0.3 | A | GCT | 2.6 |
| 124 | A | GCC | 0.7 | 1.3 | 0.3 | 0.4 | 1.1 | 0 | 32.8 | 0.3 | A | GCC | 0 |
| 124 | A | GCC | 0.7 | 1.3 | 0.3 | 0.4 | 1.1 | 0 | 32.8 | 0.3 | A | GCA | 1.2 |
| 124 | A | GCC | 0.7 | 1.3 | 0.3 | 0.4 | 1.1 | 0 | 32.8 | 0.3 | A | GCG | 3.6 |
| 124 | N | AAC | 0 | 2.4 | 0.6 | 0.1 | 0 | 0.3 | 1.6 | 0.6 | A | GCT | 10.2 |
| 124 | N | AAC | 0 | 2.4 | 0.6 | 0.1 | 0 | 0.3 | 1.6 | 0.6 | A | GCC | 9.2 |
| 124 | N | AAC | 0 | 2.4 | 0.6 | 0.1 | 0 | 0.3 | 1.6 | 0.6 | A | GCA | 10.5 |
| 124 | N | AAC | 0 | 2.4 | 0.6 | 0.1 | 0 | 0.3 | 1.6 | 0.6 | A | GCG | 30.2 |
| 124 | A | GCA | 2.1 | 0.4 | 1 | 0.1 | 0.9 | 2.9 | 2.3 | 1.2 | A | GCT | 7.4 |
| 124 | A | GCA | 2.1 | 0.4 | 1 | 0.1 | 0.9 | 2.9 | 2.3 | 1.2 | A | GCC | 1.8 |
| 124 | A | GCA | 2.1 | 0.4 | 1 | 0.1 | 0.9 | 2.9 | 2.3 | 1.2 | A | GCA | 0 |
| 124 | A | GCA | 2.1 | 0.4 | 1 | 0.1 | 0.9 | 2.9 | 2.3 | 1.2 | A | GCG | 1 |
| 124 | A | GCG | 0.7 | 0.7 | 0.3 | 0.8 | 0.6 | 7.9 | 0 | 0 | A | GCT | 13.5 |
| 124 | A | GCG | 0.7 | 0.7 | 0.3 | 0.8 | 0.6 | 7.9 | 0 | 0 | A | GCC | 5.8 |
| 124 | A | GCG | 0.7 | 0.7 | 0.3 | 0.8 | 0.6 | 7.9 | 0 | 0 | A | GCA | 1 |
| 124 | A | GCG | 0.7 | 0.7 | 0.3 | 0.8 | 0.6 | 7.9 | 0 | 0 | A | GCG | 0 |
| 124 | G | GGT | 2.1 | 0.1 | 0.5 | 0.8 | 2.6 | 3.2 | 3.3 | 2.5 | A | GCT | 5.8 |
| 124 | G | GGT | 2.1 | 0.1 | 0.5 | 0.8 | 2.6 | 3.2 | 3.3 | 2.5 | A | GCC | 9.2 |
| 124 | G | GGT | 2.1 | 0.1 | 0.5 | 0.8 | 2.6 | 3.2 | 3.3 | 2.5 | A | GCA | 11.3 |
| 124 | G | GGT | 2.1 | 0.1 | 0.5 | 0.8 | 2.6 | 3.2 | 3.3 | 2.5 | A | GCG | 48 |
| 124 | T | ACA | 0.2 | 1.1 | 0.2 | 0.1 | 0.2 | 0 | 0 | 0 | A | GCT | 7.4 |
| 124 | T | ACA | 0.2 | 1.1 | 0.2 | 0.1 | 0.2 | 0 | 0 | 0 | A | GCC | 9.2 |
| 124 | T | ACA | 0.2 | 1.1 | 0.2 | 0.1 | 0.2 | 0 | 0 | 0 | A | GCA | 1 |
| 124 | T | ACA | 0.2 | 1.1 | 0.2 | 0.1 | 0.2 | 0 | 0 | 0 | A | GCG | 3.5 |
| 125 | A | GCA | 90.4 | 11.4 | 81.7 | 96.1 | 88 | 41.5 | 20.6 | 73.9 | K | AAA | 5 |
| 125 | A | GCA | 90.4 | 11.4 | 81.7 | 96.1 | 88 | 41.5 | 20.6 | 73.9 | K | AAG | 8.3 |
| 125 | T | ACA | 4.3 | 38.6 | 13.9 | 1.2 | 6.8 | 23.2 | 6 | 18 | K | AAA | 1.2 |
| 125 | T | ACA | 4.3 | 38.6 | 13.9 | 1.2 | 6.8 | 23.2 | 6 | 18 | K | AAG | 3.7 |
| 125 | T | ACG | 0 | 22.3 | 0 | 0.1 | 0 | 0.9 | 8 | 0 | K | AAA | 2.2 |
| 125 | T | ACG | 0 | 22.3 | 0 | 0.1 | 0 | 0.9 | 8 | 0 | K | AAG | 1.2 |
| 125 | A | GCG | 1.1 | 13.6 | 1.1 | 1 | 0.7 | 2.1 | 41.2 | 3.7 | K | AAA | 6.4 |
| 125 | A | GCG | 1.1 | 13.6 | 1.1 | 1 | 0.7 | 2.1 | 41.2 | 3.7 | K | AAG | 5 |

| | | | | | | | | | | | | | |
|---|---|---|---|---|---|---|---|---|---|---|---|---|---|
| 125 | V | GTG | 0.1 | 2.8 | 0.2 | 0 | 0 | 0.6 | 17.7 | 0 | K | AAA | 7.9 |
| 125 | V | GTG | 0.1 | 2.8 | 0.2 | 0 | 0 | 0.6 | 17.7 | 0 | K | AAG | 6.5 |
| 125 | T | ACT | 0 | 3.1 | 0 | 0 | 0.1 | 9.4 | 1.9 | 0.6 | K | AAA | 6.7 |
| 125 | T | ACT | 0 | 3.1 | 0 | 0 | 0.1 | 9.4 | 1.9 | 0.6 | K | AAG | 43.4 |
| 125 | T | ACC | 0 | 2.1 | 0.1 | 0 | 0 | 5 | 0 | 0 | K | AAA | 5.2 |
| 125 | T | ACC | 0 | 2.1 | 0.1 | 0 | 0 | 5 | 0 | 0 | K | AAG | 4.6 |
| 125 | S | TCA | 1.5 | 2 | 0.1 | 0 | 0.7 | 1.2 | 0.4 | 0 | K | AAA | 7 |
| 125 | S | TCA | 1.5 | 2 | 0.1 | 0 | 0.7 | 1.2 | 0.4 | 0 | K | AAG | 10.3 |
| 125 | V | GTA | 1.3 | 0.6 | 1.3 | 0.4 | 1.5 | 1.8 | 1.4 | 2.5 | K | AAA | 6.5 |
| 125 | V | GTA | 1.3 | 0.6 | 1.3 | 0.4 | 1.5 | 1.8 | 1.4 | 2.5 | K | AAG | 9.8 |
| 125 | S | TCG | 0 | 1.3 | 0 | 0 | 0 | 0.6 | 0 | 0 | K | AAA | 8.4 |
| 125 | S | TCG | 0 | 1.3 | 0 | 0 | 0 | 0.6 | 0 | 0 | K | AAG | 7 |
| 125 | A | GCT | 0.2 | 0.5 | 0.5 | 0.5 | 0.8 | 6.2 | 1.2 | 0.6 | K | AAA | 8.9 |
| 125 | A | GCT | 0.2 | 0.5 | 0.5 | 0.5 | 0.8 | 6.2 | 1.2 | 0.6 | K | AAG | 56.2 |
| 125 | A | GCC | 0.7 | 0 | 0.2 | 0.6 | 1.2 | 7.1 | 0 | 0 | K | AAA | 6.3 |
| 125 | A | GCC | 0.7 | 0 | 0.2 | 0.6 | 1.2 | 7.1 | 0 | 0 | K | AAG | 26 |
| 128 | A | GCA | 75.9 | 1.2 | 97.5 | 98.4 | 97.6 | 10.3 | 91.4 | 96.3 | T | ACT | 7.4 |
| 128 | A | GCA | 75.9 | 1.2 | 97.5 | 98.4 | 97.6 | 10.3 | 91.4 | 96.3 | T | ACC | 9.2 |
| 128 | A | GCA | 75.9 | 1.2 | 97.5 | 98.4 | 97.6 | 10.3 | 91.4 | 96.3 | T | ACA | 1 |
| 128 | A | GCA | 75.9 | 1.2 | 97.5 | 98.4 | 97.6 | 10.3 | 91.4 | 96.3 | T | ACG | 3.5 |
| 128 | A | GCC | 1 | 79.5 | 0.8 | 0.4 | 0.7 | 74.7 | 7.4 | 1.2 | T | ACT | 2.9 |
| 128 | A | GCC | 1 | 79.5 | 0.8 | 0.4 | 0.7 | 74.7 | 7.4 | 1.2 | T | ACC | 1 |
| 128 | A | GCC | 1 | 79.5 | 0.8 | 0.4 | 0.7 | 74.7 | 7.4 | 1.2 | T | ACA | 5 |
| 128 | A | GCC | 1 | 79.5 | 0.8 | 0.4 | 0.7 | 74.7 | 7.4 | 1.2 | T | ACG | 4.4 |
| 128 | A | GCT | 22.4 | 16.5 | 0.2 | 0.4 | 0.8 | 13.2 | 0.4 | 2.2 | T | ACT | 1 |
| 128 | A | GCT | 22.4 | 16.5 | 0.2 | 0.4 | 0.8 | 13.2 | 0.4 | 2.2 | T | ACC | 4.4 |
| 128 | A | GCT | 22.4 | 16.5 | 0.2 | 0.4 | 0.8 | 13.2 | 0.4 | 2.2 | T | ACA | 6.5 |
| 128 | A | GCT | 22.4 | 16.5 | 0.2 | 0.4 | 0.8 | 13.2 | 0.4 | 2.2 | T | ACG | 43.2 |
| 128 | A | GCG | 0.5 | 2.4 | 0.7 | 0.6 | 0.8 | 1.8 | 0.4 | 0.3 | T | ACT | 21.2 |
| 128 | A | GCG | 0.5 | 2.4 | 0.7 | 0.6 | 0.8 | 1.8 | 0.4 | 0.3 | T | ACC | 3.9 |
| 128 | A | GCG | 0.5 | 2.4 | 0.7 | 0.6 | 0.8 | 1.8 | 0.4 | 0.3 | T | ACA | 2 |
| 128 | A | GCG | 0.5 | 2.4 | 0.7 | 0.6 | 0.8 | 1.8 | 0.4 | 0.3 | T | ACG | 1 |
| 138 | E | GAA | 97.6 | 97.3 | 97 | 79.7 | 97.5 | 97.1 | 92.6 | 98.1 | A | GCT | 8.2 |
| 138 | E | GAA | 97.6 | 97.3 | 97 | 79.7 | 97.5 | 97.1 | 92.6 | 98.1 | A | GCC | 10 |

| | | | | | | | | | | | | |
|---|---|---|---|---|---|---|---|---|---|---|---|---|
| 138 | E | GAA | 97.6 | 97.3 | 97 | 79.7 | 97.5 | 97.1 | 92.6 | 98.1 | A | GCA | 1.8 |
| 138 | E | GAA | 97.6 | 97.3 | 97 | 79.7 | 97.5 | 97.1 | 92.6 | 98.1 | A | GCG | 4.3 |
| 138 | E | GAG | 2 | 1.8 | 1.6 | 19.8 | 2.3 | 1.8 | 6.2 | 1.9 | A | GCT | 22 |
| 138 | E | GAG | 2 | 1.8 | 1.6 | 19.8 | 2.3 | 1.8 | 6.2 | 1.9 | A | GCC | 4.7 |
| 138 | E | GAG | 2 | 1.8 | 1.6 | 19.8 | 2.3 | 1.8 | 6.2 | 1.9 | A | GCA | 2.8 |
| 138 | E | GAG | 2 | 1.8 | 1.6 | 19.8 | 2.3 | 1.8 | 6.2 | 1.9 | A | GCG | 1.8 |
| 138 | D | GAC | 0.4 | 0.5 | 1 | 0.4 | 0 | 0.9 | 1.2 | 0 | A | GCT | 3.7 |
| 138 | D | GAC | 0.4 | 0.5 | 1 | 0.4 | 0 | 0.9 | 1.2 | 0 | A | GCC | 1.8 |
| 138 | D | GAC | 0.4 | 0.5 | 1 | 0.4 | 0 | 0.9 | 1.2 | 0 | A | GCA | 5.8 |
| 138 | D | GAC | 0.4 | 0.5 | 1 | 0.4 | 0 | 0.9 | 1.2 | 0 | A | GCG | 5.2 |
| 138 | E | GAA | 97.6 | 97.3 | 97 | 79.7 | 97.5 | 97.1 | 92.6 | 98.1 | T | ACT | 17.5 |
| 138 | E | GAA | 97.6 | 97.3 | 97 | 79.7 | 97.5 | 97.1 | 92.6 | 98.1 | T | ACC | 14.9 |
| 138 | E | GAA | 97.6 | 97.3 | 97 | 79.7 | 97.5 | 97.1 | 92.6 | 98.1 | T | ACA | 9.2 |
| 138 | E | GAA | 97.6 | 97.3 | 97 | 79.7 | 97.5 | 97.1 | 92.6 | 98.1 | T | ACG | 12.5 |
| 138 | E | GAG | 2 | 1.8 | 1.6 | 19.8 | 2.3 | 1.8 | 6.2 | 1.9 | T | ACT | 54.9 |
| 138 | E | GAG | 2 | 1.8 | 1.6 | 19.8 | 2.3 | 1.8 | 6.2 | 1.9 | T | ACC | 47.4 |
| 138 | E | GAG | 2 | 1.8 | 1.6 | 19.8 | 2.3 | 1.8 | 6.2 | 1.9 | T | ACA | 10.6 |
| 138 | E | GAG | 2 | 1.8 | 1.6 | 19.8 | 2.3 | 1.8 | 6.2 | 1.9 | T | ACG | 9.2 |
| 138 | D | GAC | 0.4 | 0.5 | 1 | 0.4 | 0 | 0.9 | 1.2 | 0 | T | ACT | 10.2 |
| 138 | D | GAC | 0.4 | 0.5 | 1 | 0.4 | 0 | 0.9 | 1.2 | 0 | T | ACC | 9.2 |
| 138 | D | GAC | 0.4 | 0.5 | 1 | 0.4 | 0 | 0.9 | 1.2 | 0 | T | ACA | 10.5 |
| 138 | D | GAC | 0.4 | 0.5 | 1 | 0.4 | 0 | 0.9 | 1.2 | 0 | T | ACG | 30.2 |
| 138 | E | GAA | 97.6 | 97.3 | 97 | 79.7 | 97.5 | 97.1 | 92.6 | 98.1 | K | AAA | 1 |
| 138 | E | GAA | 97.6 | 97.3 | 97 | 79.7 | 97.5 | 97.1 | 92.6 | 98.1 | K | AAG | 3.5 |
| 138 | E | GAG | 2 | 1.8 | 1.6 | 19.8 | 2.3 | 1.8 | 6.2 | 1.9 | K | AAA | 2 |
| 138 | E | GAG | 2 | 1.8 | 1.6 | 19.8 | 2.3 | 1.8 | 6.2 | 1.9 | K | AAG | 1 |
| 138 | D | GAC | 0.4 | 0.5 | 1 | 0.4 | 0 | 0.9 | 1.2 | 0 | K | AAA | 5 |
| 138 | D | GAC | 0.4 | 0.5 | 1 | 0.4 | 0 | 0.9 | 1.2 | 0 | K | AAG | 4.4 |
| 140 | G | GGC | 0.2 | 83.6 | 0.2 | 0.2 | 0 | 15.9 | 5.8 | 1.2 | A | GCT | 7.7 |
| 140 | G | GGC | 0.2 | 83.6 | 0.2 | 0.2 | 0 | 15.9 | 5.8 | 1.2 | A | GCC | 5.8 |
| 140 | G | GGC | 0.2 | 83.6 | 0.2 | 0.2 | 0 | 15.9 | 5.8 | 1.2 | A | GCA | 9.8 |
| 140 | G | GGC | 0.2 | 83.6 | 0.2 | 0.2 | 0 | 15.9 | 5.8 | 1.2 | A | GCG | 9.2 |
| 140 | G | GGA | 56.6 | 3.2 | 80.3 | 6.3 | 97.6 | 72.6 | 83.9 | 95.3 | A | GCT | 12.2 |
| 140 | G | GGA | 56.6 | 3.2 | 80.3 | 6.3 | 97.6 | 72.6 | 83.9 | 95.3 | A | GCC | 14 |

| | | | | | | | | | | | | | |
|---|---|---|---|---|---|---|---|---|---|---|---|---|---|
| 140 | G | GGA | 56.6 | 3.2 | 80.3 | 6.3 | 97.6 | 72.6 | 83.9 | 95.3 | A | GCA | 5.8 |
| 140 | G | GGA | 56.6 | 3.2 | 80.3 | 6.3 | 97.6 | 72.6 | 83.9 | 95.3 | A | GCG | 8.3 |
| 140 | G | GGG | 42.4 | 0.9 | 19.4 | 93.4 | 2.1 | 10.3 | 7.2 | 3.4 | A | GCT | 26 |
| 140 | G | GGG | 42.4 | 0.9 | 19.4 | 93.4 | 2.1 | 10.3 | 7.2 | 3.4 | A | GCC | 8.7 |
| 140 | G | GGG | 42.4 | 0.9 | 19.4 | 93.4 | 2.1 | 10.3 | 7.2 | 3.4 | A | GCA | 6.8 |
| 140 | G | GGG | 42.4 | 0.9 | 19.4 | 93.4 | 2.1 | 10.3 | 7.2 | 3.4 | A | GCG | 5.8 |
| 140 | G | GGT | 0.7 | 12.2 | 0 | 0.1 | 0.2 | 1.2 | 2.5 | 0 | A | GCT | 5.8 |
| 140 | G | GGT | 0.7 | 12.2 | 0 | 0.1 | 0.2 | 1.2 | 2.5 | 0 | A | GCC | 9.2 |
| 140 | G | GGT | 0.7 | 12.2 | 0 | 0.1 | 0.2 | 1.2 | 2.5 | 0 | A | GCA | 11.3 |
| 140 | G | GGT | 0.7 | 12.2 | 0 | 0.1 | 0.2 | 1.2 | 2.5 | 0 | A | GCG | 48 |
| 140 | G | GGC | 0.2 | 83.6 | 0.2 | 0.2 | 0 | 15.9 | 5.8 | 1.2 | C | TGT | 15.4 |
| 140 | G | GGC | 0.2 | 83.6 | 0.2 | 0.2 | 0 | 15.9 | 5.8 | 1.2 | C | TGC | 13.5 |
| 140 | G | GGA | 56.6 | 3.2 | 80.3 | 6.3 | 97.6 | 72.6 | 83.9 | 95.3 | C | TGT | 19.9 |
| 140 | G | GGA | 56.6 | 3.2 | 80.3 | 6.3 | 97.6 | 72.6 | 83.9 | 95.3 | C | TGC | 21.7 |
| 140 | G | GGG | 42.4 | 0.9 | 19.4 | 93.4 | 2.1 | 10.3 | 7.2 | 3.4 | C | TGT | 33.7 |
| 140 | G | GGG | 42.4 | 0.9 | 19.4 | 93.4 | 2.1 | 10.3 | 7.2 | 3.4 | C | TGC | 16.4 |
| 140 | G | GGT | 0.7 | 12.2 | 0 | 0.1 | 0.2 | 1.2 | 2.5 | 0 | C | TGT | 13.5 |
| 140 | G | GGT | 0.7 | 12.2 | 0 | 0.1 | 0.2 | 1.2 | 2.5 | 0 | C | TGC | 16.9 |
| 140 | G | GGC | 0.2 | 83.6 | 0.2 | 0.2 | 0 | 15.9 | 5.8 | 1.2 | S | TCT | 17.4 |
| 140 | G | GGC | 0.2 | 83.6 | 0.2 | 0.2 | 0 | 15.9 | 5.8 | 1.2 | S | TCC | 16.4 |
| 140 | G | GGC | 0.2 | 83.6 | 0.2 | 0.2 | 0 | 15.9 | 5.8 | 1.2 | S | TCA | 17.7 |
| 140 | G | GGC | 0.2 | 83.6 | 0.2 | 0.2 | 0 | 15.9 | 5.8 | 1.2 | S | TCG | 37.4 |
| 140 | G | GGC | 0.2 | 83.6 | 0.2 | 0.2 | 0 | 15.9 | 5.8 | 1.2 | S | AGT | 2.9 |
| 140 | G | GGC | 0.2 | 83.6 | 0.2 | 0.2 | 0 | 15.9 | 5.8 | 1.2 | S | AGC | 1 |
| 140 | G | GGA | 56.6 | 3.2 | 80.3 | 6.3 | 97.6 | 72.6 | 83.9 | 95.3 | S | TCT | 24.7 |
| 140 | G | GGA | 56.6 | 3.2 | 80.3 | 6.3 | 97.6 | 72.6 | 83.9 | 95.3 | S | TCC | 22.1 |
| 140 | G | GGA | 56.6 | 3.2 | 80.3 | 6.3 | 97.6 | 72.6 | 83.9 | 95.3 | S | TCA | 16.4 |
| 140 | G | GGA | 56.6 | 3.2 | 80.3 | 6.3 | 97.6 | 72.6 | 83.9 | 95.3 | S | TCG | 19.7 |
| 140 | G | GGA | 56.6 | 3.2 | 80.3 | 6.3 | 97.6 | 72.6 | 83.9 | 95.3 | S | AGT | 7.4 |
| 140 | G | GGA | 56.6 | 3.2 | 80.3 | 6.3 | 97.6 | 72.6 | 83.9 | 95.3 | S | AGC | 9.2 |
| 140 | G | GGG | 42.4 | 0.9 | 19.4 | 93.4 | 2.1 | 10.3 | 7.2 | 3.4 | S | TCT | 62.1 |
| 140 | G | GGG | 42.4 | 0.9 | 19.4 | 93.4 | 2.1 | 10.3 | 7.2 | 3.4 | S | TCC | 54.6 |
| 140 | G | GGG | 42.4 | 0.9 | 19.4 | 93.4 | 2.1 | 10.3 | 7.2 | 3.4 | S | TCA | 17.8 |
| 140 | G | GGG | 42.4 | 0.9 | 19.4 | 93.4 | 2.1 | 10.3 | 7.2 | 3.4 | S | TCG | 16.4 |

| | | | | | | | | | | | | |
|---|---|---|---|---|---|---|---|---|---|---|---|---|
| 140 | G | GGG | 42.4 | 0.9 | 19.4 | 93.4 | 2.1 | 10.3 | 7.2 | 3.4 | S | AGT | 21.2 |
| 140 | G | GGG | 42.4 | 0.9 | 19.4 | 93.4 | 2.1 | 10.3 | 7.2 | 3.4 | S | AGC | 3.9 |
| 140 | G | GGT | 0.7 | 12.2 | 0 | 0.1 | 0.2 | 1.2 | 2.5 | 0 | S | TCT | 16.4 |
| 140 | G | GGT | 0.7 | 12.2 | 0 | 0.1 | 0.2 | 1.2 | 2.5 | 0 | S | TCC | 18.4 |
| 140 | G | GGT | 0.7 | 12.2 | 0 | 0.1 | 0.2 | 1.2 | 2.5 | 0 | S | TCA | 20.3 |
| 140 | G | GGT | 0.7 | 12.2 | 0 | 0.1 | 0.2 | 1.2 | 2.5 | 0 | S | TCG | 67.6 |
| 140 | G | GGT | 0.7 | 12.2 | 0 | 0.1 | 0.2 | 1.2 | 2.5 | 0 | S | AGT | 1 |
| 140 | G | GGT | 0.7 | 12.2 | 0 | 0.1 | 0.2 | 1.2 | 2.5 | 0 | S | AGC | 4.4 |
| 142 | P | CCC | 89.6 | 96.7 | 99.5 | 97 | 98.7 | 95.9 | 99.6 | 98.8 | T | ACT | 3.1 |
| 142 | P | CCC | 89.6 | 96.7 | 99.5 | 97 | 98.7 | 95.9 | 99.6 | 98.8 | T | ACC | 1.2 |
| 142 | P | CCC | 89.6 | 96.7 | 99.5 | 97 | 98.7 | 95.9 | 99.6 | 98.8 | T | ACA | 5.2 |
| 142 | P | CCC | 89.6 | 96.7 | 99.5 | 97 | 98.7 | 95.9 | 99.6 | 98.8 | T | ACG | 4.6 |
| 142 | P | CCG | 0.2 | 2.4 | 0 | 0 | 0 | 2.1 | 0 | 0 | T | ACT | 21.4 |
| 142 | P | CCG | 0.2 | 2.4 | 0 | 0 | 0 | 2.1 | 0 | 0 | T | ACC | 4.1 |
| 142 | P | CCG | 0.2 | 2.4 | 0 | 0 | 0 | 2.1 | 0 | 0 | T | ACA | 2.2 |
| 142 | P | CCG | 0.2 | 2.4 | 0 | 0 | 0 | 2.1 | 0 | 0 | T | ACG | 1.2 |
| 142 | P | CCT | 9.9 | 0.5 | 0.4 | 2.8 | 1.1 | 1.5 | 0.4 | 0.6 | T | ACT | 1.2 |
| 142 | P | CCT | 9.9 | 0.5 | 0.4 | 2.8 | 1.1 | 1.5 | 0.4 | 0.6 | T | ACC | 4.6 |
| 142 | P | CCT | 9.9 | 0.5 | 0.4 | 2.8 | 1.1 | 1.5 | 0.4 | 0.6 | T | ACA | 6.7 |
| 142 | P | CCT | 9.9 | 0.5 | 0.4 | 2.8 | 1.1 | 1.5 | 0.4 | 0.6 | T | ACG | 43.4 |
| 143 | Y | TAC | 98.3 | 96.9 | 98.8 | 93.1 | 98.6 | 97.6 | 97.3 | 98.8 | A | GCT | 14.5 |
| 143 | Y | TAC | 98.3 | 96.9 | 98.8 | 93.1 | 98.6 | 97.6 | 97.3 | 98.8 | A | GCC | 13.5 |
| 143 | Y | TAC | 98.3 | 96.9 | 98.8 | 93.1 | 98.6 | 97.6 | 97.3 | 98.8 | A | GCA | 14.8 |
| 143 | Y | TAC | 98.3 | 96.9 | 98.8 | 93.1 | 98.6 | 97.6 | 97.3 | 98.8 | A | GCG | 34.5 |
| 143 | Y | TAT | 1.7 | 3.1 | 0.9 | 6.9 | 1.4 | 2.4 | 2.7 | 1.2 | A | GCT | 13.5 |
| 143 | Y | TAT | 1.7 | 3.1 | 0.9 | 6.9 | 1.4 | 2.4 | 2.7 | 1.2 | A | GCC | 15.5 |
| 143 | Y | TAT | 1.7 | 3.1 | 0.9 | 6.9 | 1.4 | 2.4 | 2.7 | 1.2 | A | GCA | 17.4 |
| 143 | Y | TAT | 1.7 | 3.1 | 0.9 | 6.9 | 1.4 | 2.4 | 2.7 | 1.2 | A | GCG | 64.7 |
| 143 | Y | TAC | 98.3 | 96.9 | 98.8 | 93.1 | 98.6 | 97.6 | 97.3 | 98.8 | C | TGT | 2.9 |
| 143 | Y | TAC | 98.3 | 96.9 | 98.8 | 93.1 | 98.6 | 97.6 | 97.3 | 98.8 | C | TGC | 1 |
| 143 | Y | TAT | 1.7 | 3.1 | 0.9 | 6.9 | 1.4 | 2.4 | 2.7 | 1.2 | C | TGT | 1 |
| 143 | Y | TAT | 1.7 | 3.1 | 0.9 | 6.9 | 1.4 | 2.4 | 2.7 | 1.2 | C | TGC | 4.4 |
| 143 | Y | TAC | 98.3 | 96.9 | 98.8 | 93.1 | 98.6 | 97.6 | 97.3 | 98.8 | G | GGT | 8.8 |
| 143 | Y | TAC | 98.3 | 96.9 | 98.8 | 93.1 | 98.6 | 97.6 | 97.3 | 98.8 | G | GGC | 7.8 |

| | | | | | | | | | | | | | |
|---|---|---|---|---|---|---|---|---|---|---|---|---|---|
| 143 | Y | TAC | 98.3 | 96.9 | 98.8 | 93.1 | 98.6 | 97.6 | 97.3 | 98.8 | G | GGA | 9.1 |
| 143 | Y | TAC | 98.3 | 96.9 | 98.8 | 93.1 | 98.6 | 97.6 | 97.3 | 98.8 | G | GGG | 28.8 |
| 143 | Y | TAT | 1.7 | 3.1 | 0.9 | 6.9 | 1.4 | 2.4 | 2.7 | 1.2 | G | GGT | 7.8 |
| 143 | Y | TAT | 1.7 | 3.1 | 0.9 | 6.9 | 1.4 | 2.4 | 2.7 | 1.2 | G | GGC | 9.8 |
| 143 | Y | TAT | 1.7 | 3.1 | 0.9 | 6.9 | 1.4 | 2.4 | 2.7 | 1.2 | G | GGA | 11.7 |
| 143 | Y | TAT | 1.7 | 3.1 | 0.9 | 6.9 | 1.4 | 2.4 | 2.7 | 1.2 | G | GGG | 59 |
| 143 | Y | TAC | 98.3 | 96.9 | 98.8 | 93.1 | 98.6 | 97.6 | 97.3 | 98.8 | H | CAT | 3.5 |
| 143 | Y | TAC | 98.3 | 96.9 | 98.8 | 93.1 | 98.6 | 97.6 | 97.3 | 98.8 | H | CAC | 1.6 |
| 143 | Y | TAT | 1.7 | 3.1 | 0.9 | 6.9 | 1.4 | 2.4 | 2.7 | 1.2 | H | CAT | 1.6 |
| 143 | Y | TAT | 1.7 | 3.1 | 0.9 | 6.9 | 1.4 | 2.4 | 2.7 | 1.2 | H | CAC | 5 |
| 143 | Y | TAC | 98.3 | 96.9 | 98.8 | 93.1 | 98.6 | 97.6 | 97.3 | 98.8 | K | AAA | 7 |
| 143 | Y | TAC | 98.3 | 96.9 | 98.8 | 93.1 | 98.6 | 97.6 | 97.3 | 98.8 | K | AAG | 6.4 |
| 143 | Y | TAT | 1.7 | 3.1 | 0.9 | 6.9 | 1.4 | 2.4 | 2.7 | 1.2 | K | AAA | 8.5 |
| 143 | Y | TAT | 1.7 | 3.1 | 0.9 | 6.9 | 1.4 | 2.4 | 2.7 | 1.2 | K | AAG | 45.2 |
| 143 | Y | TAC | 98.3 | 96.9 | 98.8 | 93.1 | 98.6 | 97.6 | 97.3 | 98.8 | S | TCT | 3.7 |
| 143 | Y | TAC | 98.3 | 96.9 | 98.8 | 93.1 | 98.6 | 97.6 | 97.3 | 98.8 | S | TCC | 1.8 |
| 143 | Y | TAC | 98.3 | 96.9 | 98.8 | 93.1 | 98.6 | 97.6 | 97.3 | 98.8 | S | TCA | 5.8 |
| 143 | Y | TAC | 98.3 | 96.9 | 98.8 | 93.1 | 98.6 | 97.6 | 97.3 | 98.8 | S | TCG | 5.2 |
| 143 | Y | TAC | 98.3 | 96.9 | 98.8 | 93.1 | 98.6 | 97.6 | 97.3 | 98.8 | S | AGT | 6.5 |
| 143 | Y | TAC | 98.3 | 96.9 | 98.8 | 93.1 | 98.6 | 97.6 | 97.3 | 98.8 | S | AGC | 5.5 |
| 143 | Y | TAT | 1.7 | 3.1 | 0.9 | 6.9 | 1.4 | 2.4 | 2.7 | 1.2 | S | TCT | 1.8 |
| 143 | Y | TAT | 1.7 | 3.1 | 0.9 | 6.9 | 1.4 | 2.4 | 2.7 | 1.2 | S | TCC | 5.2 |
| 143 | Y | TAT | 1.7 | 3.1 | 0.9 | 6.9 | 1.4 | 2.4 | 2.7 | 1.2 | S | TCA | 7.3 |
| 143 | Y | TAT | 1.7 | 3.1 | 0.9 | 6.9 | 1.4 | 2.4 | 2.7 | 1.2 | S | TCG | 44 |
| 143 | Y | TAT | 1.7 | 3.1 | 0.9 | 6.9 | 1.4 | 2.4 | 2.7 | 1.2 | S | AGT | 5.5 |
| 143 | Y | TAT | 1.7 | 3.1 | 0.9 | 6.9 | 1.4 | 2.4 | 2.7 | 1.2 | S | AGC | 7.5 |
| 143 | Y | TAC | 98.3 | 96.9 | 98.8 | 93.1 | 98.6 | 97.6 | 97.3 | 98.8 | R | CGT | 5.1 |
| 143 | Y | TAC | 98.3 | 96.9 | 98.8 | 93.1 | 98.6 | 97.6 | 97.3 | 98.8 | R | CGC | 4.1 |
| 143 | Y | TAC | 98.3 | 96.9 | 98.8 | 93.1 | 98.6 | 97.6 | 97.3 | 98.8 | R | CGA | 5.4 |
| 143 | Y | TAC | 98.3 | 96.9 | 98.8 | 93.1 | 98.6 | 97.6 | 97.3 | 98.8 | R | CGG | 25.1 |
| 143 | Y | TAC | 98.3 | 96.9 | 98.8 | 93.1 | 98.6 | 97.6 | 97.3 | 98.8 | R | AGA | 6.8 |
| 143 | Y | TAC | 98.3 | 96.9 | 98.8 | 93.1 | 98.6 | 97.6 | 97.3 | 98.8 | R | AGG | 26.5 |
| 143 | Y | TAT | 1.7 | 3.1 | 0.9 | 6.9 | 1.4 | 2.4 | 2.7 | 1.2 | R | CGT | 4.1 |
| 143 | Y | TAT | 1.7 | 3.1 | 0.9 | 6.9 | 1.4 | 2.4 | 2.7 | 1.2 | R | CGC | 6.1 |

| | | | | | | | | | | | | |
|---|---|---|---|---|---|---|---|---|---|---|---|---|
| 143 | Y | TAT | 1.7 | 3.1 | 0.9 | 6.9 | 1.4 | 2.4 | 2.7 | 1.2 | R | CGA | 8 |
| 143 | Y | TAT | 1.7 | 3.1 | 0.9 | 6.9 | 1.4 | 2.4 | 2.7 | 1.2 | R | CGG | 55.3 |
| 143 | Y | TAT | 1.7 | 3.1 | 0.9 | 6.9 | 1.4 | 2.4 | 2.7 | 1.2 | R | AGA | 9.4 |
| 143 | Y | TAT | 1.7 | 3.1 | 0.9 | 6.9 | 1.4 | 2.4 | 2.7 | 1.2 | R | AGG | 56.7 |
| 145 | P | CCC | 98.3 | 91.4 | 97.3 | 92.5 | 99.3 | 91.5 | 98.4 | 100 | S | TCT | 4.5 |
| 145 | P | CCC | 98.3 | 91.4 | 97.3 | 92.5 | 99.3 | 91.5 | 98.4 | 100 | S | TCC | 2.6 |
| 145 | P | CCC | 98.3 | 91.4 | 97.3 | 92.5 | 99.3 | 91.5 | 98.4 | 100 | S | TCA | 6.6 |
| 145 | P | CCC | 98.3 | 91.4 | 97.3 | 92.5 | 99.3 | 91.5 | 98.4 | 100 | S | TCG | 6 |
| 145 | P | CCC | 98.3 | 91.4 | 97.3 | 92.5 | 99.3 | 91.5 | 98.4 | 100 | S | AGT | 5.6 |
| 145 | P | CCC | 98.3 | 91.4 | 97.3 | 92.5 | 99.3 | 91.5 | 98.4 | 100 | S | AGC | 4.6 |
| 145 | P | CCT | 1.2 | 5.8 | 2.2 | 6.7 | 0.5 | 6.5 | 1.2 | 0 | S | TCT | 2.6 |
| 145 | P | CCT | 1.2 | 5.8 | 2.2 | 6.7 | 0.5 | 6.5 | 1.2 | 0 | S | TCC | 6 |
| 145 | P | CCT | 1.2 | 5.8 | 2.2 | 6.7 | 0.5 | 6.5 | 1.2 | 0 | S | TCA | 8.1 |
| 145 | P | CCT | 1.2 | 5.8 | 2.2 | 6.7 | 0.5 | 6.5 | 1.2 | 0 | S | TCG | 44.8 |
| 145 | P | CCT | 1.2 | 5.8 | 2.2 | 6.7 | 0.5 | 6.5 | 1.2 | 0 | S | AGT | 4.6 |
| 145 | P | CCT | 1.2 | 5.8 | 2.2 | 6.7 | 0.5 | 6.5 | 1.2 | 0 | S | AGC | 6.6 |
| 145 | P | CCG | 0.2 | 2.4 | 0 | 0 | 0 | 1.8 | 0 | 0 | S | TCT | 22.8 |
| 145 | P | CCG | 0.2 | 2.4 | 0 | 0 | 0 | 1.8 | 0 | 0 | S | TCC | 5.5 |
| 145 | P | CCG | 0.2 | 2.4 | 0 | 0 | 0 | 1.8 | 0 | 0 | S | TCA | 3.6 |
| 145 | P | CCG | 0.2 | 2.4 | 0 | 0 | 0 | 1.8 | 0 | 0 | S | TCG | 2.6 |
| 145 | P | CCG | 0.2 | 2.4 | 0 | 0 | 0 | 1.8 | 0 | 0 | S | AGT | 50.3 |
| 145 | P | CCG | 0.2 | 2.4 | 0 | 0 | 0 | 1.8 | 0 | 0 | S | AGC | 42.8 |
| 146 | Q | CAA | 98.8 | 96.1 | 98.6 | 97.9 | 93.8 | 97.6 | 97.1 | 95 | I | ATT | 15.9 |
| 146 | Q | CAA | 98.8 | 96.1 | 98.6 | 97.9 | 93.8 | 97.6 | 97.1 | 95 | I | ATC | 13.3 |
| 146 | Q | CAA | 98.8 | 96.1 | 98.6 | 97.9 | 93.8 | 97.6 | 97.1 | 95 | I | ATA | 7.6 |
| 146 | Q | CAG | 1.2 | 3.8 | 1.4 | 2.1 | 6.2 | 2.4 | 2.9 | 4.7 | I | ATT | 53.3 |
| 146 | Q | CAG | 1.2 | 3.8 | 1.4 | 2.1 | 6.2 | 2.4 | 2.9 | 4.7 | I | ATC | 45.8 |
| 146 | Q | CAG | 1.2 | 3.8 | 1.4 | 2.1 | 6.2 | 2.4 | 2.9 | 4.7 | I | ATA | 9 |
| 146 | Q | CAA | 98.8 | 96.1 | 98.6 | 97.9 | 93.8 | 97.6 | 97.1 | 95 | K | AAA | 1.2 |
| 146 | Q | CAA | 98.8 | 96.1 | 98.6 | 97.9 | 93.8 | 97.6 | 97.1 | 95 | K | AAG | 3.7 |
| 146 | Q | CAG | 1.2 | 3.8 | 1.4 | 2.1 | 6.2 | 2.4 | 2.9 | 4.7 | K | AAA | 2.2 |
| 146 | Q | CAG | 1.2 | 3.8 | 1.4 | 2.1 | 6.2 | 2.4 | 2.9 | 4.7 | K | AAG | 1.2 |
| 146 | Q | CAA | 98.8 | 96.1 | 98.6 | 97.9 | 93.8 | 97.6 | 97.1 | 95 | L | TTA | 9 |
| 146 | Q | CAA | 98.8 | 96.1 | 98.6 | 97.9 | 93.8 | 97.6 | 97.1 | 95 | L | TTG | 12.3 |

| | | | | | | | | | | | | |
|---|---|---|---|---|---|---|---|---|---|---|---|---|
| 146 | Q | CAA | 98.8 | 96.1 | 98.6 | 97.9 | 93.8 | 97.6 | 97.1 | 95 | L | CTT | 13.8 |
| 146 | Q | CAA | 98.8 | 96.1 | 98.6 | 97.9 | 93.8 | 97.6 | 97.1 | 95 | L | CTC | 15.6 |
| 146 | Q | CAA | 98.8 | 96.1 | 98.6 | 97.9 | 93.8 | 97.6 | 97.1 | 95 | L | CTA | 7.4 |
| 146 | Q | CAA | 98.8 | 96.1 | 98.6 | 97.9 | 93.8 | 97.6 | 97.1 | 95 | L | CTG | 9.9 |
| 146 | Q | CAG | 1.2 | 3.8 | 1.4 | 2.1 | 6.2 | 2.4 | 2.9 | 4.7 | L | TTA | 10.4 |
| 146 | Q | CAG | 1.2 | 3.8 | 1.4 | 2.1 | 6.2 | 2.4 | 2.9 | 4.7 | L | TTG | 9 |
| 146 | Q | CAG | 1.2 | 3.8 | 1.4 | 2.1 | 6.2 | 2.4 | 2.9 | 4.7 | L | CTT | 27.6 |
| 146 | Q | CAG | 1.2 | 3.8 | 1.4 | 2.1 | 6.2 | 2.4 | 2.9 | 4.7 | L | CTC | 10.3 |
| 146 | Q | CAG | 1.2 | 3.8 | 1.4 | 2.1 | 6.2 | 2.4 | 2.9 | 4.7 | L | CTA | 8.4 |
| 146 | Q | CAG | 1.2 | 3.8 | 1.4 | 2.1 | 6.2 | 2.4 | 2.9 | 4.7 | L | CTG | 7.4 |
| 146 | Q | CAA | 98.8 | 96.1 | 98.6 | 97.9 | 93.8 | 97.6 | 97.1 | 95 | P | CCT | 8.2 |
| 146 | Q | CAA | 98.8 | 96.1 | 98.6 | 97.9 | 93.8 | 97.6 | 97.1 | 95 | P | CCC | 10 |
| 146 | Q | CAA | 98.8 | 96.1 | 98.6 | 97.9 | 93.8 | 97.6 | 97.1 | 95 | P | CCA | 1.8 |
| 146 | Q | CAA | 98.8 | 96.1 | 98.6 | 97.9 | 93.8 | 97.6 | 97.1 | 95 | P | CCG | 4.3 |
| 146 | Q | CAG | 1.2 | 3.8 | 1.4 | 2.1 | 6.2 | 2.4 | 2.9 | 4.7 | P | CCT | 22 |
| 146 | Q | CAG | 1.2 | 3.8 | 1.4 | 2.1 | 6.2 | 2.4 | 2.9 | 4.7 | P | CCC | 4.7 |
| 146 | Q | CAG | 1.2 | 3.8 | 1.4 | 2.1 | 6.2 | 2.4 | 2.9 | 4.7 | P | CCA | 2.8 |
| 146 | Q | CAG | 1.2 | 3.8 | 1.4 | 2.1 | 6.2 | 2.4 | 2.9 | 4.7 | P | CCG | 1.8 |
| 146 | Q | CAA | 98.8 | 96.1 | 98.6 | 97.9 | 93.8 | 97.6 | 97.1 | 95 | R | CGT | 7.4 |
| 146 | Q | CAA | 98.8 | 96.1 | 98.6 | 97.9 | 93.8 | 97.6 | 97.1 | 95 | R | CGC | 9.2 |
| 146 | Q | CAA | 98.8 | 96.1 | 98.6 | 97.9 | 93.8 | 97.6 | 97.1 | 95 | R | CGA | 1 |
| 146 | Q | CAA | 98.8 | 96.1 | 98.6 | 97.9 | 93.8 | 97.6 | 97.1 | 95 | R | CGG | 3.5 |
| 146 | Q | CAA | 98.8 | 96.1 | 98.6 | 97.9 | 93.8 | 97.6 | 97.1 | 95 | R | AGA | 3.7 |
| 146 | Q | CAA | 98.8 | 96.1 | 98.6 | 97.9 | 93.8 | 97.6 | 97.1 | 95 | R | AGG | 7 |
| 146 | Q | CAG | 1.2 | 3.8 | 1.4 | 2.1 | 6.2 | 2.4 | 2.9 | 4.7 | R | CGT | 21.2 |
| 146 | Q | CAG | 1.2 | 3.8 | 1.4 | 2.1 | 6.2 | 2.4 | 2.9 | 4.7 | R | CGC | 3.9 |
| 146 | Q | CAG | 1.2 | 3.8 | 1.4 | 2.1 | 6.2 | 2.4 | 2.9 | 4.7 | R | CGA | 2 |
| 146 | Q | CAG | 1.2 | 3.8 | 1.4 | 2.1 | 6.2 | 2.4 | 2.9 | 4.7 | R | CGG | 1 |
| 146 | Q | CAG | 1.2 | 3.8 | 1.4 | 2.1 | 6.2 | 2.4 | 2.9 | 4.7 | R | AGA | 5.1 |
| 146 | Q | CAG | 1.2 | 3.8 | 1.4 | 2.1 | 6.2 | 2.4 | 2.9 | 4.7 | R | AGG | 3.7 |
| 147 | S | AGT | 96.3 | 94.4 | 96.8 | 93 | 18.4 | 93.5 | 97.5 | 19.6 | G | GGT | 1 |
| 147 | S | AGT | 96.3 | 94.4 | 96.8 | 93 | 18.4 | 93.5 | 97.5 | 19.6 | G | GGC | 4.4 |
| 147 | S | AGT | 96.3 | 94.4 | 96.8 | 93 | 18.4 | 93.5 | 97.5 | 19.6 | G | GGA | 6.5 |
| 147 | S | AGT | 96.3 | 94.4 | 96.8 | 93 | 18.4 | 93.5 | 97.5 | 19.6 | G | GGG | 43.2 |

| | | | | | | | | | | | | |
|---|---|---|---|---|---|---|---|---|---|---|---|---|
| 147 | S | AGC | 3.7 | 5.6 | 3.1 | 6.8 | 81.6 | 6.5 | 2.5 | 79.8 | G | GGT | 2.9 |
| 147 | S | AGC | 3.7 | 5.6 | 3.1 | 6.8 | 81.6 | 6.5 | 2.5 | 79.8 | G | GGC | 1 |
| 147 | S | AGC | 3.7 | 5.6 | 3.1 | 6.8 | 81.6 | 6.5 | 2.5 | 79.8 | G | GGA | 5 |
| 147 | S | AGC | 3.7 | 5.6 | 3.1 | 6.8 | 81.6 | 6.5 | 2.5 | 79.8 | G | GGG | 4.4 |
| 148 | Q | CAA | 98 | 88.5 | 7.8 | 97.5 | 99.2 | 95.3 | 95.5 | 97.5 | E | GAA | 3.6 |
| 148 | Q | CAA | 98 | 88.5 | 7.8 | 97.5 | 99.2 | 95.3 | 95.5 | 97.5 | E | GAG | 6.1 |
| 148 | Q | CAG | 2 | 11.5 | 92.2 | 2.4 | 0.6 | 4.7 | 4.5 | 1.9 | E | GAA | 4.6 |
| 148 | Q | CAG | 2 | 11.5 | 92.2 | 2.4 | 0.6 | 4.7 | 4.5 | 1.9 | E | GAG | 3.6 |
| 148 | Q | CAA | 98 | 88.5 | 7.8 | 97.5 | 99.2 | 95.3 | 95.5 | 97.5 | H | CAT | 7.4 |
| 148 | Q | CAA | 98 | 88.5 | 7.8 | 97.5 | 99.2 | 95.3 | 95.5 | 97.5 | H | CAC | 1.8 |
| 148 | Q | CAG | 2 | 11.5 | 92.2 | 2.4 | 0.6 | 4.7 | 4.5 | 1.9 | H | CAT | 13.5 |
| 148 | Q | CAG | 2 | 11.5 | 92.2 | 2.4 | 0.6 | 4.7 | 4.5 | 1.9 | H | CAC | 5.8 |
| 148 | Q | CAA | 98 | 88.5 | 7.8 | 97.5 | 99.2 | 95.3 | 95.5 | 97.5 | G | GGT | 14.4 |
| 148 | Q | CAA | 98 | 88.5 | 7.8 | 97.5 | 99.2 | 95.3 | 95.5 | 97.5 | G | GGC | 11.8 |
| 148 | Q | CAA | 98 | 88.5 | 7.8 | 97.5 | 99.2 | 95.3 | 95.5 | 97.5 | G | GGA | 6.1 |
| 148 | Q | CAA | 98 | 88.5 | 7.8 | 97.5 | 99.2 | 95.3 | 95.5 | 97.5 | G | GGG | 9.4 |
| 148 | Q | CAG | 2 | 11.5 | 92.2 | 2.4 | 0.6 | 4.7 | 4.5 | 1.9 | G | GGT | 51.8 |
| 148 | Q | CAG | 2 | 11.5 | 92.2 | 2.4 | 0.6 | 4.7 | 4.5 | 1.9 | G | GGC | 44.3 |
| 148 | Q | CAG | 2 | 11.5 | 92.2 | 2.4 | 0.6 | 4.7 | 4.5 | 1.9 | G | GGA | 7.5 |
| 148 | Q | CAG | 2 | 11.5 | 92.2 | 2.4 | 0.6 | 4.7 | 4.5 | 1.9 | G | GGG | 6.1 |
| 148 | Q | CAA | 98 | 88.5 | 7.8 | 97.5 | 99.2 | 95.3 | 95.5 | 97.5 | K | AAA | 1.2 |
| 148 | Q | CAA | 98 | 88.5 | 7.8 | 97.5 | 99.2 | 95.3 | 95.5 | 97.5 | K | AAG | 3.7 |
| 148 | Q | CAG | 2 | 11.5 | 92.2 | 2.4 | 0.6 | 4.7 | 4.5 | 1.9 | K | AAA | 2.2 |
| 148 | Q | CAG | 2 | 11.5 | 92.2 | 2.4 | 0.6 | 4.7 | 4.5 | 1.9 | K | AAG | 1.2 |
| 148 | Q | CAA | 98 | 88.5 | 7.8 | 97.5 | 99.2 | 95.3 | 95.5 | 97.5 | R | CGT | 7.4 |
| 148 | Q | CAA | 98 | 88.5 | 7.8 | 97.5 | 99.2 | 95.3 | 95.5 | 97.5 | R | CGC | 9.2 |
| 148 | Q | CAA | 98 | 88.5 | 7.8 | 97.5 | 99.2 | 95.3 | 95.5 | 97.5 | R | CGA | 1 |
| 148 | Q | CAA | 98 | 88.5 | 7.8 | 97.5 | 99.2 | 95.3 | 95.5 | 97.5 | R | CGG | 3.5 |
| 148 | Q | CAA | 98 | 88.5 | 7.8 | 97.5 | 99.2 | 95.3 | 95.5 | 97.5 | R | AGA | 3.7 |
| 148 | Q | CAA | 98 | 88.5 | 7.8 | 97.5 | 99.2 | 95.3 | 95.5 | 97.5 | R | AGG | 7 |
| 148 | Q | CAG | 2 | 11.5 | 92.2 | 2.4 | 0.6 | 4.7 | 4.5 | 1.9 | R | CGT | 21.2 |
| 148 | Q | CAG | 2 | 11.5 | 92.2 | 2.4 | 0.6 | 4.7 | 4.5 | 1.9 | R | CGC | 3.9 |
| 148 | Q | CAG | 2 | 11.5 | 92.2 | 2.4 | 0.6 | 4.7 | 4.5 | 1.9 | R | CGA | 2 |
| 148 | Q | CAG | 2 | 11.5 | 92.2 | 2.4 | 0.6 | 4.7 | 4.5 | 1.9 | R | CGG | 1 |

| | | | | | | | | | | | | |
|---|---|---|---|---|---|---|---|---|---|---|---|---|
| 148 | Q | CAG | 2 | 11.5 | 92.2 | 2.4 | 0.6 | 4.7 | 4.5 | 1.9 | R | AGA | 5.1 |
| 148 | Q | CAG | 2 | 11.5 | 92.2 | 2.4 | 0.6 | 4.7 | 4.5 | 1.9 | R | AGG | 3.7 |
| 148 | Q | CAA | 98 | 88.5 | 7.8 | 97.5 | 99.2 | 95.3 | 95.5 | 97.5 | N | AAT | 7.6 |
| 148 | Q | CAA | 98 | 88.5 | 7.8 | 97.5 | 99.2 | 95.3 | 95.5 | 97.5 | N | AAC | 9.4 |
| 148 | Q | CAG | 2 | 11.5 | 92.2 | 2.4 | 0.6 | 4.7 | 4.5 | 1.9 | N | AAT | 21.4 |
| 148 | Q | CAG | 2 | 11.5 | 92.2 | 2.4 | 0.6 | 4.7 | 4.5 | 1.9 | N | AAC | 4.1 |
| 149 | G | GGA | 98.8 | 90.2 | 99.2 | 96.5 | 97.9 | 80.6 | 97.7 | 99.4 | A | GCT | 12.2 |
| 149 | G | GGA | 98.8 | 90.2 | 99.2 | 96.5 | 97.9 | 80.6 | 97.7 | 99.4 | A | GCC | 14 |
| 149 | G | GGA | 98.8 | 90.2 | 99.2 | 96.5 | 97.9 | 80.6 | 97.7 | 99.4 | A | GCA | 5.8 |
| 149 | G | GGA | 98.8 | 90.2 | 99.2 | 96.5 | 97.9 | 80.6 | 97.7 | 99.4 | A | GCG | 8.3 |
| 149 | G | GGG | 0.7 | 6.8 | 0.5 | 2.2 | 1.6 | 16.8 | 1.2 | 0.6 | A | GCT | 26 |
| 149 | G | GGG | 0.7 | 6.8 | 0.5 | 2.2 | 1.6 | 16.8 | 1.2 | 0.6 | A | GCC | 8.7 |
| 149 | G | GGG | 0.7 | 6.8 | 0.5 | 2.2 | 1.6 | 16.8 | 1.2 | 0.6 | A | GCA | 6.8 |
| 149 | G | GGG | 0.7 | 6.8 | 0.5 | 2.2 | 1.6 | 16.8 | 1.2 | 0.6 | A | GCG | 5.8 |
| 149 | G | GGC | 0.5 | 2.4 | 0.2 | 0.1 | 0 | 1.8 | 0 | 0 | A | GCT | 7.7 |
| 149 | G | GGC | 0.5 | 2.4 | 0.2 | 0.1 | 0 | 1.8 | 0 | 0 | A | GCC | 5.8 |
| 149 | G | GGC | 0.5 | 2.4 | 0.2 | 0.1 | 0 | 1.8 | 0 | 0 | A | GCA | 9.8 |
| 149 | G | GGC | 0.5 | 2.4 | 0.2 | 0.1 | 0 | 1.8 | 0 | 0 | A | GCG | 9.2 |
| 149 | G | GGT | 0 | 0.4 | 0.1 | 1.2 | 0.3 | 0.9 | 1.2 | 0 | A | GCT | 5.8 |
| 149 | G | GGT | 0 | 0.4 | 0.1 | 1.2 | 0.3 | 0.9 | 1.2 | 0 | A | GCC | 9.2 |
| 149 | G | GGT | 0 | 0.4 | 0.1 | 1.2 | 0.3 | 0.9 | 1.2 | 0 | A | GCA | 11.3 |
| 149 | G | GGT | 0 | 0.4 | 0.1 | 1.2 | 0.3 | 0.9 | 1.2 | 0 | A | GCG | 48 |
| 151 | V | GTA | 16.5 | 88.8 | 98.9 | 90.9 | 12.2 | 83.2 | 96.1 | 18.6 | A | GCT | 8 |
| 151 | V | GTA | 16.5 | 88.8 | 98.9 | 90.9 | 12.2 | 83.2 | 96.1 | 18.6 | A | GCC | 9.8 |
| 151 | V | GTA | 16.5 | 88.8 | 98.9 | 90.9 | 12.2 | 83.2 | 96.1 | 18.6 | A | GCA | 1.6 |
| 151 | V | GTA | 16.5 | 88.8 | 98.9 | 90.9 | 12.2 | 83.2 | 96.1 | 18.6 | A | GCG | 4.1 |
| 151 | V | GTG | 83.5 | 5.6 | 0.7 | 6.8 | 87.3 | 16.2 | 3.1 | 80.7 | A | GCT | 21.8 |
| 151 | V | GTG | 83.5 | 5.6 | 0.7 | 6.8 | 87.3 | 16.2 | 3.1 | 80.7 | A | GCC | 4.5 |
| 151 | V | GTG | 83.5 | 5.6 | 0.7 | 6.8 | 87.3 | 16.2 | 3.1 | 80.7 | A | GCA | 2.6 |
| 151 | V | GTG | 83.5 | 5.6 | 0.7 | 6.8 | 87.3 | 16.2 | 3.1 | 80.7 | A | GCG | 1.6 |
| 151 | I | ATA | 0 | 5.2 | 0.3 | 1.5 | 0.3 | 0 | 0.8 | 0.6 | A | GCT | 12.7 |
| 151 | I | ATA | 0 | 5.2 | 0.3 | 1.5 | 0.3 | 0 | 0.8 | 0.6 | A | GCC | 10.1 |
| 151 | I | ATA | 0 | 5.2 | 0.3 | 1.5 | 0.3 | 0 | 0.8 | 0.6 | A | GCA | 4.4 |
| 151 | I | ATA | 0 | 5.2 | 0.3 | 1.5 | 0.3 | 0 | 0.8 | 0.6 | A | GCG | 7.7 |

| | | | | | | | | | | | | | |
|---|---|---|---|---|---|---|---|---|---|---|---|---|---|
| 151 | V | GTA | 16.5 | 88.8 | 98.9 | 90.9 | 12.2 | 83.2 | 96.1 | 18.6 | I | ATT | 7.4 |
| 151 | V | GTA | 16.5 | 88.8 | 98.9 | 90.9 | 12.2 | 83.2 | 96.1 | 18.6 | I | ATC | 9.2 |
| 151 | V | GTA | 16.5 | 88.8 | 98.9 | 90.9 | 12.2 | 83.2 | 96.1 | 18.6 | I | ATA | 1 |
| 151 | V | GTG | 83.5 | 5.6 | 0.7 | 6.8 | 87.3 | 16.2 | 3.1 | 80.7 | I | ATT | 21.2 |
| 151 | V | GTG | 83.5 | 5.6 | 0.7 | 6.8 | 87.3 | 16.2 | 3.1 | 80.7 | I | ATC | 3.9 |
| 151 | V | GTG | 83.5 | 5.6 | 0.7 | 6.8 | 87.3 | 16.2 | 3.1 | 80.7 | I | ATA | 2 |
| 151 | I | ATA | 0 | 5.2 | 0.3 | 1.5 | 0.3 | 0 | 0.8 | 0.6 | I | ATT | 7.4 |
| 151 | I | ATA | 0 | 5.2 | 0.3 | 1.5 | 0.3 | 0 | 0.8 | 0.6 | I | ATC | 1.8 |
| 151 | I | ATA | 0 | 5.2 | 0.3 | 1.5 | 0.3 | 0 | 0.8 | 0.6 | I | ATA | 0 |
| 151 | V | GTA | 16.5 | 88.8 | 98.9 | 90.9 | 12.2 | 83.2 | 96.1 | 18.6 | L | TTA | 13.5 |
| 151 | V | GTA | 16.5 | 88.8 | 98.9 | 90.9 | 12.2 | 83.2 | 96.1 | 18.6 | L | TTG | 16 |
| 151 | V | GTA | 16.5 | 88.8 | 98.9 | 90.9 | 12.2 | 83.2 | 96.1 | 18.6 | L | CTT | 12.2 |
| 151 | V | GTA | 16.5 | 88.8 | 98.9 | 90.9 | 12.2 | 83.2 | 96.1 | 18.6 | L | CTC | 14 |
| 151 | V | GTA | 16.5 | 88.8 | 98.9 | 90.9 | 12.2 | 83.2 | 96.1 | 18.6 | L | CTA | 5.8 |
| 151 | V | GTA | 16.5 | 88.8 | 98.9 | 90.9 | 12.2 | 83.2 | 96.1 | 18.6 | L | CTG | 8.3 |
| 151 | V | GTG | 83.5 | 5.6 | 0.7 | 6.8 | 87.3 | 16.2 | 3.1 | 80.7 | L | TTA | 14.5 |
| 151 | V | GTG | 83.5 | 5.6 | 0.7 | 6.8 | 87.3 | 16.2 | 3.1 | 80.7 | L | TTG | 13.5 |
| 151 | V | GTG | 83.5 | 5.6 | 0.7 | 6.8 | 87.3 | 16.2 | 3.1 | 80.7 | L | CTT | 26 |
| 151 | V | GTG | 83.5 | 5.6 | 0.7 | 6.8 | 87.3 | 16.2 | 3.1 | 80.7 | L | CTC | 8.7 |
| 151 | V | GTG | 83.5 | 5.6 | 0.7 | 6.8 | 87.3 | 16.2 | 3.1 | 80.7 | L | CTA | 6.8 |
| 151 | V | GTG | 83.5 | 5.6 | 0.7 | 6.8 | 87.3 | 16.2 | 3.1 | 80.7 | L | CTG | 5.8 |
| 151 | I | ATA | 0 | 5.2 | 0.3 | 1.5 | 0.3 | 0 | 0.8 | 0.6 | L | TTA | 7.4 |
| 151 | I | ATA | 0 | 5.2 | 0.3 | 1.5 | 0.3 | 0 | 0.8 | 0.6 | L | TTG | 9.9 |
| 151 | I | ATA | 0 | 5.2 | 0.3 | 1.5 | 0.3 | 0 | 0.8 | 0.6 | L | CTT | 8.2 |
| 151 | I | ATA | 0 | 5.2 | 0.3 | 1.5 | 0.3 | 0 | 0.8 | 0.6 | L | CTC | 10 |
| 151 | I | ATA | 0 | 5.2 | 0.3 | 1.5 | 0.3 | 0 | 0.8 | 0.6 | L | CTA | 1.8 |
| 151 | I | ATA | 0 | 5.2 | 0.3 | 1.5 | 0.3 | 0 | 0.8 | 0.6 | L | CTG | 4.3 |
| 153 | S | TCT | 89.1 | 86.4 | 6.2 | 78.7 | 91.5 | 90.2 | 91.1 | 92.2 | F | TTT | 2.6 |
| 153 | S | TCT | 89.1 | 86.4 | 6.2 | 78.7 | 91.5 | 90.2 | 91.1 | 92.2 | F | TTC | 6 |
| 153 | S | TCC | 7.9 | 8.6 | 85.6 | 19.1 | 6 | 6.8 | 6.2 | 5.3 | F | TTT | 4.5 |
| 153 | S | TCC | 7.9 | 8.6 | 85.6 | 19.1 | 6 | 6.8 | 6.2 | 5.3 | F | TTC | 2.6 |
| 153 | S | TCA | 1.7 | 2.1 | 7.9 | 1.5 | 1.4 | 1.2 | 1.6 | 1.9 | F | TTT | 9 |
| 153 | S | TCA | 1.7 | 2.1 | 7.9 | 1.5 | 1.4 | 1.2 | 1.6 | 1.9 | F | TTC | 10.8 |
| 153 | S | AGC | 0.2 | 2.4 | 0.1 | 0 | 0 | 1.8 | 0 | 0 | F | TTT | 28.6 |

| | | | | | | | | | | | | | |
|---|---|---|---|---|---|---|---|---|---|---|---|---|---|
| 153 | S | AGC | 0.2 | 2.4 | 0.1 | 0 | 0 | 1.8 | 0 | 0 | F | TTC | 27.6 |
| 153 | S | TCT | 89.1 | 86.4 | 6.2 | 78.7 | 91.5 | 90.2 | 91.1 | 92.2 | Y | TAT | 1.2 |
| 153 | S | TCT | 89.1 | 86.4 | 6.2 | 78.7 | 91.5 | 90.2 | 91.1 | 92.2 | Y | TAC | 4.6 |
| 153 | S | TCC | 7.9 | 8.6 | 85.6 | 19.1 | 6 | 6.8 | 6.2 | 5.3 | Y | TAT | 3.1 |
| 153 | S | TCC | 7.9 | 8.6 | 85.6 | 19.1 | 6 | 6.8 | 6.2 | 5.3 | Y | TAC | 1.2 |
| 153 | S | TCA | 1.7 | 2.1 | 7.9 | 1.5 | 1.4 | 1.2 | 1.6 | 1.9 | Y | TAT | 7.6 |
| 153 | S | TCA | 1.7 | 2.1 | 7.9 | 1.5 | 1.4 | 1.2 | 1.6 | 1.9 | Y | TAC | 9.4 |
| 153 | S | AGC | 0.2 | 2.4 | 0.1 | 0 | 0 | 1.8 | 0 | 0 | Y | TAT | 9.4 |
| 153 | S | AGC | 0.2 | 2.4 | 0.1 | 0 | 0 | 1.8 | 0 | 0 | Y | TAC | 8.4 |
| 154 | M | ATG | 99.5 | 95.5 | 99.5 | 96.4 | 98.9 | 97.6 | 96.5 | 99.7 | I | ATT | 13.5 |
| 154 | M | ATG | 99.5 | 95.5 | 99.5 | 96.4 | 98.9 | 97.6 | 96.5 | 99.7 | I | ATC | 5.8 |
| 154 | M | ATG | 99.5 | 95.5 | 99.5 | 96.4 | 98.9 | 97.6 | 96.5 | 99.7 | I | ATA | 1 |
| 154 | I | ATA | 0.2 | 2.4 | 0.2 | 3 | 0.9 | 1.5 | 3.1 | 0 | I | ATT | 7.4 |
| 154 | I | ATA | 0.2 | 2.4 | 0.2 | 3 | 0.9 | 1.5 | 3.1 | 0 | I | ATC | 1.8 |
| 154 | I | ATA | 0.2 | 2.4 | 0.2 | 3 | 0.9 | 1.5 | 3.1 | 0 | I | ATA | 0 |
| 154 | L | CTA | 0.2 | 1.6 | 0 | 0.1 | 0 | 0 | 0.4 | 0 | I | ATT | 7.6 |
| 154 | L | CTA | 0.2 | 1.6 | 0 | 0.1 | 0 | 0 | 0.4 | 0 | I | ATC | 9.4 |
| 154 | L | CTA | 0.2 | 1.6 | 0 | 0.1 | 0 | 0 | 0.4 | 0 | I | ATA | 1.2 |
| 155 | N | AAT | 96.2 | 94.7 | 97.6 | 98.2 | 97.8 | 94.1 | 95.5 | 98.8 | H | CAT | 1.8 |
| 155 | N | AAT | 96.2 | 94.7 | 97.6 | 98.2 | 97.8 | 94.1 | 95.5 | 98.8 | H | CAC | 5.2 |
| 155 | N | AAC | 3.8 | 5.3 | 2.4 | 1.7 | 2.2 | 5.9 | 4.5 | 1.2 | H | CAT | 3.7 |
| 155 | N | AAC | 3.8 | 5.3 | 2.4 | 1.7 | 2.2 | 5.9 | 4.5 | 1.2 | H | CAC | 1.8 |
| 155 | N | AAT | 96.2 | 94.7 | 97.6 | 98.2 | 97.8 | 94.1 | 95.5 | 98.8 | S | TCT | 15.6 |
| 155 | N | AAT | 96.2 | 94.7 | 97.6 | 98.2 | 97.8 | 94.1 | 95.5 | 98.8 | S | TCC | 17.6 |
| 155 | N | AAT | 96.2 | 94.7 | 97.6 | 98.2 | 97.8 | 94.1 | 95.5 | 98.8 | S | TCA | 19.5 |
| 155 | N | AAT | 96.2 | 94.7 | 97.6 | 98.2 | 97.8 | 94.1 | 95.5 | 98.8 | S | TCG | 66.8 |
| 155 | N | AAT | 96.2 | 94.7 | 97.6 | 98.2 | 97.8 | 94.1 | 95.5 | 98.8 | S | AGT | 1 |
| 155 | N | AAT | 96.2 | 94.7 | 97.6 | 98.2 | 97.8 | 94.1 | 95.5 | 98.8 | S | AGC | 4.4 |
| 155 | N | AAC | 3.8 | 5.3 | 2.4 | 1.7 | 2.2 | 5.9 | 4.5 | 1.2 | S | TCT | 16.6 |
| 155 | N | AAC | 3.8 | 5.3 | 2.4 | 1.7 | 2.2 | 5.9 | 4.5 | 1.2 | S | TCC | 15.6 |
| 155 | N | AAC | 3.8 | 5.3 | 2.4 | 1.7 | 2.2 | 5.9 | 4.5 | 1.2 | S | TCA | 16.9 |
| 155 | N | AAC | 3.8 | 5.3 | 2.4 | 1.7 | 2.2 | 5.9 | 4.5 | 1.2 | S | TCG | 36.6 |
| 155 | N | AAC | 3.8 | 5.3 | 2.4 | 1.7 | 2.2 | 5.9 | 4.5 | 1.2 | S | AGT | 2.9 |
| 155 | N | AAC | 3.8 | 5.3 | 2.4 | 1.7 | 2.2 | 5.9 | 4.5 | 1.2 | S | AGC | 1 |

| | | | | | | | | | | | | |
|---|---|---|---|---|---|---|---|---|---|---|---|---|
| 155 | N | AAT | 96.2 | 94.7 | 97.6 | 98.2 | 97.8 | 94.1 | 95.5 | 98.8 | T | ACT | 1.8 |
| 155 | N | AAT | 96.2 | 94.7 | 97.6 | 98.2 | 97.8 | 94.1 | 95.5 | 98.8 | T | ACC | 5.2 |
| 155 | N | AAT | 96.2 | 94.7 | 97.6 | 98.2 | 97.8 | 94.1 | 95.5 | 98.8 | T | ACA | 7.3 |
| 155 | N | AAT | 96.2 | 94.7 | 97.6 | 98.2 | 97.8 | 94.1 | 95.5 | 98.8 | T | ACG | 44 |
| 155 | N | AAC | 3.8 | 5.3 | 2.4 | 1.7 | 2.2 | 5.9 | 4.5 | 1.2 | T | ACT | 3.7 |
| 155 | N | AAC | 3.8 | 5.3 | 2.4 | 1.7 | 2.2 | 5.9 | 4.5 | 1.2 | T | ACC | 1.8 |
| 155 | N | AAC | 3.8 | 5.3 | 2.4 | 1.7 | 2.2 | 5.9 | 4.5 | 1.2 | T | ACA | 5.8 |
| 155 | N | AAC | 3.8 | 5.3 | 2.4 | 1.7 | 2.2 | 5.9 | 4.5 | 1.2 | T | ACG | 5.2 |
| 156 | K | AAA | 24.8 | 76.4 | 92.6 | 42.6 | 91.2 | 89.1 | 92.2 | 85.4 | N | AAT | 7.4 |
| 156 | K | AAA | 24.8 | 76.4 | 92.6 | 42.6 | 91.2 | 89.1 | 92.2 | 85.4 | N | AAC | 1.8 |
| 156 | K | AAG | 74.4 | 4.7 | 6.6 | 57.2 | 8.6 | 7.6 | 4.7 | 14 | N | AAT | 13.5 |
| 156 | K | AAG | 74.4 | 4.7 | 6.6 | 57.2 | 8.6 | 7.6 | 4.7 | 14 | N | AAC | 5.8 |
| 156 | N | AAT | 0.5 | 17.6 | 0.3 | 0.1 | 0.2 | 1.5 | 2.3 | 0 | N | AAT | 0 |
| 156 | N | AAT | 0.5 | 17.6 | 0.3 | 0.1 | 0.2 | 1.5 | 2.3 | 0 | N | AAC | 1.6 |
| 157 | E | GAA | 94.1 | 93.6 | 94.8 | 96 | 87 | 91.5 | 43.6 | 95.3 | Q | CAA | 5.8 |
| 157 | E | GAA | 94.1 | 93.6 | 94.8 | 96 | 87 | 91.5 | 43.6 | 95.3 | Q | CAG | 8.3 |
| 157 | E | GAG | 4 | 3.5 | 4.3 | 2.4 | 3.9 | 3.2 | 55.8 | 3.1 | Q | CAA | 6.8 |
| 157 | E | GAG | 4 | 3.5 | 4.3 | 2.4 | 3.9 | 3.2 | 55.8 | 3.1 | Q | CAG | 5.8 |
| 157 | Q | CAA | 1.7 | 2.3 | 0.4 | 1.1 | 8.3 | 3.5 | 0 | 0.9 | Q | CAA | 0 |
| 157 | Q | CAA | 1.7 | 2.3 | 0.4 | 1.1 | 8.3 | 3.5 | 0 | 0.9 | Q | CAG | 1 |
| 160 | K | AAA | 95 | 93 | 94.5 | 92.8 | 92.9 | 88.2 | 48.4 | 96.3 | N | AAT | 7.4 |
| 160 | K | AAA | 95 | 93 | 94.5 | 92.8 | 92.9 | 88.2 | 48.4 | 96.3 | N | AAC | 1.8 |
| 160 | K | AAG | 3.2 | 4 | 2.6 | 2.9 | 2.3 | 6.2 | 51.2 | 3.1 | N | AAT | 13.5 |
| 160 | K | AAG | 3.2 | 4 | 2.6 | 2.9 | 2.3 | 6.2 | 51.2 | 3.1 | N | AAC | 5.8 |
| 160 | Q | CAA | 1 | 1.6 | 0.5 | 1 | 3.9 | 1.5 | 0 | 0.6 | N | AAT | 7.6 |
| 160 | Q | CAA | 1 | 1.6 | 0.5 | 1 | 3.9 | 1.5 | 0 | 0.6 | N | AAC | 9.4 |
| 160 | R | AGA | 0.9 | 0.3 | 1.8 | 2.4 | 0.3 | 2.1 | 0 | 0 | N | AAT | 7.4 |
| 160 | R | AGA | 0.9 | 0.3 | 1.8 | 2.4 | 0.3 | 2.1 | 0 | 0 | N | AAC | 9.2 |
| 160 | N | AAC | 0 | 0.2 | 0.1 | 0.2 | 0.2 | 1.2 | 0 | 0 | N | AAT | 2.6 |
| 160 | N | AAC | 0 | 0.2 | 0.1 | 0.2 | 0.2 | 1.2 | 0 | 0 | N | AAC | 0 |
| 163 | G | GGA | 24.3 | 77.5 | 19.3 | 9.5 | 12.4 | 79.4 | 63.8 | 13.4 | K | AAA | 2 |
| 163 | G | GGA | 24.3 | 77.5 | 19.3 | 9.5 | 12.4 | 79.4 | 63.8 | 13.4 | K | AAG | 5.3 |
| 163 | G | GGG | 72.9 | 11.3 | 75.4 | 88.7 | 83.1 | 10.6 | 8.4 | 79.2 | K | AAA | 3.4 |
| 163 | G | GGG | 72.9 | 11.3 | 75.4 | 88.7 | 83.1 | 10.6 | 8.4 | 79.2 | K | AAG | 2 |

| | | | | | | | | | | | | |
|---|---|---|---|---|---|---|---|---|---|---|---|---|
| 163 | E | GAA | 0 | 5.1 | 0.2 | 0.1 | 0.2 | 2.1 | 2.9 | 0.6 | K | AAA | 1 |
| 163 | E | GAA | 0 | 5.1 | 0.2 | 0.1 | 0.2 | 2.1 | 2.9 | 0.6 | K | AAG | 3.5 |
| 163 | G | GGC | 0.7 | 3 | 0.1 | 0 | 0 | 1.8 | 0.6 | 0 | K | AAA | 3.3 |
| 163 | G | GGC | 0.7 | 3 | 0.1 | 0 | 0 | 1.8 | 0.6 | 0 | K | AAG | 23 |
| 163 | E | GAG | 0.9 | 0.5 | 2.4 | 1.2 | 2.4 | 0.6 | 0.8 | 1.2 | K | AAA | 2 |
| 163 | E | GAG | 0.9 | 0.5 | 2.4 | 1.2 | 2.4 | 0.6 | 0.8 | 1.2 | K | AAG | 1 |
| 163 | A | GCA | 0.2 | 0.5 | 0.2 | 0.1 | 0.5 | 0.6 | 1.4 | 0.6 | K | AAA | 5 |
| 163 | A | GCA | 0.2 | 0.5 | 0.2 | 0.1 | 0.5 | 0.6 | 1.4 | 0.6 | K | AAG | 8.3 |
| 163 | R | AGA | 0.1 | 0.2 | 0 | 0 | 0 | 0.6 | 6.2 | 0 | K | AAA | 1 |
| 163 | R | AGA | 0.1 | 0.2 | 0 | 0 | 0 | 0.6 | 6.2 | 0 | K | AAG | 3.5 |
| 163 | K | AAA | 0.1 | 0 | 0 | 0 | 0 | 0 | 5.4 | 0 | K | AAA | 0 |
| 163 | K | AAA | 0.1 | 0 | 0 | 0 | 0 | 0 | 5.4 | 0 | K | AAG | 1 |
| 163 | T | ACA | 0.2 | 0.3 | 0.1 | 0 | 0 | 2.9 | 1.8 | 0 | K | AAA | 1.2 |
| 163 | T | ACA | 0.2 | 0.3 | 0.1 | 0 | 0 | 2.9 | 1.8 | 0 | K | AAG | 3.7 |
| 163 | S | AGC | 0 | 0 | 0.1 | 0 | 0 | 0 | 2.5 | 0 | K | AAA | 5 |
| 163 | S | AGC | 0 | 0 | 0.1 | 0 | 0 | 0 | 2.5 | 0 | K | AAG | 4.4 |
| 163 | S | AGT | 0 | 0 | 0 | 0 | 0 | 0 | 2.9 | 0 | K | AAA | 6.5 |
| 163 | S | AGT | 0 | 0 | 0 | 0 | 0 | 0 | 2.9 | 0 | K | AAG | 43.2 |
| 163 | A | GCG | 0 | 0 | 0 | 0.1 | 0.4 | 0 | 0 | 1.9 | K | AAA | 6.4 |
| 163 | A | GCG | 0 | 0 | 0 | 0.1 | 0.4 | 0 | 0 | 1.9 | K | AAG | 5 |
| 163 | M | ATG | 0 | 0 | 0.2 | 0 | 0 | 0 | 0 | 1.2 | K | AAA | 4 |
| 163 | M | ATG | 0 | 0 | 0.2 | 0 | 0 | 0 | 0 | 1.2 | K | AAG | 3 |
| 163 | G | GGA | 24.3 | 77.5 | 19.3 | 9.5 | 12.4 | 79.4 | 63.8 | 13.4 | R | CGT | 12.2 |
| 163 | G | GGA | 24.3 | 77.5 | 19.3 | 9.5 | 12.4 | 79.4 | 63.8 | 13.4 | R | CGC | 14 |
| 163 | G | GGA | 24.3 | 77.5 | 19.3 | 9.5 | 12.4 | 79.4 | 63.8 | 13.4 | R | CGA | 5.8 |
| 163 | G | GGA | 24.3 | 77.5 | 19.3 | 9.5 | 12.4 | 79.4 | 63.8 | 13.4 | R | CGG | 8.3 |
| 163 | G | GGA | 24.3 | 77.5 | 19.3 | 9.5 | 12.4 | 79.4 | 63.8 | 13.4 | R | AGA | 1 |
| 163 | G | GGA | 24.3 | 77.5 | 19.3 | 9.5 | 12.4 | 79.4 | 63.8 | 13.4 | R | AGG | 3.5 |
| 163 | G | GGG | 72.9 | 11.3 | 75.4 | 88.7 | 83.1 | 10.6 | 8.4 | 79.2 | R | CGT | 26 |
| 163 | G | GGG | 72.9 | 11.3 | 75.4 | 88.7 | 83.1 | 10.6 | 8.4 | 79.2 | R | CGC | 8.7 |
| 163 | G | GGG | 72.9 | 11.3 | 75.4 | 88.7 | 83.1 | 10.6 | 8.4 | 79.2 | R | CGA | 6.8 |
| 163 | G | GGG | 72.9 | 11.3 | 75.4 | 88.7 | 83.1 | 10.6 | 8.4 | 79.2 | R | CGG | 5.8 |
| 163 | G | GGG | 72.9 | 11.3 | 75.4 | 88.7 | 83.1 | 10.6 | 8.4 | 79.2 | R | AGA | 2 |
| 163 | G | GGG | 72.9 | 11.3 | 75.4 | 88.7 | 83.1 | 10.6 | 8.4 | 79.2 | R | AGG | 1 |

| | | | | | | | | | | | | | |
|---|---|---|---|---|---|---|---|---|---|---|---|---|---|
| 163 | E | GAA | 0 | 5.1 | 0.2 | 0.1 | 0.2 | 2.1 | 2.9 | 0.6 | R | CGT | 16.6 |
| 163 | E | GAA | 0 | 5.1 | 0.2 | 0.1 | 0.2 | 2.1 | 2.9 | 0.6 | R | CGC | 14 |
| 163 | E | GAA | 0 | 5.1 | 0.2 | 0.1 | 0.2 | 2.1 | 2.9 | 0.6 | R | CGA | 8.3 |
| 163 | E | GAA | 0 | 5.1 | 0.2 | 0.1 | 0.2 | 2.1 | 2.9 | 0.6 | R | CGG | 11.6 |
| 163 | E | GAA | 0 | 5.1 | 0.2 | 0.1 | 0.2 | 2.1 | 2.9 | 0.6 | R | AGA | 3.5 |
| 163 | E | GAA | 0 | 5.1 | 0.2 | 0.1 | 0.2 | 2.1 | 2.9 | 0.6 | R | AGG | 6.8 |
| 163 | G | GGC | 0.7 | 3 | 0.1 | 0 | 0 | 1.8 | 0.6 | 0 | R | CGT | 7.7 |
| 163 | G | GGC | 0.7 | 3 | 0.1 | 0 | 0 | 1.8 | 0.6 | 0 | R | CGC | 5.8 |
| 163 | G | GGC | 0.7 | 3 | 0.1 | 0 | 0 | 1.8 | 0.6 | 0 | R | CGA | 9.8 |
| 163 | G | GGC | 0.7 | 3 | 0.1 | 0 | 0 | 1.8 | 0.6 | 0 | R | CGG | 9.2 |
| 163 | G | GGC | 0.7 | 3 | 0.1 | 0 | 0 | 1.8 | 0.6 | 0 | R | AGA | 5 |
| 163 | G | GGC | 0.7 | 3 | 0.1 | 0 | 0 | 1.8 | 0.6 | 0 | R | AGG | 4.4 |
| 163 | E | GAG | 0.9 | 0.5 | 2.4 | 1.2 | 2.4 | 0.6 | 0.8 | 1.2 | R | CGT | 54 |
| 163 | E | GAG | 0.9 | 0.5 | 2.4 | 1.2 | 2.4 | 0.6 | 0.8 | 1.2 | R | CGC | 46.5 |
| 163 | E | GAG | 0.9 | 0.5 | 2.4 | 1.2 | 2.4 | 0.6 | 0.8 | 1.2 | R | CGA | 9.7 |
| 163 | E | GAG | 0.9 | 0.5 | 2.4 | 1.2 | 2.4 | 0.6 | 0.8 | 1.2 | R | CGG | 8.3 |
| 163 | E | GAG | 0.9 | 0.5 | 2.4 | 1.2 | 2.4 | 0.6 | 0.8 | 1.2 | R | AGA | 4.9 |
| 163 | E | GAG | 0.9 | 0.5 | 2.4 | 1.2 | 2.4 | 0.6 | 0.8 | 1.2 | R | AGG | 3.5 |
| 163 | A | GCA | 0.2 | 0.5 | 0.2 | 0.1 | 0.5 | 0.6 | 1.4 | 0.6 | R | CGT | 17.5 |
| 163 | A | GCA | 0.2 | 0.5 | 0.2 | 0.1 | 0.5 | 0.6 | 1.4 | 0.6 | R | CGC | 14.9 |
| 163 | A | GCA | 0.2 | 0.5 | 0.2 | 0.1 | 0.5 | 0.6 | 1.4 | 0.6 | R | CGA | 9.2 |
| 163 | A | GCA | 0.2 | 0.5 | 0.2 | 0.1 | 0.5 | 0.6 | 1.4 | 0.6 | R | CGG | 12.5 |
| 163 | A | GCA | 0.2 | 0.5 | 0.2 | 0.1 | 0.5 | 0.6 | 1.4 | 0.6 | R | AGA | 4.4 |
| 163 | A | GCA | 0.2 | 0.5 | 0.2 | 0.1 | 0.5 | 0.6 | 1.4 | 0.6 | R | AGG | 7.7 |
| 163 | R | AGA | 0.1 | 0.2 | 0 | 0 | 0 | 0.6 | 6.2 | 0 | R | CGT | 8.2 |
| 163 | R | AGA | 0.1 | 0.2 | 0 | 0 | 0 | 0.6 | 6.2 | 0 | R | CGC | 10 |
| 163 | R | AGA | 0.1 | 0.2 | 0 | 0 | 0 | 0.6 | 6.2 | 0 | R | CGA | 1.8 |
| 163 | R | AGA | 0.1 | 0.2 | 0 | 0 | 0 | 0.6 | 6.2 | 0 | R | CGG | 4.3 |
| 163 | R | AGA | 0.1 | 0.2 | 0 | 0 | 0 | 0.6 | 6.2 | 0 | R | AGA | 0 |
| 163 | R | AGA | 0.1 | 0.2 | 0 | 0 | 0 | 0.6 | 6.2 | 0 | R | AGG | 1 |
| 163 | K | AAA | 0.1 | 0 | 0 | 0 | 0 | 0 | 5.4 | 0 | R | CGT | 12.6 |
| 163 | K | AAA | 0.1 | 0 | 0 | 0 | 0 | 0 | 5.4 | 0 | R | CGC | 10 |
| 163 | K | AAA | 0.1 | 0 | 0 | 0 | 0 | 0 | 5.4 | 0 | R | CGA | 4.3 |
| 163 | K | AAA | 0.1 | 0 | 0 | 0 | 0 | 0 | 5.4 | 0 | R | CGG | 7.6 |

| | | | | | | | | | | | | |
|---|---|---|---|---|---|---|---|---|---|---|---|---|
| 163 | K | AAA | 0.1 | 0 | 0 | 0 | 0 | 0 | 5.4 | 0 | R | AGA | 1 |
| 163 | K | AAA | 0.1 | 0 | 0 | 0 | 0 | 0 | 5.4 | 0 | R | AGG | 3.5 |
| 163 | T | ACA | 0.2 | 0.3 | 0.1 | 0 | 0 | 2.9 | 1.8 | 0 | R | CGT | 13.5 |
| 163 | T | ACA | 0.2 | 0.3 | 0.1 | 0 | 0 | 2.9 | 1.8 | 0 | R | CGC | 10.9 |
| 163 | T | ACA | 0.2 | 0.3 | 0.1 | 0 | 0 | 2.9 | 1.8 | 0 | R | CGA | 5.2 |
| 163 | T | ACA | 0.2 | 0.3 | 0.1 | 0 | 0 | 2.9 | 1.8 | 0 | R | CGG | 8.5 |
| 163 | T | ACA | 0.2 | 0.3 | 0.1 | 0 | 0 | 2.9 | 1.8 | 0 | R | AGA | 3.6 |
| 163 | T | ACA | 0.2 | 0.3 | 0.1 | 0 | 0 | 2.9 | 1.8 | 0 | R | AGG | 6.1 |
| 163 | S | AGC | 0 | 0 | 0.1 | 0 | 0 | 0 | 2.5 | 0 | R | CGT | 3.7 |
| 163 | S | AGC | 0 | 0 | 0.1 | 0 | 0 | 0 | 2.5 | 0 | R | CGC | 1.8 |
| 163 | S | AGC | 0 | 0 | 0.1 | 0 | 0 | 0 | 2.5 | 0 | R | CGA | 5.8 |
| 163 | S | AGC | 0 | 0 | 0.1 | 0 | 0 | 0 | 2.5 | 0 | R | CGG | 5.2 |
| 163 | S | AGC | 0 | 0 | 0.1 | 0 | 0 | 0 | 2.5 | 0 | R | AGA | 1.2 |
| 163 | S | AGC | 0 | 0 | 0.1 | 0 | 0 | 0 | 2.5 | 0 | R | AGG | 3.6 |
| 163 | S | AGT | 0 | 0 | 0 | 0 | 0 | 0 | 2.9 | 0 | R | CGT | 1.8 |
| 163 | S | AGT | 0 | 0 | 0 | 0 | 0 | 0 | 2.9 | 0 | R | CGC | 5.2 |
| 163 | S | AGT | 0 | 0 | 0 | 0 | 0 | 0 | 2.9 | 0 | R | CGA | 7.3 |
| 163 | S | AGT | 0 | 0 | 0 | 0 | 0 | 0 | 2.9 | 0 | R | CGG | 44 |
| 163 | S | AGT | 0 | 0 | 0 | 0 | 0 | 0 | 2.9 | 0 | R | AGA | 3 |
| 163 | S | AGT | 0 | 0 | 0 | 0 | 0 | 0 | 2.9 | 0 | R | AGG | 5.3 |
| 163 | A | GCG | 0 | 0 | 0 | 0.1 | 0.4 | 0 | 0 | 1.9 | R | CGT | 54.9 |
| 163 | A | GCG | 0 | 0 | 0 | 0.1 | 0.4 | 0 | 0 | 1.9 | R | CGC | 47.4 |
| 163 | A | GCG | 0 | 0 | 0 | 0.1 | 0.4 | 0 | 0 | 1.9 | R | CGA | 10.6 |
| 163 | A | GCG | 0 | 0 | 0 | 0.1 | 0.4 | 0 | 0 | 1.9 | R | CGG | 9.2 |
| 163 | A | GCG | 0 | 0 | 0 | 0.1 | 0.4 | 0 | 0 | 1.9 | R | AGA | 5.8 |
| 163 | A | GCG | 0 | 0 | 0 | 0.1 | 0.4 | 0 | 0 | 1.9 | R | AGG | 4.4 |
| 163 | M | ATG | 0 | 0 | 0.2 | 0 | 0 | 0 | 0 | 1.2 | R | CGT | 89.7 |
| 163 | M | ATG | 0 | 0 | 0.2 | 0 | 0 | 0 | 0 | 1.2 | R | CGC | 82.2 |
| 163 | M | ATG | 0 | 0 | 0.2 | 0 | 0 | 0 | 0 | 1.2 | R | CGA | 45.4 |
| 163 | M | ATG | 0 | 0 | 0.2 | 0 | 0 | 0 | 0 | 1.2 | R | CGG | 44 |
| 163 | M | ATG | 0 | 0 | 0.2 | 0 | 0 | 0 | 0 | 1.2 | R | AGA | 6.3 |
| 163 | M | ATG | 0 | 0 | 0.2 | 0 | 0 | 0 | 0 | 1.2 | R | AGG | 5.3 |
| 165 | V | GTA | 85 | 89.4 | 92.8 | 80.8 | 4.4 | 88.2 | 65.2 | 9 | I | ATT | 7.4 |
| 165 | V | GTA | 85 | 89.4 | 92.8 | 80.8 | 4.4 | 88.2 | 65.2 | 9 | I | ATC | 9.2 |

| | | | | | | | | | | | | | |
|---|---|---|---|---|---|---|---|---|---|---|---|---|---|
| 165 | V | GTA | 85 | 89.4 | 92.8 | 80.8 | 4.4 | 88.2 | 65.2 | 9 | I | ATA | 1 |
| 165 | V | GTC | 9.5 | 0.9 | 0.5 | 0.7 | 83.1 | 1.2 | 0.8 | 70.8 | I | ATT | 2.9 |
| 165 | V | GTC | 9.5 | 0.9 | 0.5 | 0.7 | 83.1 | 1.2 | 0.8 | 70.8 | I | ATC | 1 |
| 165 | V | GTC | 9.5 | 0.9 | 0.5 | 0.7 | 83.1 | 1.2 | 0.8 | 70.8 | I | ATA | 5 |
| 165 | I | ATA | 2.2 | 3.4 | 3.9 | 17.2 | 0.5 | 6.2 | 29.8 | 1.9 | I | ATT | 7.4 |
| 165 | I | ATA | 2.2 | 3.4 | 3.9 | 17.2 | 0.5 | 6.2 | 29.8 | 1.9 | I | ATC | 1.8 |
| 165 | I | ATA | 2.2 | 3.4 | 3.9 | 17.2 | 0.5 | 6.2 | 29.8 | 1.9 | I | ATA | 0 |
| 165 | V | GTG | 1 | 5.9 | 2.5 | 0.9 | 0.2 | 3.8 | 2.7 | 0.3 | I | ATT | 21.2 |
| 165 | V | GTG | 1 | 5.9 | 2.5 | 0.9 | 0.2 | 3.8 | 2.7 | 0.3 | I | ATC | 3.9 |
| 165 | V | GTG | 1 | 5.9 | 2.5 | 0.9 | 0.2 | 3.8 | 2.7 | 0.3 | I | ATA | 2 |
| 165 | V | GTT | 1 | 0.3 | 0.4 | 0.4 | 10.2 | 0.6 | 0.8 | 14 | I | ATT | 1 |
| 165 | V | GTT | 1 | 0.3 | 0.4 | 0.4 | 10.2 | 0.6 | 0.8 | 14 | I | ATC | 4.4 |
| 165 | V | GTT | 1 | 0.3 | 0.4 | 0.4 | 10.2 | 0.6 | 0.8 | 14 | I | ATA | 6.5 |
| 165 | I | ATC | 1 | 0 | 0 | 0 | 1.3 | 0 | 0.4 | 3.4 | I | ATT | 2.6 |
| 165 | I | ATC | 1 | 0 | 0 | 0 | 1.3 | 0 | 0.4 | 3.4 | I | ATC | 0 |
| 165 | I | ATC | 1 | 0 | 0 | 0 | 1.3 | 0 | 0.4 | 3.4 | I | ATA | 1.2 |
| 166 | R | AGA | 84.3 | 93.4 | 95.1 | 92.8 | 13.7 | 93.2 | 94.4 | 39.1 | S | TCT | 18.6 |
| 166 | R | AGA | 84.3 | 93.4 | 95.1 | 92.8 | 13.7 | 93.2 | 94.4 | 39.1 | S | TCC | 16 |
| 166 | R | AGA | 84.3 | 93.4 | 95.1 | 92.8 | 13.7 | 93.2 | 94.4 | 39.1 | S | TCA | 10.3 |
| 166 | R | AGA | 84.3 | 93.4 | 95.1 | 92.8 | 13.7 | 93.2 | 94.4 | 39.1 | S | TCG | 13.6 |
| 166 | R | AGA | 84.3 | 93.4 | 95.1 | 92.8 | 13.7 | 93.2 | 94.4 | 39.1 | S | AGT | 7.4 |
| 166 | R | AGA | 84.3 | 93.4 | 95.1 | 92.8 | 13.7 | 93.2 | 94.4 | 39.1 | S | AGC | 1.8 |
| 166 | R | AGG | 15.1 | 3.7 | 4.1 | 6.5 | 86.2 | 3.2 | 4.9 | 59.9 | S | TCT | 56 |
| 166 | R | AGG | 15.1 | 3.7 | 4.1 | 6.5 | 86.2 | 3.2 | 4.9 | 59.9 | S | TCC | 48.5 |
| 166 | R | AGG | 15.1 | 3.7 | 4.1 | 6.5 | 86.2 | 3.2 | 4.9 | 59.9 | S | TCA | 11.7 |
| 166 | R | AGG | 15.1 | 3.7 | 4.1 | 6.5 | 86.2 | 3.2 | 4.9 | 59.9 | S | TCG | 10.3 |
| 166 | R | AGG | 15.1 | 3.7 | 4.1 | 6.5 | 86.2 | 3.2 | 4.9 | 59.9 | S | AGT | 13.5 |
| 166 | R | AGG | 15.1 | 3.7 | 4.1 | 6.5 | 86.2 | 3.2 | 4.9 | 59.9 | S | AGC | 5.8 |
| 166 | R | CGT | 0.2 | 2.2 | 0 | 0 | 0 | 1.2 | 0 | 0 | S | TCT | 5.5 |
| 166 | R | CGT | 0.2 | 2.2 | 0 | 0 | 0 | 1.2 | 0 | 0 | S | TCC | 7.5 |
| 166 | R | CGT | 0.2 | 2.2 | 0 | 0 | 0 | 1.2 | 0 | 0 | S | TCA | 9.4 |
| 166 | R | CGT | 0.2 | 2.2 | 0 | 0 | 0 | 1.2 | 0 | 0 | S | TCG | 56.7 |
| 166 | R | CGT | 0.2 | 2.2 | 0 | 0 | 0 | 1.2 | 0 | 0 | S | AGT | 1.2 |
| 166 | R | CGT | 0.2 | 2.2 | 0 | 0 | 0 | 1.2 | 0 | 0 | S | AGC | 4.6 |

| | | | | | | | | | | | | | |
|---|---|---|---|---|---|---|---|---|---|---|---|---|---|
| 166 | R | CGA | 0.1 | 0.4 | 0.2 | 0.5 | 0.1 | 1.2 | 0.4 | 0.3 | S | TCT | 13.8 |
| 166 | R | CGA | 0.1 | 0.4 | 0.2 | 0.5 | 0.1 | 1.2 | 0.4 | 0.3 | S | TCC | 11.2 |
| 166 | R | CGA | 0.1 | 0.4 | 0.2 | 0.5 | 0.1 | 1.2 | 0.4 | 0.3 | S | TCA | 5.5 |
| 166 | R | CGA | 0.1 | 0.4 | 0.2 | 0.5 | 0.1 | 1.2 | 0.4 | 0.3 | S | TCG | 8.8 |
| 166 | R | CGA | 0.1 | 0.4 | 0.2 | 0.5 | 0.1 | 1.2 | 0.4 | 0.3 | S | AGT | 7.6 |
| 166 | R | CGA | 0.1 | 0.4 | 0.2 | 0.5 | 0.1 | 1.2 | 0.4 | 0.3 | S | AGC | 9.4 |
| 166 | K | AAA | 0.2 | 0.3 | 0.1 | 0.1 | 0 | 1.2 | 0.4 | 0 | S | TCT | 23.9 |
| 166 | K | AAA | 0.2 | 0.3 | 0.1 | 0.1 | 0 | 1.2 | 0.4 | 0 | S | TCC | 21.3 |
| 166 | K | AAA | 0.2 | 0.3 | 0.1 | 0.1 | 0 | 1.2 | 0.4 | 0 | S | TCA | 15.6 |
| 166 | K | AAA | 0.2 | 0.3 | 0.1 | 0.1 | 0 | 1.2 | 0.4 | 0 | S | TCG | 18.9 |
| 166 | K | AAA | 0.2 | 0.3 | 0.1 | 0.1 | 0 | 1.2 | 0.4 | 0 | S | AGT | 7.4 |
| 166 | K | AAA | 0.2 | 0.3 | 0.1 | 0.1 | 0 | 1.2 | 0.4 | 0 | S | AGC | 9.2 |
| 170 | E | GAA | 95.4 | 94.7 | 9 | 94.2 | 96.7 | 93.2 | 92.2 | 96.2 | A | GCT | 8.2 |
| 170 | E | GAA | 95.4 | 94.7 | 9 | 94.2 | 96.7 | 93.2 | 92.2 | 96.2 | A | GCC | 10 |
| 170 | E | GAA | 95.4 | 94.7 | 9 | 94.2 | 96.7 | 93.2 | 92.2 | 96.2 | A | GCA | 1.8 |
| 170 | E | GAA | 95.4 | 94.7 | 9 | 94.2 | 96.7 | 93.2 | 92.2 | 96.2 | A | GCG | 4.3 |
| 170 | E | GAG | 4.6 | 5.2 | 91 | 5.7 | 3.3 | 6.8 | 7.8 | 3.8 | A | GCT | 22 |
| 170 | E | GAG | 4.6 | 5.2 | 91 | 5.7 | 3.3 | 6.8 | 7.8 | 3.8 | A | GCC | 4.7 |
| 170 | E | GAG | 4.6 | 5.2 | 91 | 5.7 | 3.3 | 6.8 | 7.8 | 3.8 | A | GCA | 2.8 |
| 170 | E | GAG | 4.6 | 5.2 | 91 | 5.7 | 3.3 | 6.8 | 7.8 | 3.8 | A | GCG | 1.8 |
| 201 | I | ATA | 94.9 | 40.3 | 93.7 | 98.7 | 98.2 | 95.9 | 95.9 | 96.9 | I | ATT | 7.4 |
| 201 | I | ATA | 94.9 | 40.3 | 93.7 | 98.7 | 98.2 | 95.9 | 95.9 | 96.9 | I | ATC | 1.8 |
| 201 | I | ATA | 94.9 | 40.3 | 93.7 | 98.7 | 98.2 | 95.9 | 95.9 | 96.9 | I | ATA | 0 |
| 201 | V | GTA | 4.9 | 56.6 | 6.2 | 1.1 | 1.6 | 1.8 | 3.7 | 3.1 | I | ATT | 7.4 |
| 201 | V | GTA | 4.9 | 56.6 | 6.2 | 1.1 | 1.6 | 1.8 | 3.7 | 3.1 | I | ATC | 9.2 |
| 201 | V | GTA | 4.9 | 56.6 | 6.2 | 1.1 | 1.6 | 1.8 | 3.7 | 3.1 | I | ATA | 1 |
| 201 | V | GTG | 0 | 2.3 | 0.1 | 0 | 0 | 0 | 0 | 0 | I | ATT | 21.2 |
| 201 | V | GTG | 0 | 2.3 | 0.1 | 0 | 0 | 0 | 0 | 0 | I | ATC | 3.9 |
| 201 | V | GTG | 0 | 2.3 | 0.1 | 0 | 0 | 0 | 0 | 0 | I | ATA | 2 |
| 201 | I | ATT | 0.2 | 0.7 | 0 | 0 | 0.2 | 1.8 | 0 | 0 | I | ATT | 0 |
| 201 | I | ATT | 0.2 | 0.7 | 0 | 0 | 0.2 | 1.8 | 0 | 0 | I | ATC | 1.6 |
| 201 | I | ATT | 0.2 | 0.7 | 0 | 0 | 0.2 | 1.8 | 0 | 0 | I | ATA | 3 |
| 203 | I | ATA | 93.8 | 90.1 | 97.2 | 94.8 | 97.4 | 79.4 | 98.8 | 96.2 | M | ATG | 1 |
| 203 | M | ATG | 5.5 | 6.4 | 2.2 | 4.2 | 1.7 | 15.3 | 0.8 | 3.4 | M | ATG | 0 |

| | | | | | | | | | | | | | |
|---|---|---|---|---|---|---|---|---|---|---|---|---|---|
| 203 | I | ATT | 0.2 | 2.4 | 0 | 0.2 | 0.1 | 1.8 | 0 | 0 | M | ATG | 5.3 |
| 203 | I | ATC | 0.5 | 0.6 | 0.5 | 0.8 | 0.8 | 3.5 | 0.4 | 0.3 | M | ATG | 3.6 |
| 203 | I | ATA | 93.8 | 90.1 | 97.2 | 94.8 | 97.4 | 79.4 | 98.8 | 96.2 | R | CGT | 52.3 |
| 203 | I | ATA | 93.8 | 90.1 | 97.2 | 94.8 | 97.4 | 79.4 | 98.8 | 96.2 | R | CGC | 49.7 |
| 203 | I | ATA | 93.8 | 90.1 | 97.2 | 94.8 | 97.4 | 79.4 | 98.8 | 96.2 | R | CGA | 44 |
| 203 | I | ATA | 93.8 | 90.1 | 97.2 | 94.8 | 97.4 | 79.4 | 98.8 | 96.2 | R | CGG | 47.3 |
| 203 | I | ATA | 93.8 | 90.1 | 97.2 | 94.8 | 97.4 | 79.4 | 98.8 | 96.2 | R | AGA | 5.3 |
| 203 | I | ATA | 93.8 | 90.1 | 97.2 | 94.8 | 97.4 | 79.4 | 98.8 | 96.2 | R | AGG | 7.8 |
| 203 | M | ATG | 5.5 | 6.4 | 2.2 | 4.2 | 1.7 | 15.3 | 0.8 | 3.4 | R | CGT | 89.7 |
| 203 | M | ATG | 5.5 | 6.4 | 2.2 | 4.2 | 1.7 | 15.3 | 0.8 | 3.4 | R | CGC | 82.2 |
| 203 | M | ATG | 5.5 | 6.4 | 2.2 | 4.2 | 1.7 | 15.3 | 0.8 | 3.4 | R | CGA | 45.4 |
| 203 | M | ATG | 5.5 | 6.4 | 2.2 | 4.2 | 1.7 | 15.3 | 0.8 | 3.4 | R | CGG | 44 |
| 203 | M | ATG | 5.5 | 6.4 | 2.2 | 4.2 | 1.7 | 15.3 | 0.8 | 3.4 | R | AGA | 6.3 |
| 203 | M | ATG | 5.5 | 6.4 | 2.2 | 4.2 | 1.7 | 15.3 | 0.8 | 3.4 | R | AGG | 5.3 |
| 203 | I | ATT | 0.2 | 2.4 | 0 | 0.2 | 0.1 | 1.8 | 0 | 0 | R | CGT | 44 |
| 203 | I | ATT | 0.2 | 2.4 | 0 | 0.2 | 0.1 | 1.8 | 0 | 0 | R | CGC | 46 |
| 203 | I | ATT | 0.2 | 2.4 | 0 | 0.2 | 0.1 | 1.8 | 0 | 0 | R | CGA | 47.9 |
| 203 | I | ATT | 0.2 | 2.4 | 0 | 0.2 | 0.1 | 1.8 | 0 | 0 | R | CGG | 95.2 |
| 203 | I | ATT | 0.2 | 2.4 | 0 | 0.2 | 0.1 | 1.8 | 0 | 0 | R | AGA | 10.8 |
| 203 | I | ATT | 0.2 | 2.4 | 0 | 0.2 | 0.1 | 1.8 | 0 | 0 | R | AGG | 47.5 |
| 203 | I | ATC | 0.5 | 0.6 | 0.5 | 0.8 | 0.8 | 3.5 | 0.4 | 0.3 | R | CGT | 45 |
| 203 | I | ATC | 0.5 | 0.6 | 0.5 | 0.8 | 0.8 | 3.5 | 0.4 | 0.3 | R | CGC | 44 |
| 203 | I | ATC | 0.5 | 0.6 | 0.5 | 0.8 | 0.8 | 3.5 | 0.4 | 0.3 | R | CGA | 45.3 |
| 203 | I | ATC | 0.5 | 0.6 | 0.5 | 0.8 | 0.8 | 3.5 | 0.4 | 0.3 | R | CGG | 65 |
| 203 | I | ATC | 0.5 | 0.6 | 0.5 | 0.8 | 0.8 | 3.5 | 0.4 | 0.3 | R | AGA | 9.3 |
| 203 | I | ATC | 0.5 | 0.6 | 0.5 | 0.8 | 0.8 | 3.5 | 0.4 | 0.3 | R | AGG | 8.7 |
| 206 | T | ACA | 93.5 | 83.7 | 90.7 | 95.3 | 4.7 | 84.7 | 85 | 6.2 | S | TCT | 13.8 |
| 206 | T | ACA | 93.5 | 83.7 | 90.7 | 95.3 | 4.7 | 84.7 | 85 | 6.2 | S | TCC | 15.6 |
| 206 | T | ACA | 93.5 | 83.7 | 90.7 | 95.3 | 4.7 | 84.7 | 85 | 6.2 | S | TCA | 7.4 |
| 206 | T | ACA | 93.5 | 83.7 | 90.7 | 95.3 | 4.7 | 84.7 | 85 | 6.2 | S | TCG | 9.9 |
| 206 | T | ACA | 93.5 | 83.7 | 90.7 | 95.3 | 4.7 | 84.7 | 85 | 6.2 | S | AGT | 10 |
| 206 | T | ACA | 93.5 | 83.7 | 90.7 | 95.3 | 4.7 | 84.7 | 85 | 6.2 | S | AGC | 11.8 |
| 206 | S | TCA | 5.9 | 13.7 | 8.9 | 4.5 | 95 | 13.5 | 15 | 93.8 | S | TCT | 7.4 |
| 206 | S | TCA | 5.9 | 13.7 | 8.9 | 4.5 | 95 | 13.5 | 15 | 93.8 | S | TCC | 1.8 |

| | | | | | | | | | | | | |
|---|---|---|---|---|---|---|---|---|---|---|---|---|
| 206 | S | TCA | 5.9 | 13.7 | 8.9 | 4.5 | 95 | 13.5 | 15 | 93.8 | S | TCA | 0 |
| 206 | S | TCA | 5.9 | 13.7 | 8.9 | 4.5 | 95 | 13.5 | 15 | 93.8 | S | TCG | 1 |
| 206 | S | TCA | 5.9 | 13.7 | 8.9 | 4.5 | 95 | 13.5 | 15 | 93.8 | S | AGT | 14.7 |
| 206 | S | TCA | 5.9 | 13.7 | 8.9 | 4.5 | 95 | 13.5 | 15 | 93.8 | S | AGC | 12.1 |
| 206 | T | ACC | 0.2 | 2 | 0 | 0.1 | 0 | 0.6 | 0 | 0 | S | TCT | 9.3 |
| 206 | T | ACC | 0.2 | 2 | 0 | 0.1 | 0 | 0.6 | 0 | 0 | S | TCC | 7.4 |
| 206 | T | ACC | 0.2 | 2 | 0 | 0.1 | 0 | 0.6 | 0 | 0 | S | TCA | 11.4 |
| 206 | T | ACC | 0.2 | 2 | 0 | 0.1 | 0 | 0.6 | 0 | 0 | S | TCG | 10.8 |
| 206 | T | ACC | 0.2 | 2 | 0 | 0.1 | 0 | 0.6 | 0 | 0 | S | AGT | 5.5 |
| 206 | T | ACC | 0.2 | 2 | 0 | 0.1 | 0 | 0.6 | 0 | 0 | S | AGC | 3.6 |
| 206 | S | AGC | 0 | 0.4 | 0 | 0 | 0 | 1.2 | 0 | 0 | S | TCT | 11.3 |
| 206 | S | AGC | 0 | 0.4 | 0 | 0 | 0 | 1.2 | 0 | 0 | S | TCC | 10.3 |
| 206 | S | AGC | 0 | 0.4 | 0 | 0 | 0 | 1.2 | 0 | 0 | S | TCA | 11.6 |
| 206 | S | AGC | 0 | 0.4 | 0 | 0 | 0 | 1.2 | 0 | 0 | S | TCG | 31.3 |
| 206 | S | AGC | 0 | 0.4 | 0 | 0 | 0 | 1.2 | 0 | 0 | S | AGT | 2.6 |
| 206 | S | AGC | 0 | 0.4 | 0 | 0 | 0 | 1.2 | 0 | 0 | S | AGC | 0 |
| 230 | S | AGC | 98.5 | 89.2 | 98.4 | 99.2 | 99.6 | 97.1 | 98.6 | 98.1 | G | GGT | 2.9 |
| 230 | S | AGC | 98.5 | 89.2 | 98.4 | 99.2 | 99.6 | 97.1 | 98.6 | 98.1 | G | GGC | 1 |
| 230 | S | AGC | 98.5 | 89.2 | 98.4 | 99.2 | 99.6 | 97.1 | 98.6 | 98.1 | G | GGA | 5 |
| 230 | S | AGC | 98.5 | 89.2 | 98.4 | 99.2 | 99.6 | 97.1 | 98.6 | 98.1 | G | GGG | 4.4 |
| 230 | N | AAC | 1.5 | 10.4 | 1.5 | 0.7 | 0.3 | 2.9 | 1.2 | 1.9 | G | GGT | 4.5 |
| 230 | N | AAC | 1.5 | 10.4 | 1.5 | 0.7 | 0.3 | 2.9 | 1.2 | 1.9 | G | GGC | 3.5 |
| 230 | N | AAC | 1.5 | 10.4 | 1.5 | 0.7 | 0.3 | 2.9 | 1.2 | 1.9 | G | GGA | 4.8 |
| 230 | N | AAC | 1.5 | 10.4 | 1.5 | 0.7 | 0.3 | 2.9 | 1.2 | 1.9 | G | GGG | 24.5 |
| 230 | S | AGC | 98.5 | 89.2 | 98.4 | 99.2 | 99.6 | 97.1 | 98.6 | 98.1 | N | AAT | 2.9 |
| 230 | S | AGC | 98.5 | 89.2 | 98.4 | 99.2 | 99.6 | 97.1 | 98.6 | 98.1 | N | AAC | 1 |
| 230 | N | AAC | 1.5 | 10.4 | 1.5 | 0.7 | 0.3 | 2.9 | 1.2 | 1.9 | N | AAT | 2.6 |
| 230 | N | AAC | 1.5 | 10.4 | 1.5 | 0.7 | 0.3 | 2.9 | 1.2 | 1.9 | N | AAC | 0 |
| 230 | S | AGC | 98.5 | 89.2 | 98.4 | 99.2 | 99.6 | 97.1 | 98.6 | 98.1 | R | CGT | 3.7 |
| 230 | S | AGC | 98.5 | 89.2 | 98.4 | 99.2 | 99.6 | 97.1 | 98.6 | 98.1 | R | CGC | 1.8 |
| 230 | S | AGC | 98.5 | 89.2 | 98.4 | 99.2 | 99.6 | 97.1 | 98.6 | 98.1 | R | CGA | 5.8 |
| 230 | S | AGC | 98.5 | 89.2 | 98.4 | 99.2 | 99.6 | 97.1 | 98.6 | 98.1 | R | CGG | 5.2 |
| 230 | S | AGC | 98.5 | 89.2 | 98.4 | 99.2 | 99.6 | 97.1 | 98.6 | 98.1 | R | AGA | 1.2 |
| 230 | S | AGC | 98.5 | 89.2 | 98.4 | 99.2 | 99.6 | 97.1 | 98.6 | 98.1 | R | AGG | 3.6 |

| | | | | | | | | | | | | | |
|---|---|---|---|---|---|---|---|---|---|---|---|---|---|
| 230 | N | AAC | 1.5 | 10.4 | 1.5 | 0.7 | 0.3 | 2.9 | 1.2 | 1.9 | R | CGT | 5.3 |
| 230 | N | AAC | 1.5 | 10.4 | 1.5 | 0.7 | 0.3 | 2.9 | 1.2 | 1.9 | R | CGC | 4.3 |
| 230 | N | AAC | 1.5 | 10.4 | 1.5 | 0.7 | 0.3 | 2.9 | 1.2 | 1.9 | R | CGA | 5.6 |
| 230 | N | AAC | 1.5 | 10.4 | 1.5 | 0.7 | 0.3 | 2.9 | 1.2 | 1.9 | R | CGG | 25.3 |
| 230 | N | AAC | 1.5 | 10.4 | 1.5 | 0.7 | 0.3 | 2.9 | 1.2 | 1.9 | R | AGA | 5 |
| 230 | N | AAC | 1.5 | 10.4 | 1.5 | 0.7 | 0.3 | 2.9 | 1.2 | 1.9 | R | AGG | 4.4 |
| 232 | D | GAT | 68.9 | 86.9 | 12.4 | 13.4 | 32.8 | 90.8 | 7.8 | 15.3 | N | AAT | 1 |
| 232 | D | GAT | 68.9 | 86.9 | 12.4 | 13.4 | 32.8 | 90.8 | 7.8 | 15.3 | N | AAC | 4.4 |
| 232 | D | GAC | 28.5 | 8.8 | 87.3 | 86.2 | 66 | 9.2 | 90.2 | 82.2 | N | AAT | 2.9 |
| 232 | D | GAC | 28.5 | 8.8 | 87.3 | 86.2 | 66 | 9.2 | 90.2 | 82.2 | N | AAC | 1 |
| 232 | E | GAG | 0.7 | 3 | 0 | 0.1 | 0.2 | 0 | 0 | 0 | N | AAT | 21.2 |
| 232 | E | GAG | 0.7 | 3 | 0 | 0.1 | 0.2 | 0 | 0 | 0 | N | AAC | 3.9 |
| 232 | E | GAA | 0.1 | 0.9 | 0.2 | 0.3 | 1 | 0 | 2 | 1.9 | N | AAT | 7.4 |
| 232 | E | GAA | 0.1 | 0.9 | 0.2 | 0.3 | 1 | 0 | 2 | 1.9 | N | AAC | 9.2 |
| 232 | N | AAT | 1.5 | 0.2 | 0 | 0 | 0 | 0 | 0 | 0.6 | N | AAT | 0 |
| 232 | N | AAT | 1.5 | 0.2 | 0 | 0 | 0 | 0 | 0 | 0.6 | N | AAC | 1.6 |
| 263 | R | AGA | 94.1 | 91.9 | 9.7 | 97 | 94.8 | 90.6 | 91.4 | 94.3 | K | AAA | 1 |
| 263 | R | AGA | 94.1 | 91.9 | 9.7 | 97 | 94.8 | 90.6 | 91.4 | 94.3 | K | AAG | 3.5 |
| 263 | R | AGG | 5.2 | 5.4 | 90.1 | 3 | 4.4 | 7.6 | 8.6 | 5.7 | K | AAA | 2 |
| 263 | R | AGG | 5.2 | 5.4 | 90.1 | 3 | 4.4 | 7.6 | 8.6 | 5.7 | K | AAG | 1 |
| 263 | R | CGT | 0.2 | 2.4 | 0 | 0 | 0 | 1.8 | 0 | 0 | K | AAA | 6.1 |
| 263 | R | CGT | 0.2 | 2.4 | 0 | 0 | 0 | 1.8 | 0 | 0 | K | AAG | 53.4 |

### 6 Codon distributions and cost scores

The population genetic barrier for a resistance-associated mutation and a HIV-1 subtype is a value that results from a complex interplay of different factors, including variability at the codon level and the evolutionary score associated with each triplet. While the main manuscript report differences in genetic barrier, we here try to disentangle the different contributions, supporting other figures on triplet entropy and the prevalence of resistance-associated mutations.

For the HIV-1 drug class of integrase inhibitors (INSTIs), and for each resistance-associated mutation, a heatmap is shown to illustrate the extent of nucleotide triplet diversity and predominance across HIV-1 subtypes, and the associated score based on the transformed sum of all possible cumulative costs, as explained in the manuscript of Theys et al. The x-axis reports for each triplet present at the position the following information:
- the amino acid which corresponds with the wild-type triplet
- the wild-type triplet
- the associated score binned in windows of 0.1
- the maximum number of subtypes sharing the predominant triplet

Figure 7: Codon distributions for each resistance-associated mutation

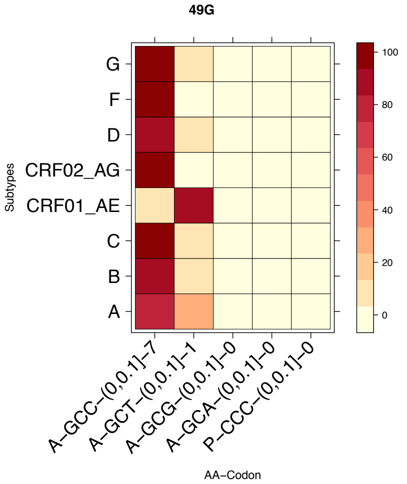
49G
Subtypes
G
F
D
CRF02_AG
CRF01_AE
C
B
A
A–GCC–(0,0.1]–7
A–GCT–(0,0.1]–1
A–GCG–(0,0.1]–0
A–GCA–(0,0.1]–0
P–CCC–(0,0.1]–0
AA–Codon
100
80
60
40
20
0

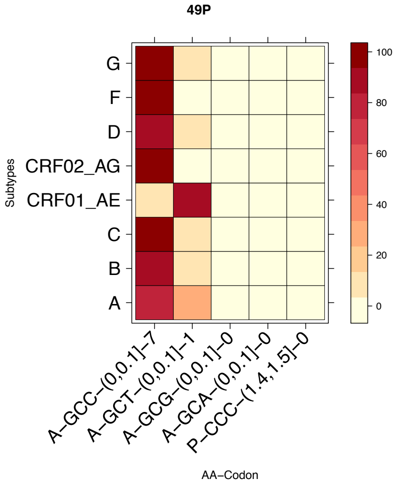
49P
Subtypes
G
F
D
CRF02_AG
CRF01_AE
C
B
A
A–GCC–(0,0.1]–7
A–GCT–(0,0.1]–1
A–GCG–(0,0.1]–0
A–GCA–(0,0.1]–0
P–CCC–(1.4,1.5]–0
AA–Codon
100
80
60
40
20
0

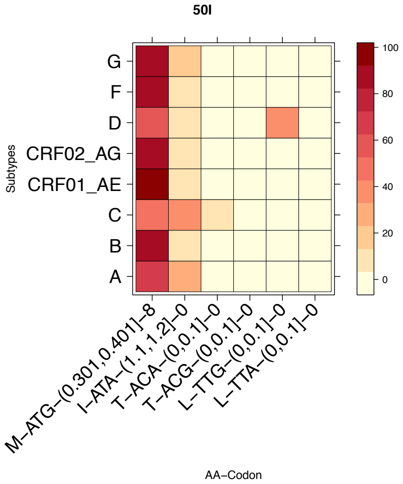
50I
Subtypes
G
F
D
CRF02_AG
CRF01_AE
C
B
A
M–ATG–(0.301,0.401]–8
I–ATA–(1.1,1.2]–0
T–ACA–(0,0.1]–0
T–ACG–(0,0.1]–0
L–TTG–(0,0.1]–0
L–TTA–(0,0.1]–0
AA–Codon
100
80
60
40
20
0

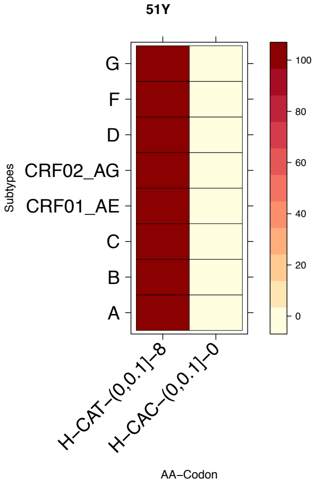
51Y
Subtypes
G
F
D
CRF02_AG
CRF01_AE
C
B
A
H–CAT–(0,0.1]–8
H–CAC–(0,0.1]–0
AA–Codon
100
80
60
40
20
0

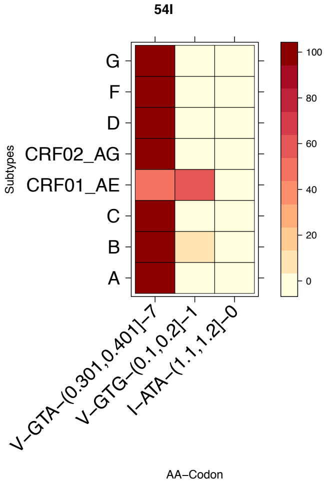
54I
Subtypes
G
F
D
CRF02_AG
CRF01_AE
C
B
A
V–GTA–(0.301,0.401]–7
V–GTG–(0.1,0.2]–1
I–ATA–(1.1,1.2]–0
AA–Codon
100
80
60
40
20
0

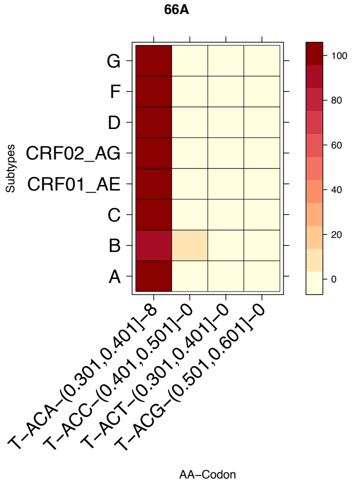
66A
Subtypes
G
F
D
CRF02_AG
CRF01_AE
C
B
A
T–ACA–(0.301,0.401]–8
T–ACC–(0.401,0.501]–0
T–ACT–(0.301,0.401]–0
T–ACG–(0.501,0.601]–0
AA–Codon
100
80
60
40
20
0

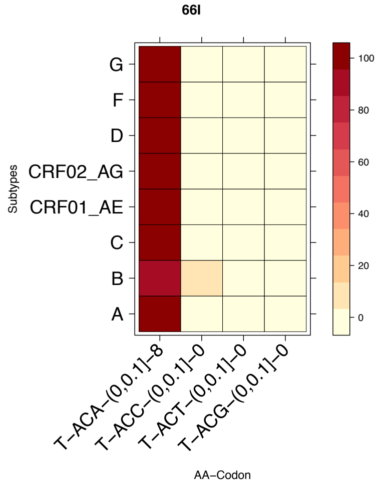
66I
Subtypes
G
F
D
CRF02_AG
CRF01_AE
C
B
A
T–ACA–(0,0.1]–8
T–ACC–(0,0.1]–0
T–ACT–(0,0.1]–0
T–ACG–(0,0.1]–0
AA–Codon
100
80
60
40
20
0

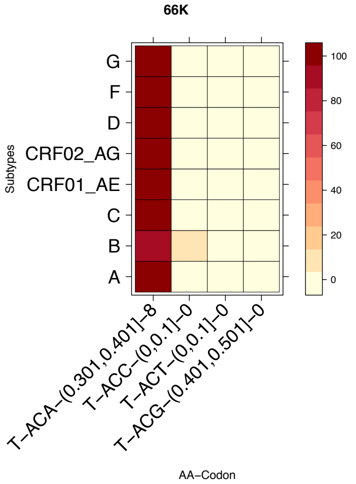
66K
Subtypes
G
F
D
CRF02_AG
CRF01_AE
C
B
A
T–ACA–(0.301,0.401]–8
T–ACC–(0,0.1]–0
T–ACT–(0,0.1]–0
T–ACG–(0.401,0.501]–0
AA–Codon
100
80
60
40
20
0

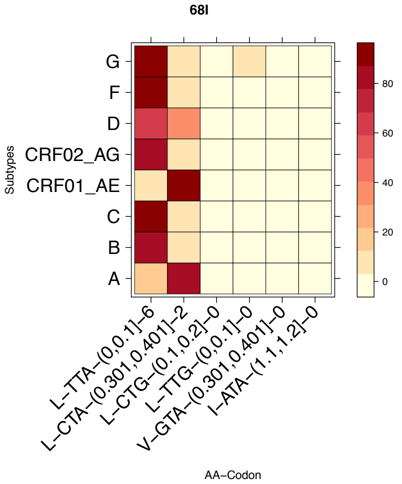
68I
Subtypes
G
F
D
CRF02_AG
CRF01_AE
C
B
A
L–TTA–(0,0.1]–6
L–CTA–(0.301,0.401]–2
L–CTG–(0.1,0.2]–0
L–TTG–(0,0.1]–0
V–GTA–(0.301,0.401]–0
I–ATA–(1.1,1.2]–0
AA–Codon
80
60
40
20
0

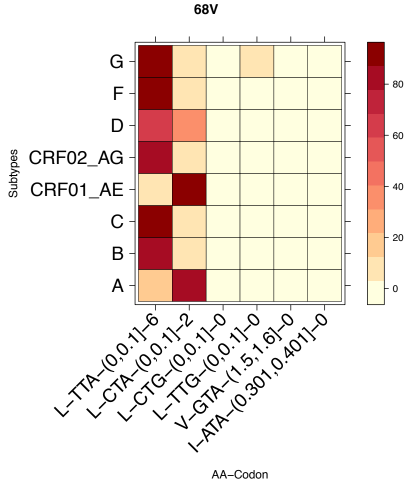
68V
Subtypes
G
F
D
CRF02_AG
CRF01_AE
C
B
A
L–TTA–(0,0.1]–6
L–CTA–(0,0.1]–2
L–CTG–(0,0.1]–0
L–TTG–(0,0.1]–0
V–GTA–(1.5,1.6]–0
I–ATA–(0.301,0.401]–0
AA–Codon
80
60
40
20
0

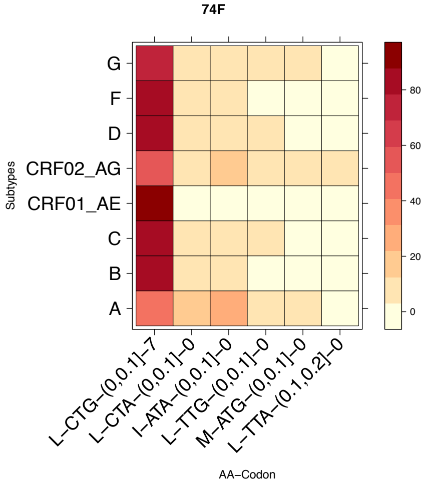
74F
Subtypes
G
F
D
CRF02_AG
CRF01_AE
C
B
A
L–CTG–(0,0.1]–7
L–CTA–(0,0.1]–0
I–ATA–(0,0.1]–0
L–TTG–(0,0.1]–0
M–ATG–(0,0.1]–0
L–TTA–(0.1,0.2]–0
AA–Codon
80
60
40
20
0

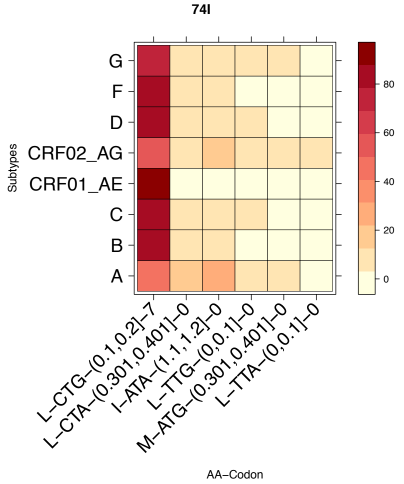
74I
Subtypes
G
F
D
CRF02_AG
CRF01_AE
C
B
A
L–CTG–(0.1,0.2]–7
L–CTA–(0.301,0.401]–0
I–ATA–(1.1,1.2]–0
L–TTG–(0,0.1]–0
M–ATG–(0.301,0.401]–0
L–TTA–(0,0.1]–0
AA–Codon
80
60
40
20
0

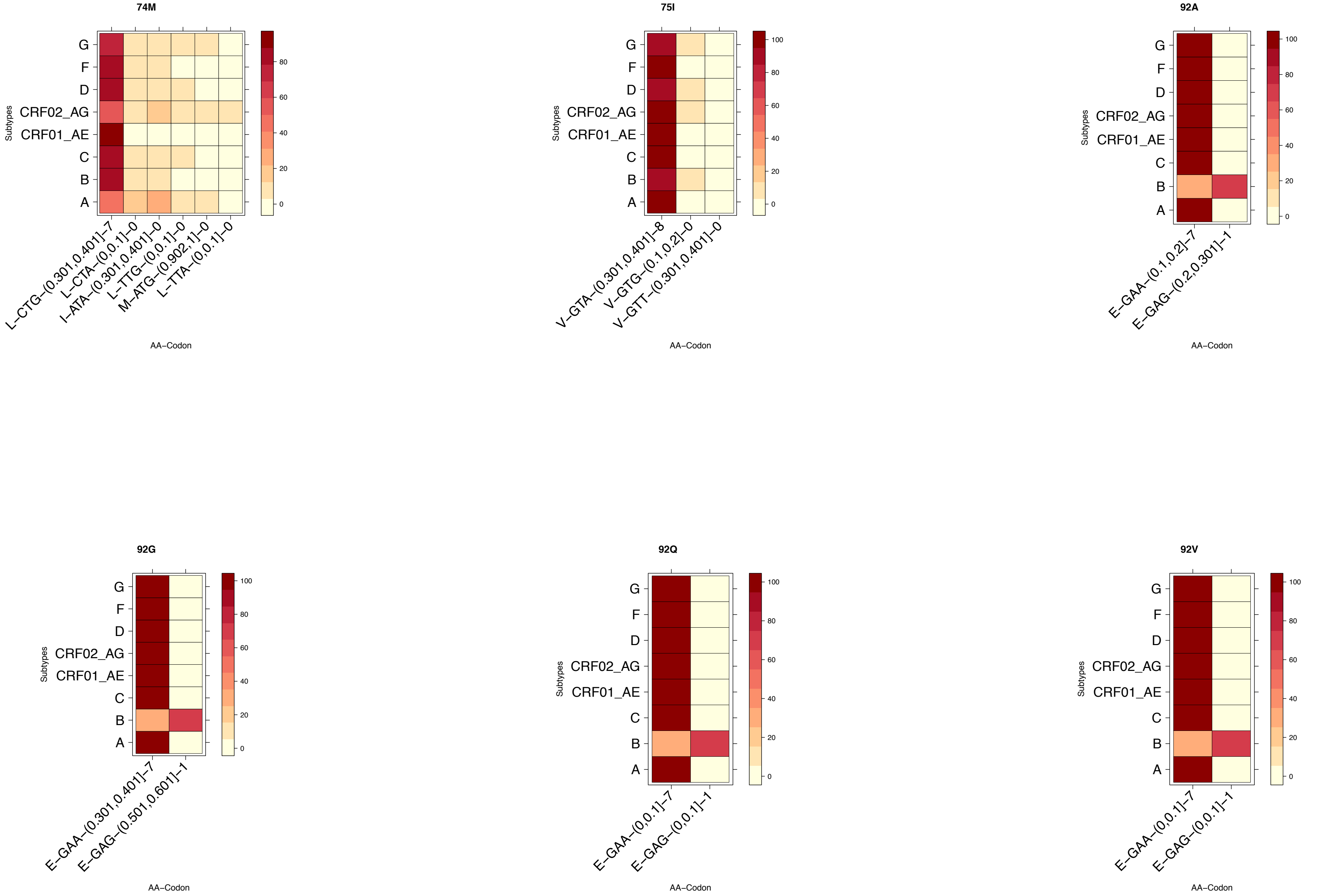
74M
75I
92A
92G
92Q
92V
Subtypes
G
F
D
CRF02_AG
CRF01_AE
C
B
A
AA−Codon
L−CTG−(0.301,0.401]−7
L−CTA−(0,0.1]−0
I−ATA−(0.301,0.401]−0
L−TTG−(0,0.1]−0
M−ATG−(0.902,1]−0
L−TTA−(0,0.1]−0
V−GTA−(0.301,0.401]−8
V−GTG−(0.1,0.2]−0
V−GTT−(0.301,0.401]−0
E−GAA−(0.1,0.2]−7
E−GAG−(0.2,0.301]−1
E−GAA−(0.301,0.401]−7
E−GAG−(0.501,0.601]−1
E−GAA−(0,0.1]−7
E−GAG−(0,0.1]−1
0
20
40
60
80
100

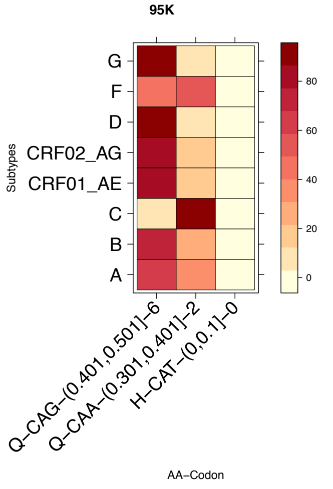
95K
Subtypes
G
F
D
CRF02_AG
CRF01_AE
C
B
A
Q–CAG–(0.401,0.501]–6
Q–CAA–(0.301,0.401]–2
H–CAT–(0,0.1]–0
AA–Codon
0
20
40
60
80

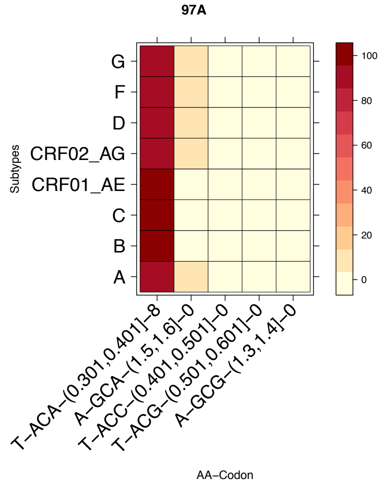
97A
Subtypes
G
F
D
CRF02_AG
CRF01_AE
C
B
A
T–ACA–(0.301,0.401]–8
A–GCA–(1.5,1.6]–0
T–ACC–(0.401,0.501]–0
T–ACG–(0.501,0.601]–0
A–GCG–(1.3,1.4]–0
AA–Codon
0
20
40
60
80
100

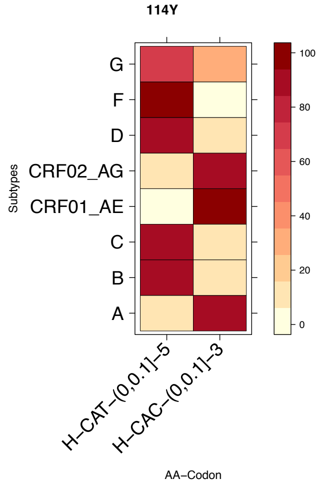
114Y
Subtypes
G
F
D
CRF02_AG
CRF01_AE
C
B
A
H–CAT–(0,0.1]–5
H–CAC–(0,0.1]–3
AA–Codon
0
20
40
60
80
100

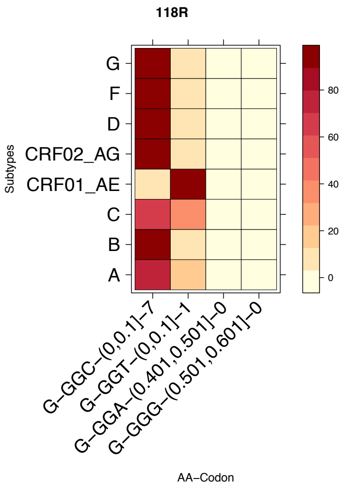
118R
Subtypes
G
F
D
CRF02_AG
CRF01_AE
C
B
A
G–GGC–(0,0.1]–7
G–GGT–(0,0.1]–1
G–GGA–(0.401,0.501]–0
G–GGG–(0.501,0.601]–0
AA–Codon
0
20
40
60
80

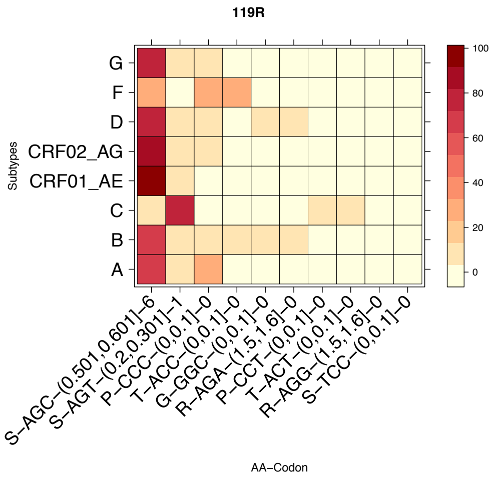
119R
Subtypes
G
F
D
CRF02_AG
CRF01_AE
C
B
A
S–AGC–(0.501,0.601]–6
S–AGT–(0.2,0.301]–1
P–CCC–(0,0.1]–0
T–ACC–(0,0.1]–0
G–GGC–(0,0.1]–0
R–AGA–(1.5,1.6]–0
P–CCT–(0,0.1]–0
T–ACT–(0,0.1]–0
R–AGG–(1.5,1.6]–0
S–TCC–(0,0.1]–0
AA–Codon
0
20
40
60
80
100

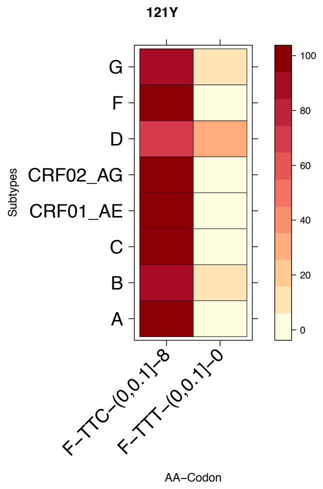
121Y
Subtypes
G
F
D
CRF02_AG
CRF01_AE
C
B
A
F–TTC–(0,0.1]–8
F–TTT–(0,0.1]–0
AA–Codon
0
20
40
60
80
100

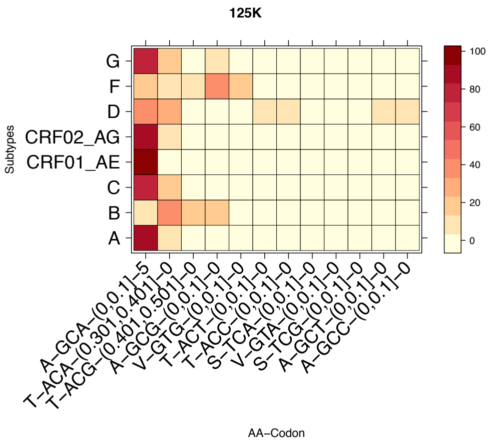
125K
G
F
D
CRF02_AG
CRF01_AE
C
B
A
Subtypes
A–GCA–(0,0.1]–5
T–ACA–(0.301,0.401]–0
T–ACG–(0.401,0.501]–0
A–GCG–(0,0.1]–0
V–GTG–(0,0.1]–0
T–ACT–(0,0.1]–0
T–ACC–(0,0.1]–0
S–TCA–(0,0.1]–0
V–GTA–(0,0.1]–0
S–TCG–(0,0.1]–0
A–GCT–(0,0.1]–0
A–GCC–(0,0.1]–0
AA–Codon
100
80
60
40
20
0

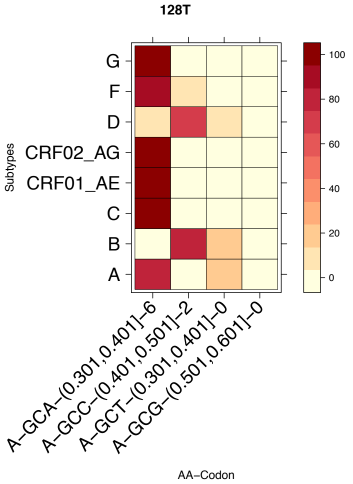
128T
G
F
D
CRF02_AG
CRF01_AE
C
B
A
Subtypes
A–GCA–(0.301,0.401]–6
A–GCC–(0.401,0.501]–2
A–GCT–(0.301,0.401]–0
A–GCG–(0.501,0.601]–0
AA–Codon
100
80
60
40
20
0

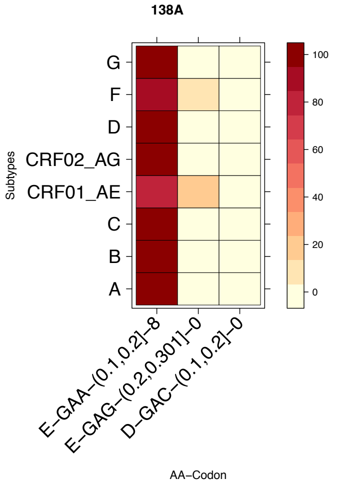
138A
G
F
D
CRF02_AG
CRF01_AE
C
B
A
Subtypes
E–GAA–(0.1,0.2]–8
E–GAG–(0.2,0.301]–0
D–GAC–(0.1,0.2]–0
AA–Codon
100
80
60
40
20
0

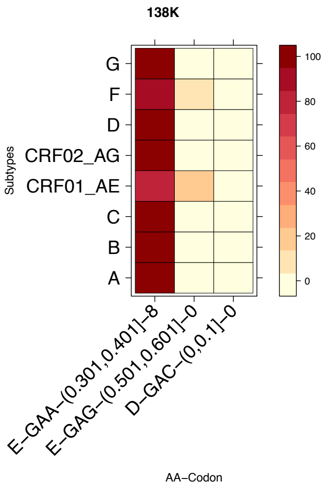
138K
G
F
D
CRF02_AG
CRF01_AE
C
B
A
Subtypes
E–GAA–(0.301,0.401]–8
E–GAG–(0.501,0.601]–0
D–GAC–(0,0.1]–0
AA–Codon
100
80
60
40
20
0

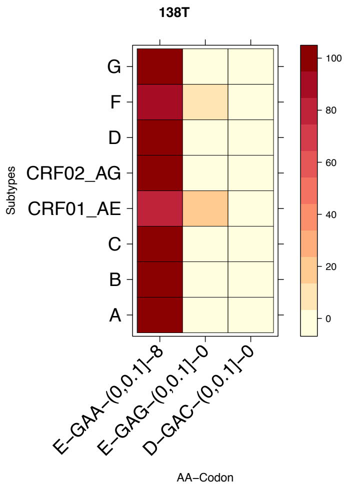
138T
G
F
D
CRF02_AG
CRF01_AE
C
B
A
Subtypes
E–GAA–(0,0.1]–8
E–GAG–(0,0.1]–0
D–GAC–(0,0.1]–0
AA–Codon
100
80
60
40
20
0

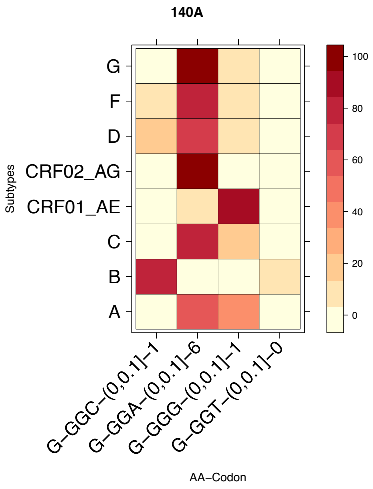
140A
G
F
D
CRF02_AG
CRF01_AE
C
B
A
Subtypes
G–GGC–(0,0.1]–1
G–GGA–(0,0.1]–6
G–GGG–(0,0.1]–1
G–GGT–(0,0.1]–0
AA–Codon
100
80
60
40
20
0

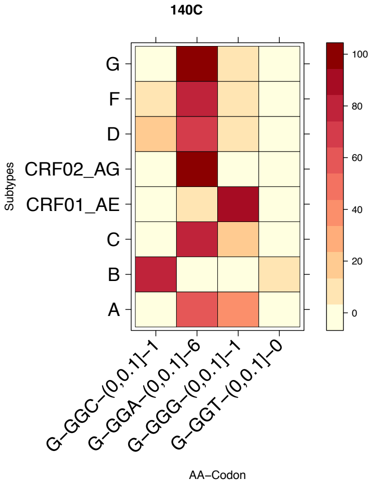
140C
Subtypes
G
F
D
CRF02_AG
CRF01_AE
C
B
A
G–GGC–(0,0.1]–1
G–GGA–(0,0.1]–6
G–GGG–(0,0.1]–1
G–GGT–(0,0.1]–0
AA–Codon
100
80
60
40
20
0

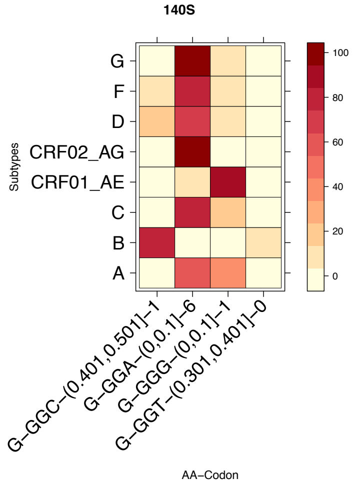
140S
Subtypes
G
F
D
CRF02_AG
CRF01_AE
C
B
A
G–GGC–(0.401,0.501]–1
G–GGA–(0,0.1]–6
G–GGG–(0,0.1]–1
G–GGT–(0.301,0.401]–0
AA–Codon
100
80
60
40
20
0

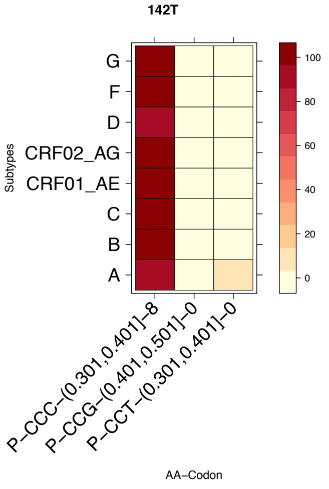
142T
Subtypes
G
F
D
CRF02_AG
CRF01_AE
C
B
A
P–CCC–(0.301,0.401]–8
P–CCG–(0.401,0.501]–0
P–CCT–(0.301,0.401]–0
AA–Codon
100
80
60
40
20
0

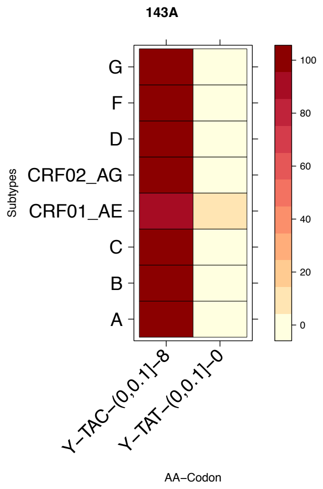
143A
Subtypes
G
F
D
CRF02_AG
CRF01_AE
C
B
A
Y–TAC–(0,0.1]–8
Y–TAT–(0,0.1]–0
AA–Codon
100
80
60
40
20
0

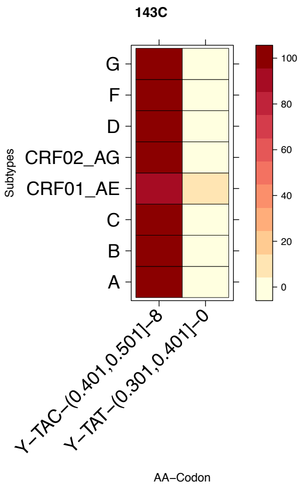
143C
Subtypes
G
F
D
CRF02_AG
CRF01_AE
C
B
A
Y–TAC–(0.401,0.501]–8
Y–TAT–(0.301,0.401]–0
AA–Codon
100
80
60
40
20
0

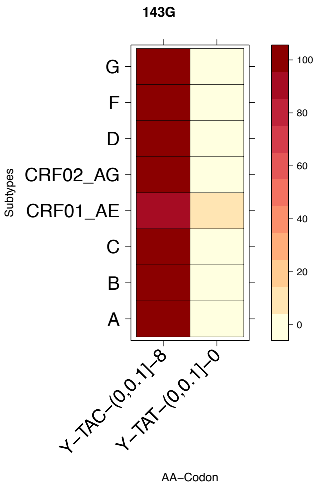
143G
Subtypes
G
F
D
CRF02_AG
CRF01_AE
C
B
A
Y–TAC–(0,0.1]–8
Y–TAT–(0,0.1]–0
AA–Codon
100
80
60
40
20
0

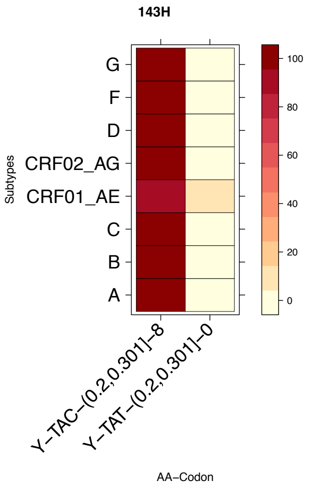
143H
G
F
D
CRF02_AG
CRF01_AE
C
B
A
Subtypes
100
80
60
40
20
0
Y–TAC–(0.2,0.301]–8
Y–TAT–(0.2,0.301]–0
AA–Codon

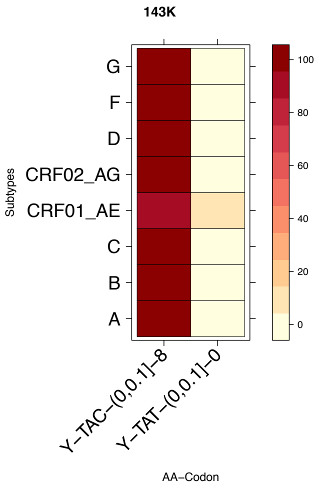
143K
G
F
D
CRF02_AG
CRF01_AE
C
B
A
Subtypes
100
80
60
40
20
0
Y–TAC–(0,0.1]–8
Y–TAT–(0,0.1]–0
AA–Codon

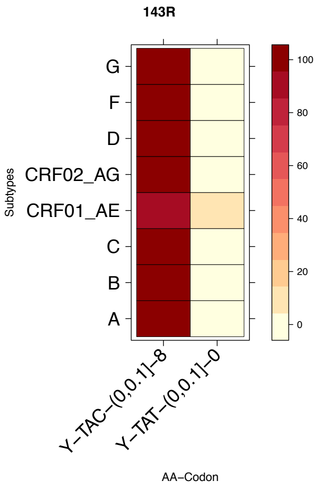
143R
G
F
D
CRF02_AG
CRF01_AE
C
B
A
Subtypes
100
80
60
40
20
0
Y–TAC–(0,0.1]–8
Y–TAT–(0,0.1]–0
AA–Codon

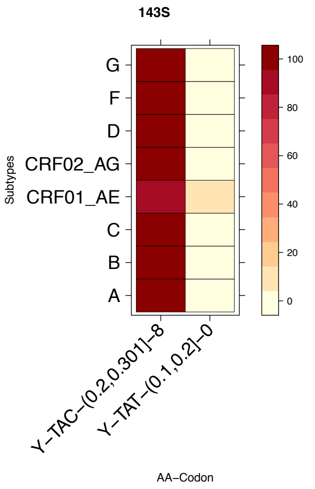
143S
G
F
D
CRF02_AG
CRF01_AE
C
B
A
Subtypes
100
80
60
40
20
0
Y–TAC–(0.2,0.301]–8
Y–TAT–(0.1,0.2]–0
AA–Codon

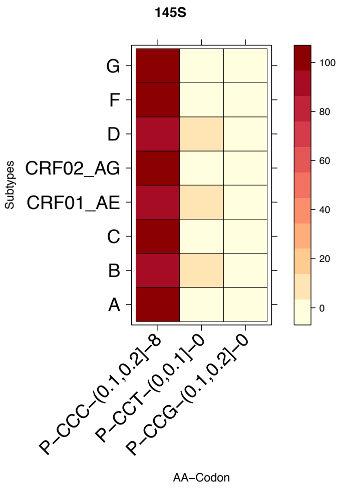
145S
G
F
D
CRF02_AG
CRF01_AE
C
B
A
Subtypes
100
80
60
40
20
0
P–CCC–(0.1,0.2]–8
P–CCT–(0,0.1]–0
P–CCG–(0.1,0.2]–0
AA–Codon

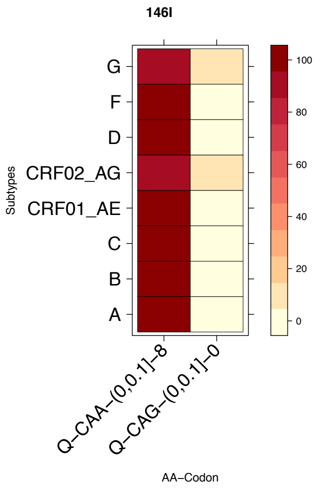
146I
G
F
D
CRF02_AG
CRF01_AE
C
B
A
Subtypes
100
80
60
40
20
0
Q–CAA–(0,0.1]–8
Q–CAG–(0,0.1]–0
AA–Codon

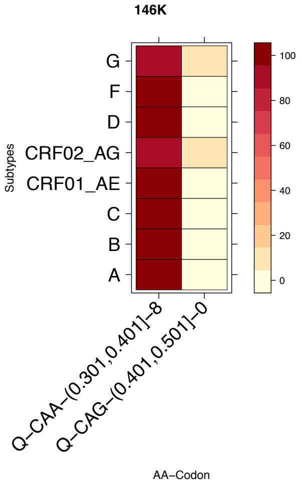
146K
Subtypes
G
F
D
CRF02_AG
CRF01_AE
C
B
A
Q–CAA–(0.301,0.401]–8
Q–CAG–(0.401,0.501]–0
AA–Codon
100
80
60
40
20
0

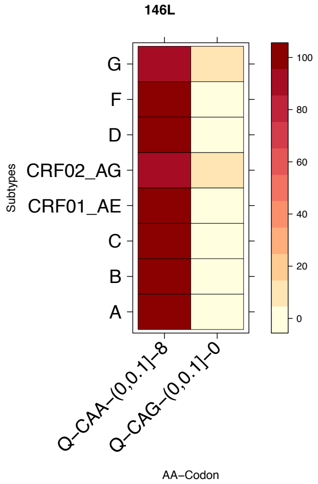
146L
Subtypes
G
F
D
CRF02_AG
CRF01_AE
C
B
A
Q–CAA–(0,0.1]–8
Q–CAG–(0,0.1]–0
AA–Codon
100
80
60
40
20
0

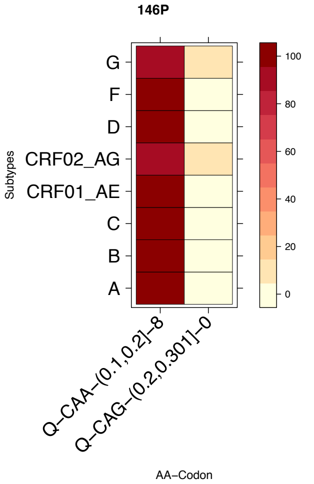
146P
Subtypes
G
F
D
CRF02_AG
CRF01_AE
C
B
A
Q–CAA–(0.1,0.2]–8
Q–CAG–(0.2,0.301]–0
AA–Codon
100
80
60
40
20
0

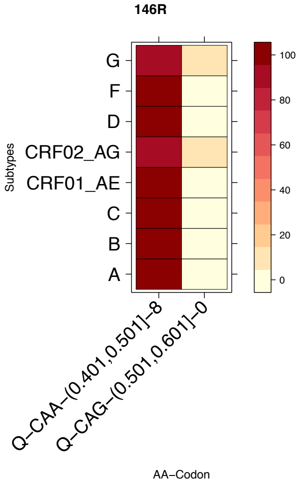
146R
Subtypes
G
F
D
CRF02_AG
CRF01_AE
C
B
A
Q–CAA–(0.401,0.501]–8
Q–CAG–(0.501,0.601]–0
AA–Codon
100
80
60
40
20
0

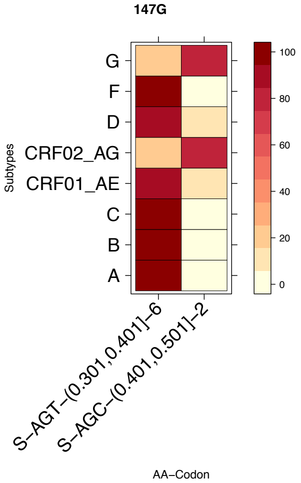
147G
Subtypes
G
F
D
CRF02_AG
CRF01_AE
C
B
A
S–AGT–(0.301,0.401]–6
S–AGC–(0.401,0.501]–2
AA–Codon
100
80
60
40
20
0

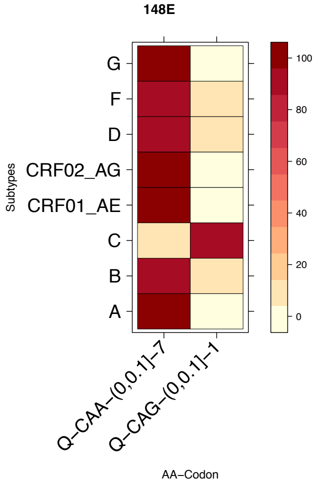
148E
Subtypes
G
F
D
CRF02_AG
CRF01_AE
C
B
A
Q–CAA–(0,0.1]–7
Q–CAG–(0,0.1]–1
AA–Codon
100
80
60
40
20
0

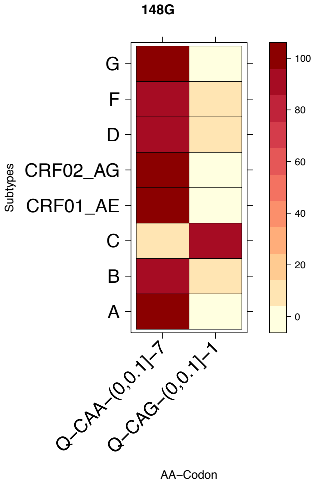
148G
G
F
D
CRF02_AG
CRF01_AE
C
B
A
Subtypes
Q−CAA−(0,0.1]−7
Q−CAG−(0,0.1]−1
AA−Codon
100
80
60
40
20
0

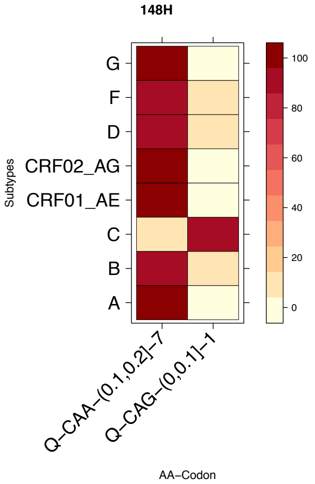
148H
G
F
D
CRF02_AG
CRF01_AE
C
B
A
Subtypes
Q−CAA−(0.1,0.2]−7
Q−CAG−(0,0.1]−1
AA−Codon
100
80
60
40
20
0

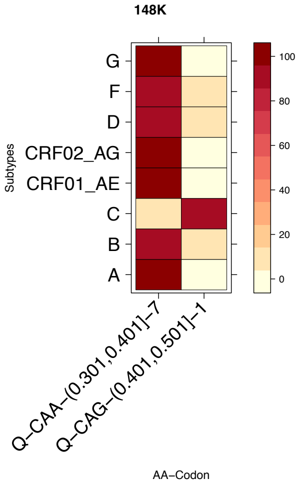
148K
G
F
D
CRF02_AG
CRF01_AE
C
B
A
Subtypes
Q−CAA−(0.301,0.401]−7
Q−CAG−(0.401,0.501]−1
AA−Codon
100
80
60
40
20
0

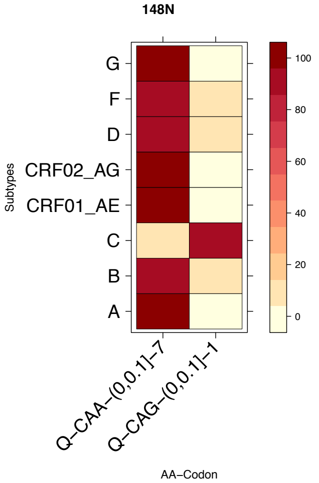
148N
G
F
D
CRF02_AG
CRF01_AE
C
B
A
Subtypes
Q−CAA−(0,0.1]−7
Q−CAG−(0,0.1]−1
AA−Codon
100
80
60
40
20
0

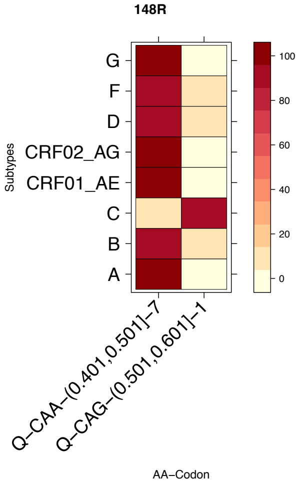
148R
G
F
D
CRF02_AG
CRF01_AE
C
B
A
Subtypes
Q−CAA−(0.401,0.501]−7
Q−CAG−(0.501,0.601]−1
AA−Codon
100
80
60
40
20
0

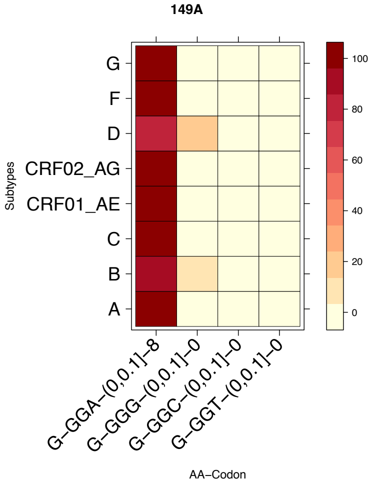
149A
G
F
D
CRF02_AG
CRF01_AE
C
B
A
Subtypes
G−GGA−(0,0.1]−8
G−GGG−(0,0.1]−0
G−GGC−(0,0.1]−0
G−GGT−(0,0.1]−0
AA−Codon
100
80
60
40
20
0

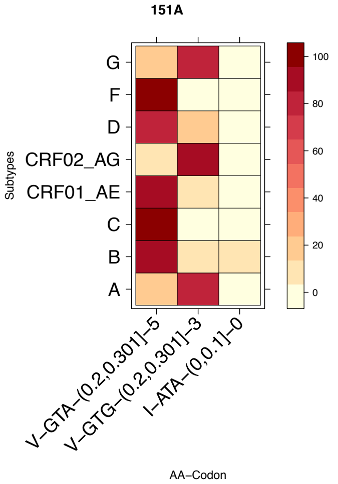
151A
G
F
D
CRF02_AG
CRF01_AE
C
B
A
Subtypes
V–GTA–(0.2,0.301]–5
V–GTG–(0.2,0.301]–3
I–ATA–(0,0.1]–0
AA–Codon
100
80
60
40
20
0

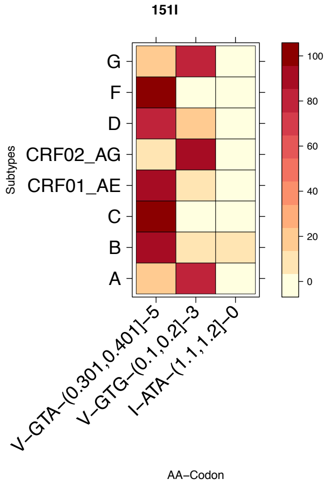
151I
G
F
D
CRF02_AG
CRF01_AE
C
B
A
Subtypes
V–GTA–(0.301,0.401]–5
V–GTG–(0.1,0.2]–3
I–ATA–(1.1,1.2]–0
AA–Codon
100
80
60
40
20
0

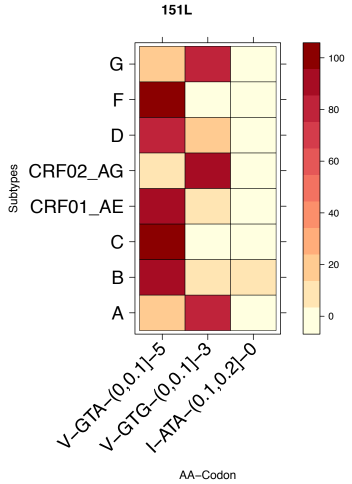
151L
G
F
D
CRF02_AG
CRF01_AE
C
B
A
Subtypes
V–GTA–(0,0.1]–5
V–GTG–(0,0.1]–3
I–ATA–(0.1,0.2]–0
AA–Codon
100
80
60
40
20
0

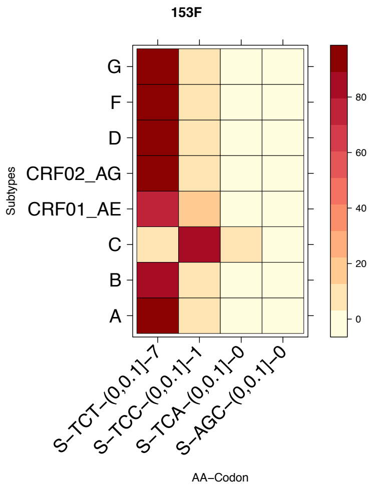
153F
G
F
D
CRF02_AG
CRF01_AE
C
B
A
Subtypes
S–TCT–(0,0.1]–7
S–TCC–(0,0.1]–1
S–TCA–(0,0.1]–0
S–AGC–(0,0.1]–0
AA–Codon
80
60
40
20
0

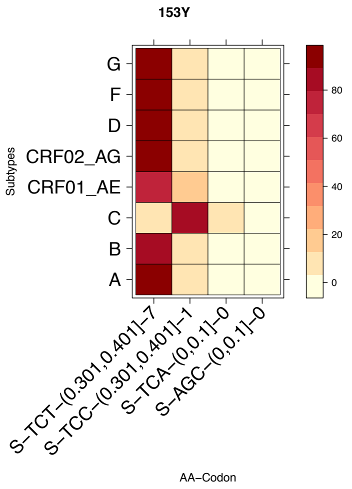
153Y
G
F
D
CRF02_AG
CRF01_AE
C
B
A
Subtypes
S–TCT–(0.301,0.401]–7
S–TCC–(0.301,0.401]–1
S–TCA–(0,0.1]–0
S–AGC–(0,0.1]–0
AA–Codon
80
60
40
20
0

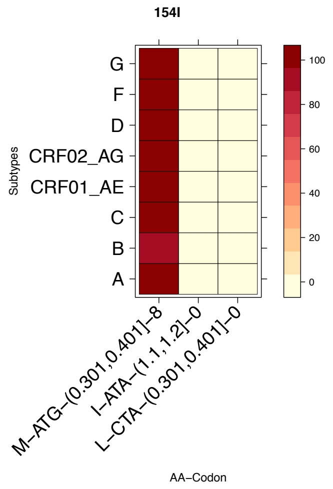
154I
G
F
D
CRF02_AG
CRF01_AE
C
B
A
Subtypes
M–ATG–(0.301,0.401]–8
I–ATA–(1.1,1.2]–0
L–CTA–(0.301,0.401]–0
AA–Codon
100
80
60
40
20
0

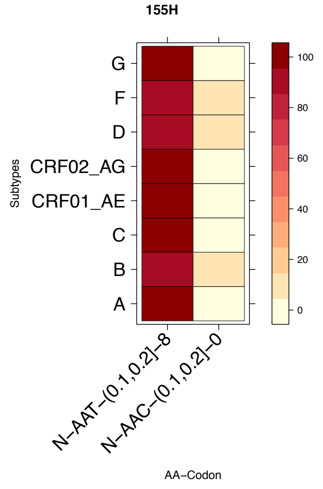
155H
G
F
D
CRF02_AG
CRF01_AE
C
B
A
Subtypes
N–AAT–(0.1,0.2]–8
N–AAC–(0.1,0.2]–0
AA–Codon
100
80
60
40
20
0

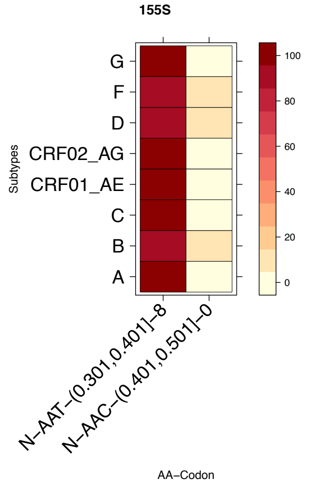
155S
G
F
D
CRF02_AG
CRF01_AE
C
B
A
Subtypes
N–AAT–(0.301,0.401]–8
N–AAC–(0.401,0.501]–0
AA–Codon
100
80
60
40
20
0

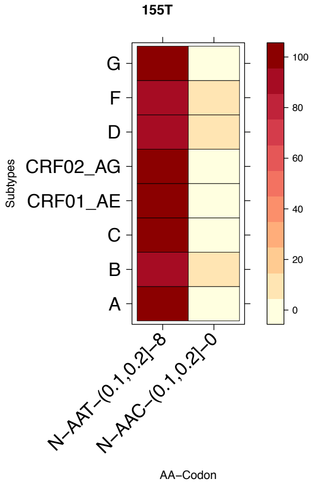
155T
G
F
D
CRF02_AG
CRF01_AE
C
B
A
Subtypes
N–AAT–(0.1,0.2]–8
N–AAC–(0.1,0.2]–0
AA–Codon
100
80
60
40
20
0

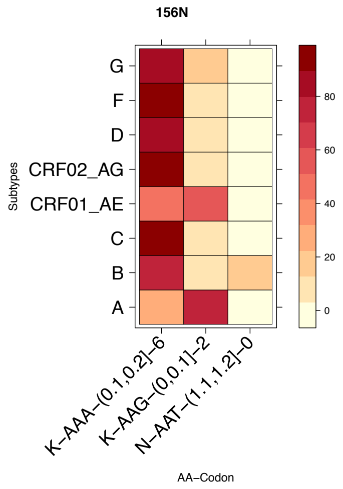
156N
G
F
D
CRF02_AG
CRF01_AE
C
B
A
Subtypes
K–AAA–(0.1,0.2]–6
K–AAG–(0,0.1]–2
N–AAT–(1.1,1.2]–0
AA–Codon
80
60
40
20
0

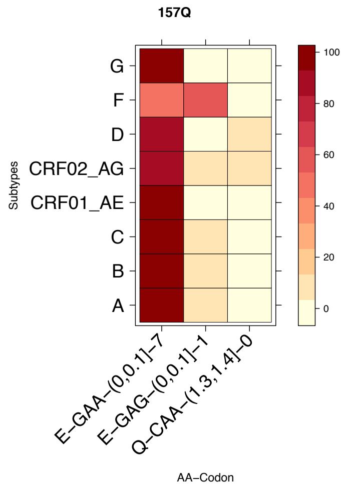
157Q
G
F
D
CRF02_AG
CRF01_AE
C
B
A
Subtypes
E–GAA–(0,0.1]–7
E–GAG–(0,0.1]–1
Q–CAA–(1.3,1.4]–0
AA–Codon
100
80
60
40
20
0

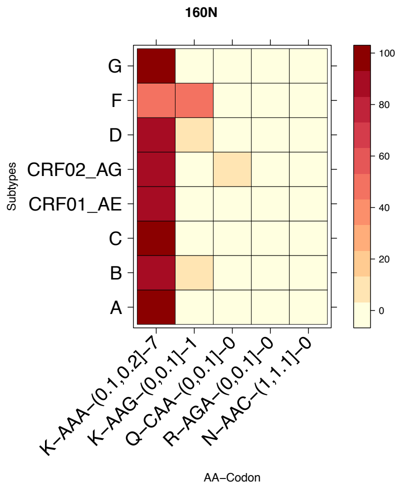
160N
G
F
D
CRF02_AG
CRF01_AE
C
B
A
Subtypes
K–AAA–(0.1,0.2]–7
K–AAG–(0,0.1]–1
Q–CAA–(0,0.1]–0
R–AGA–(0,0.1]–0
N–AAC–(1,1.1]–0
AA–Codon
100
80
60
40
20
0

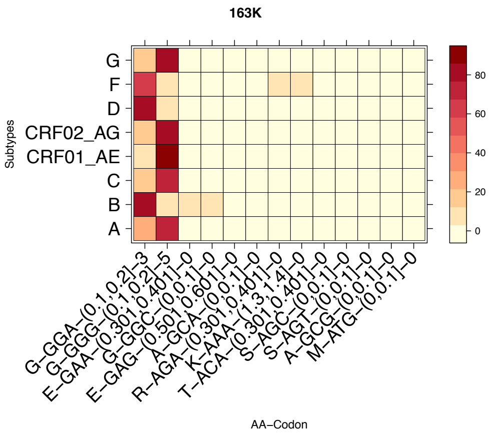
163K
Subtypes
G
F
D
CRF02_AG
CRF01_AE
C
B
A
G–GGA–(0.1,0.2]–3
G–GGG–(0.1,0.2]–5
E–GAA–(0.301,0.401]–0
G–GGC–(0,0.1]–0
E–GAG–(0.501,0.601]–0
A–GCA–(0,0.1]–0
R–AGA–(0.301,0.401]–0
K–AAA–(1.3,1.4]–0
T–ACA–(0.301,0.401]–0
S–AGC–(0,0.1]–0
S–AGT–(0,0.1]–0
A–GCG–(0,0.1]–0
M–ATG–(0,0.1]–0
AA–Codon
80
60
40
20
0

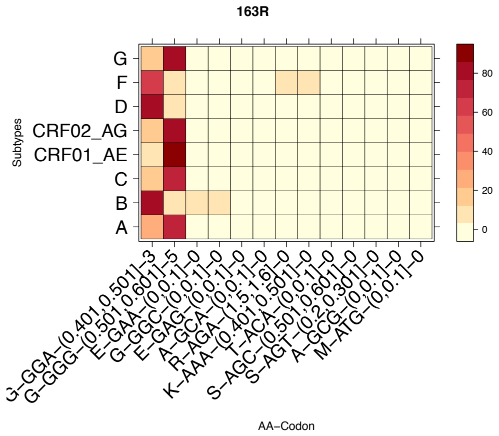
163R
Subtypes
G
F
D
CRF02_AG
CRF01_AE
C
B
A
G–GGA–(0.401,0.501]–3
G–GGG–(0.501,0.601]–5
E–GAA–(0,0.1]–0
G–GGC–(0,0.1]–0
E–GAG–(0,0.1]–0
A–GCA–(0,0.1]–0
R–AGA–(1.5,1.6]–0
K–AAA–(0.401,0.501]–0
T–ACA–(0,0.1]–0
S–AGC–(0.501,0.601]–0
S–AGT–(0.2,0.301]–0
A–GCG–(0,0.1]–0
M–ATG–(0,0.1]–0
AA–Codon
80
60
40
20
0

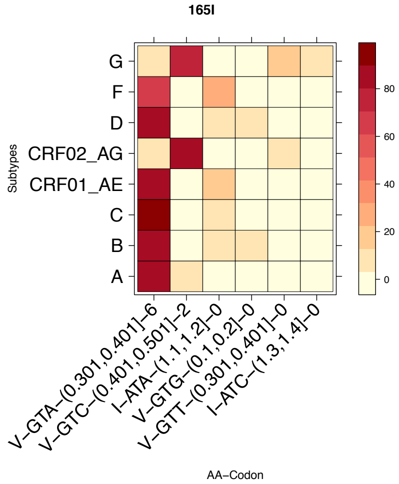
165I
Subtypes
G
F
D
CRF02_AG
CRF01_AE
C
B
A
V–GTA–(0.301,0.401]–6
V–GTC–(0.401,0.501]–2
I–ATA–(1.1,1.2]–0
V–GTG–(0.1,0.2]–0
V–GTT–(0.301,0.401]–0
I–ATC–(1.3,1.4]–0
AA–Codon
80
60
40
20
0

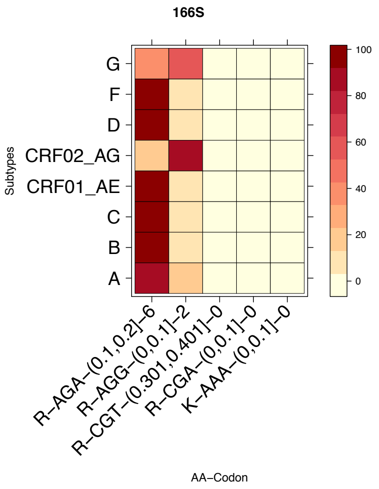
166S
Subtypes
G
F
D
CRF02_AG
CRF01_AE
C
B
A
R–AGA–(0.1,0.2]–6
R–AGG–(0,0.1]–2
R–CGT–(0.301,0.401]–0
R–CGA–(0,0.1]–0
K–AAA–(0,0.1]–0
AA–Codon
100
80
60
40
20
0

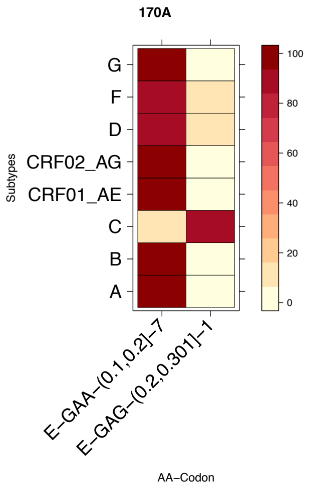
170A
Subtypes
G
F
D
CRF02_AG
CRF01_AE
C
B
A
E–GAA–(0.1,0.2]–7
E–GAG–(0.2,0.301]–1
AA–Codon
100
80
60
40
20
0

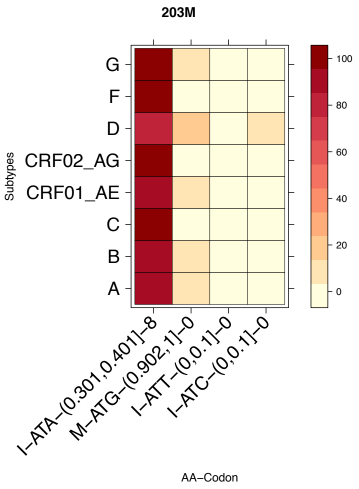
203M
Subtypes
G
F
D
CRF02_AG
CRF01_AE
C
B
A
I–ATA–(0.301,0.401]–8
M–ATG–(0.902,1]–0
I–ATT–(0,0.1]–0
I–ATC–(0,0.1]–0
AA–Codon
100
80
60
40
20
0

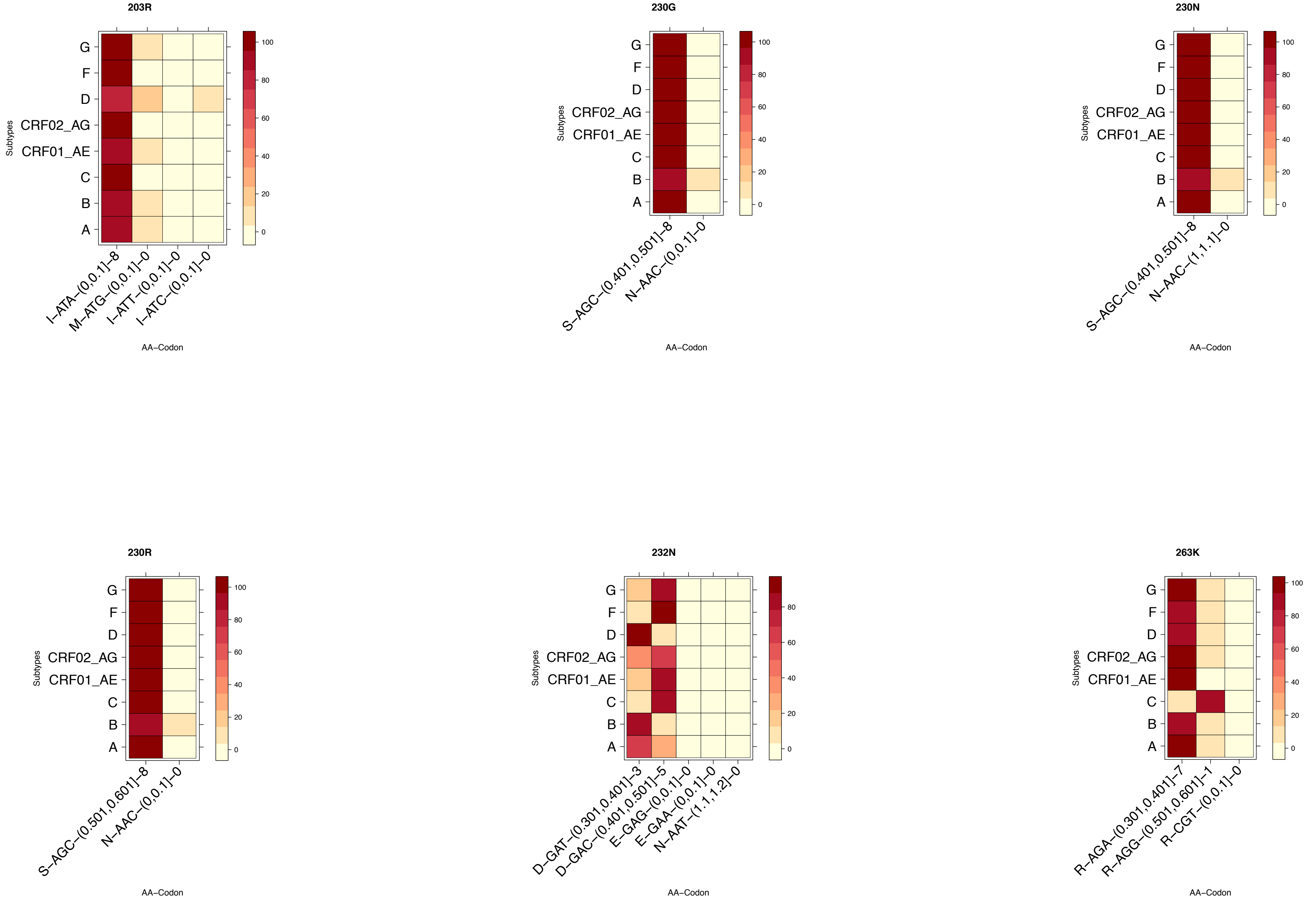
203R
G
F
D
CRF02_AG
CRF01_AE
C
B
A
Subtypes
I−ATA−(0,0.1]−8
M−ATG−(0,0.1]−0
I−ATT−(0,0.1]−0
I−ATC−(0,0.1]−0
AA−Codon
100
80
60
40
20
0
230G
S−AGC−(0.401,0.501]−8
N−AAC−(0,0.1]−0
230N
S−AGC−(0.401,0.501]−8
N−AAC−(1,1.1]−0
230R
S−AGC−(0.501,0.601]−8
N−AAC−(0,0.1]−0
232N
D−GAT−(0.301,0.401]−3
D−GAC−(0.401,0.501]−5
E−GAG−(0,0.1]−0
E−GAA−(0,0.1]−0
N−AAT−(1.1,1.2]−0
263K
R−AGA−(0.301,0.401]−7
R−AGG−(0.501,0.601]−1
R−CGT−(0,0.1]−0
